# Define and visualize pathological architectures of human tissues from spatially resolved transcriptomics using deep learning

**DOI:** 10.1101/2021.07.08.451210

**Authors:** Yuzhou Chang, Fei He, Juexin Wang, Shuo Chen, Jingyi Li, Jixin Liu, Yang Yu, Li Su, Anjun Ma, Carter Allen, Yu Lin, Shaoli Sun, Bingqiang Liu, Jose Otero, Dongjun Chung, Hongjun Fu, Zihai Li, Dong Xu, Qin Ma

## Abstract

Spatially resolved transcriptomics provides a new way to define spatial contexts and understand biological functions in complex diseases. Although some computational frameworks can characterize spatial context via various clustering methods, the detailed spatial architectures and functional zonation often cannot be revealed and localized due to the limited capacities of associating spatial information. We present RESEPT, a deep-learning framework for characterizing and visualizing tissue architecture from spatially resolved transcriptomics. Given inputs as gene expression or RNA velocity, RESEPT learns a three-dimensional embedding with a spatial retained graph neural network from the spatial transcriptomics. The embedding is then visualized by mapping as color channels in an RGB image and segmented with a supervised convolutional neural network model. Based on a benchmark of sixteen 10x Genomics Visium spatial transcriptomics datasets on the human cortex, RESEPT infers and visualizes the tissue architecture accurately. It is noteworthy that, for the in-house AD samples, RESEPT can localize cortex layers and cell types based on a pre-defined region-or cell-type-specific genes and furthermore provide critical insights into the identification of amyloid-beta plaques in Alzheimer’s disease. Interestingly, in a glioblastoma sample analysis, RESEPT distinguishes tumor-enriched, non-tumor, and regions of neuropil with infiltrating tumor cells in support of clinical and prognostic cancer applications.

## Introduction

Tissue architecture is the biological foundation of spatial heterogeneity within complex organs like the human brain^1^ and is thereby essential in understanding the underlying pathogenesis of human diseases, including cancer^2^ and Alzheimer’s disease (AD)^3^. Recent advances in spatially resolved technologies such as 10x Genomics Visium provide spatial context together with high-throughput gene expression for exploring tissue domains, cell types, cell-cell communications, and their biological consequences^4^. Some graph-based clustering methods^5,6^, statistical methods^7^, or deep learning-based methods^8,9^ can identify spatial architecture and interpret spatial heterogeneity. For example, Seurat^10^ and Giotto^11^ use a similar framework on variable gene selection, dimension reduction, followed by graph-based clustering (i.e., Louvain). STUtility^12^ uses non-negative matrix factorization to perform dimension reduction and then identifies tissue architecture based on the Seurat framework. SpaGCN^9^ proposes a convolutional graph network to integrate gene expression, spatial location, and histology in spatial transcriptomics data analysis. stLearn^8^ also integrates gene expression, spatial location, and histology information in the normalization method and applies the Louvain algorithm as a clustering method. Another approach, BayesSpace^7^ adopts a Bayesian statistical framework to adjust spatial neighborhoods for resolution enhancement and for clustering analysis. Even existing methods can provide some useful information, the intrinsic tissue architecture, however, often cannot be fully revealed due to a lack of strong spatial representation for the biological context in tissues, and these tools often do not take full advantage of spatial information and are limited in predicting tissue architectures. Therefore, it is still challenging to accurately characterize tissue architectures and the underlying biological functions from spatial transcriptomics.

We reasoned that spatial transcriptomics could be effectively represented and intuitively visualized as an image with expression abundance retaining the spatial context. To this end, we introduce **RESEPT** (***RE***constructing and ***S***egmenting ***E***xpression mapped RGB images based on s***P***atially resolved ***T***ranscriptomics), a framework for reconstructing, visualizing, and segmenting an RGB image from spatial transcriptomics to reveal tissue architecture and spatial heterogeneity. We highlight the unique features of RESEPT as follows: (*i*) Spatial transcriptomics data are converted as an RGB image by mapping a low dimensional embedding to color channels via a spatial retained graph neural network. This image represents various spatial contexts together with expression abundance faithfully, and it resists robustly to noises due to limitations of measuring technology. (*ii*) An RGB image is segmented to predict spatial cell types using a pre-trained segmentation deep-learning model and an optional segmentation quality assessment protocol. (*iii*) RNA velocity can be integrated into image training, which is effective in revealing some tissue architectures. (*iv*) With a defined panel of gene sets representing specific biological pathways or cell lineages, RESEPT can recognize the spatial pattern and detect the corresponding active functional regions. (*v*) The functional zonation boundaries of AD are determined effectively by the pre-trained image segmentation deep-learning model. (*vi*) RESEPT successfully recognized tumor architecture, non-tumor architecture, and infiltration tumor architecture in clinical and prognostic applications on glioblastoma.

## Results

### The architecture of RESEPT comprises representation learning and segmentation

Spatial transcriptomics data are represented as a spatial spot-spot graph by RESEPT (**Fig. 1**). Each observational unit within a tissue sample containing a small number of cells, *i.e*., “spot,” is modeled as a node. The measured gene expression values of the spot are treated as the node attributes, and the neighboring spots adjacent in the Euclidean space on the tissue slice are linked with an undirected edge. This lattice-like spot graph is modeled by our graph neural network (GNN) based tool scGNN^13^, which learns a three-dimensional embedding to preserve the topological relationship between all spots in the spatial space of transcriptomics. The three-dimensional embedding on gene expression is mapped to three color channels as Red, Green, and Blue in an RGB image, which is naturally visualized as an image of the spatial gene expression. Then a semantic segmentation can be performed on the image to identify the spatial architecture by classifying each spot into a spatially specific segment with a supervised convolutional neural network (CNN) model.

**Fig. 1.**
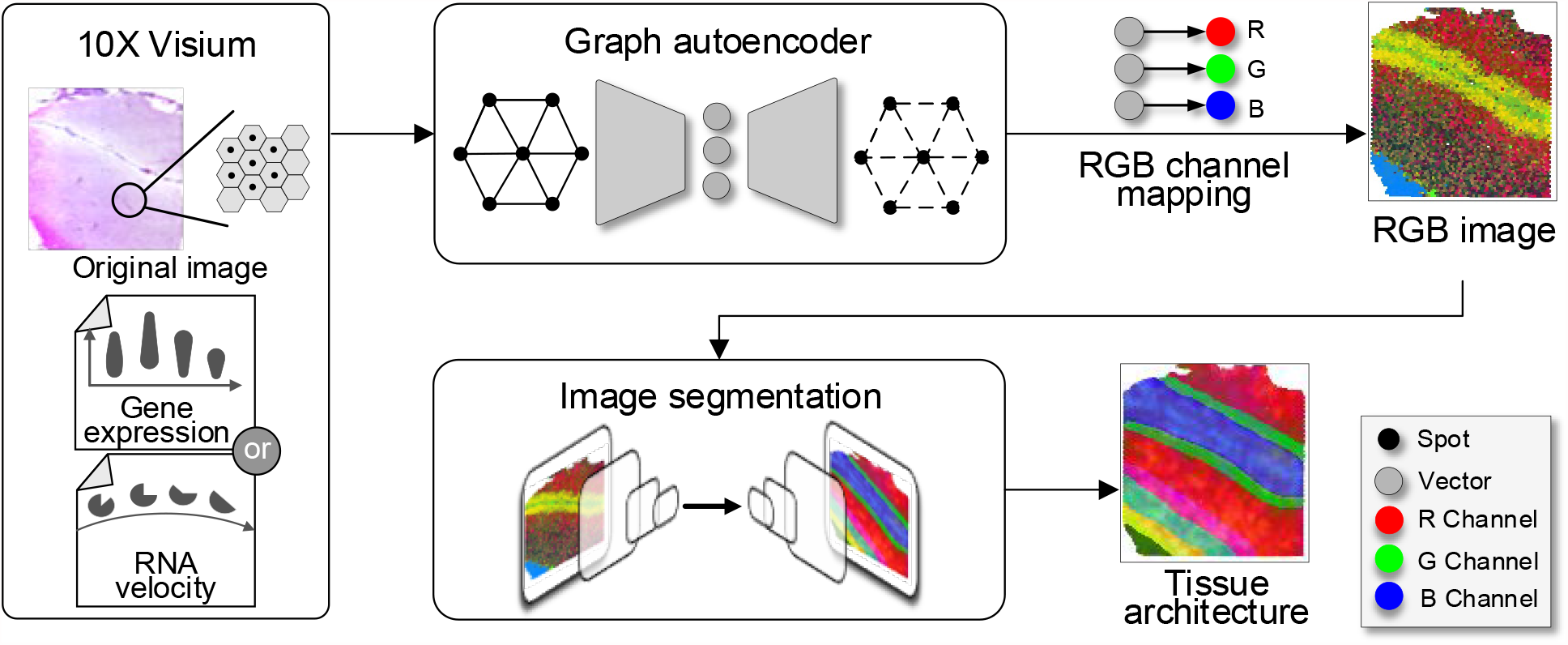
The RESEPT schema. RESEPT takes gene expression or RNA velocity from spatial transcriptomics as the input. The input is embedded into a three-dimensional representation by a spatially constrained Graph Autoencoder, then linearly mapped to an RGB color spectrum to reconstruct an RGB image. A CNN image segmentation model is trained to obtain a spatially specific architecture (from whole-gene embedding) or spatial functional regions (from panel-gene embedding).

In the 10x Visium Genome platform, each spot has six adjacent spots, so the spatial retained spot graph has a fixed node degree six for all the nodes. On the generated spatial spot-spot graph, a graph autoencoder learns a node-wise three-dimensional representation to preserve topological relations in the graph. The encoder of the graph autoencoder composes two layers of graph convolution network (GCN) to learn the 3-dimensional graph embedding. The decoder of the graph autoencoder is defined as an inner product between the graph embedding, followed by sigmoid activation function. The goal of graph autoencoder learning is to minimize the difference between the input and the reconstructed graph (**Fig. 2a**).

**Fig. 2.**
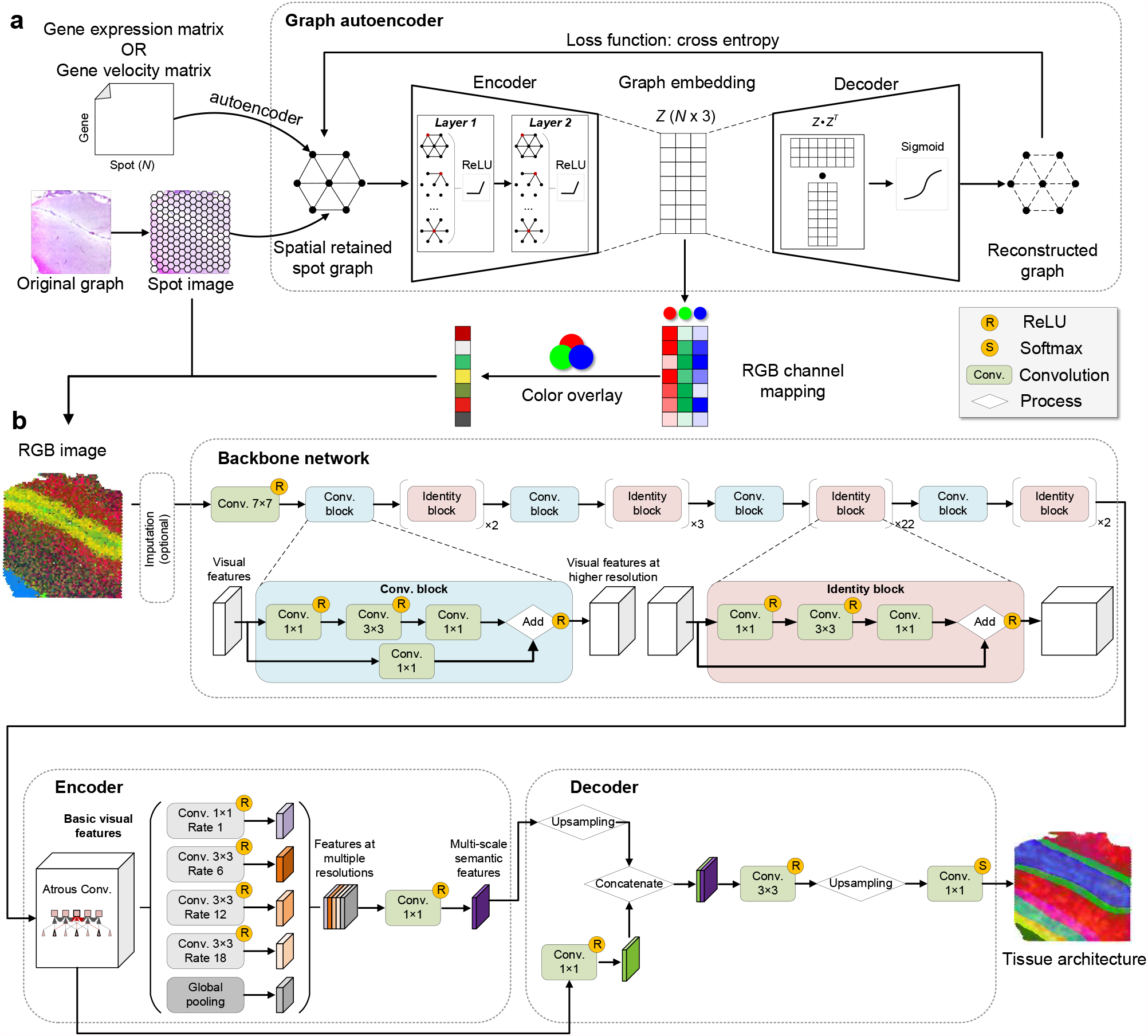
The RESEPT framework. (a) A spatial retained spot graph is established by spatial distances of spots and their expression or velocity matrix. The graph autoencoder takes the adjacent distance matrix of the spot graph as the input. Its encoder learns a 3-dimensional embedding of a spatial cell graph. The decoder reconstructs the adjacent correlations among all cells by dot products of the 3-dimensional embeddings followed by a sigmoid activation function. The graph autoencoder is trained by minimizing the cross-entropy loss between the input spatial and the reconstructed graphs. The learned 3-dimensional embeddings are mapped to a full-color spectrum to generate an RGB image revealing the spatial architecture. (b) The segmentation model takes the RGB image as the input, which may be processed with an imputation operation if missing spots exist. Its backbone network ResNet101 consists of one convolutional layer and a series of residual blocks, in which one type of residual block named convolutional block stacks 3 convolutional layers with a convolutional skip connection from the input signals to the output feature maps, and the other type of residual block identity block stacks 3 convolutional layers with a direct skip connection from the input signals to the output feature maps. This extra deep network firstly extracts rich visual features of the input image. The encoder module further extracts multi-scale semantic features by applying four atrous convolutional with different rates and sizes of filters and one global pooling layer respectively to the basic visual feature maps. And the decoder module up-samples the multi-scale features to the same size with basic visual feature maps and then concatenates them together. After a softmax activation function, the decoder module outputs a segmentation map classifying each spot into a specific spatial architecture.

The segmentation architecture is comprised of a backbone network, an encoder module, and a decoder module. The backbone network employs an extra deep network ResNet101^14^ to provide basic visual features of the input RGB image. ResNet101 stacks one convolutional layer and 33 residual blocks, each of which cascades 3 convolutional layers with a convolutional skip connection from the input signals to the output feature maps, for extracting sufficiently rich features. The encoder module utilizes atrous convolutional layers with various rates and sizes of filters and one global pooling layer respectively to detect multi-scale semantic features from ResNet101 feature maps. And the decoder module aligns the multi-scale features to the same size and outputs a segmentation map classifying each spot into a specific spatial architecture. (**Fig. 2b**)

### RESEPT accurately characterizes the spatial architecture of the human brain cortex region

Using manual annotations as the ground truth on 12 published samples^15^ and four in-house samples^16^ sequenced on the 10x Genomics Visium platform, RESEPT was benchmarked on both raw and normalized expression matrices of the 16 samples (S2-S17 in **Fig. 3a** and **Table 1**). Our results demonstrate RESEPT outperforms six existing tools, namely Seurat^10^, BayesSpace^7^, SpaGCN^9^, stLearn^8^, STUtility^12^, and Giotto^11^ on tissue architecture identification in terms of Adjusted Rand Index (ARI) 0.706 ± 0.163 (**Fig. 3b**) based on tuned parameters (**Supplementary Data 1**). Additional benchmarking results in default parameter settings with different evaluation matrices, visualization of RESEPT outcome, running time, and memory usage can be referred to **Fig. 3c, Supplementary Fig. 1**, and **Supplementary Data 2-3**. To validate the stability of our model, we generated simulation data with gradient decreasing sequencing depth based on two selected datasets S5 and S6 (**Fig. 3a**). The RGB images at low read depth presented more intra-regional diversity in their color distributions (**Supplementary Fig. 2** and **Supplementary Data 4**). In the downsampling read depth gradients from very low depth to full depth, RESEPT demonstrated its robustness by ARI 0.454 ± 0.014 on S5, and ARI 0.809 ± 0.006 on S6 (**Fig. 3d-g**). It is noteworthy that RGB images generated from RNA velocity^17,18^ can reveal clear spatial separation between segments from the identified architecture on the AD sample S4 (Moran’s I 0.920 vs 0.787), which is consistent with the brain development zonation (**Fig. 3h**). On the same sample, RESEPT reveals better tissue architecture than the other tools in ARI 0.409 (**Fig. 3i**). More visualization results from different normalization methods can be referred to **Supplementary Fig. 1** and **Supplementary Data 5**. All the data used in the study are summarized in **Supplementary Table 1**, while datasets on 10x Genomics, Spatial Transcriptomics (ST), and High-Definition Spatial Transcriptomics (HDST) platforms without manual annotations were analyzed by RESEPT detailed in **Supplementary Fig. 3**.

**Table 1:** *Details of 10x Visium data used in the study*. The table lists 17 samples of information. The Sample column indicates the sample number in this study. The Protocol column indicates the revision number of two Visium protocols. The Tissue column indicates the sample disease’s status. The # of spot column indicates the number of spots. The Mean reads column indicates the number of reads for each spot from the bam file. The Median gene column indicates the median number of detected genes for each spot. The Total reads column indicates the total number of each sample calculated from the expression matrix. The Mean reads column indicates the mean read of each spot from the expression matrix. The SD reads column indicates the standard deviation of each spot calculated from the expression matrix. Abbreviations: Alzheimer’s disease (AD), CG000239 -Visium Spatial Gene Expression Reagent Kits-User Guide Rev D, Oct.2020 (Rev D), CG000239 -Visium Spatial Gene Expression Reagent Kits-User Guide Rev A, Nov. 2019 (Rev A).

| Sample | Protocol | Tissue | Web reported |  |  | Expression matrix |  |  |
| --- | --- | --- | --- | --- | --- | --- | --- | --- |
|  |  |  | #spot | Mean reads | Median gene | Total reads | Mean reads | SD reads |
| S1 | Rev D | Tumor | 3,468 | 11,596 | 4,326 | 43,841,318 | 12,641.670 | 7,204.035 |
| S2 | Rev D | Health brain | 4,701 | 42,484 | 3,022 | 43,225,942 | 9,195.053 | 8,771.039 |
| S3 | Rev D | AD | 3,445 | 36,569 | 3,722 | 30,383,719 | 8,819.657 | 4,275.528 |
| S4 | Rev D | AD | 4,832 | 33,660 | 3,664 | 41,180,024 | 8,522.356 | 4,789.882 |
| S5 | Rev D | health brain | 4,225 | 43,186 | 3,458 | 33,815,249 | 8,003.609 | 3,527.456 |
| S6 | Rev A | health brain | 3,672 | 223,921 | 2,610 | 21,699,243 | 5,907.771 | 2,848.429 |
| S7 | Rev A | Health brain | 3,641 | 82,583 | 2,113 | 16,701,265 | 4,589.520 | 2,356.537 |
| S8 | Rev A | Health brain | 4,111 | 118,826 | 1,854 | 16,042,438 | 3,903.270 | 1,955.168 |
| S9 | Rev A | Health brain | 3,459 | 92,729 | 1,813 | 13,391,960 | 3,870.509 | 1,931.687 |
| S10 | Rev A | Health brain | 4,634 | 58,483 | 1,344 | 13,823,583 | 3,775.904 | 1,920.972 |
| S11 | Rev A | Health brain | 4,021 | 69,839 | 1,742 | 14,590,115 | 3,633.902 | 1,789.643 |
| S12 | Rev A | Health brain | 3,592 | 65,000 | 1,695 | 12,923,757 | 3,597.928 | 1,988.986 |
| S13 | Rev A | Health brain | 3,499 | 65,523 | 1,607 | 12,007,005 | 3,432.534 | 1,772.158 |
| S14 | Rev A | Health brain | 4,226 | 76,928 | 1,384 | 10,955,668 | 2,592.444 | 1,198.877 |
| S15 | Rev A | Health brain | 4,787 | 58,813 | 1,407 | 12,243,054 | 2,556.495 | 1,250.268 |
| S16 | Rev A | Health brain | 3,662 | 91,654 | 1,736 | 11,356,262 | 2,450.639 | 1,150.815 |
| S17 | Rev A | Health brain | 4,383 | 60,244 | 1,159 | 9,325,211 | 2,127.101 | 1,046.434 |

**Fig. 3.**
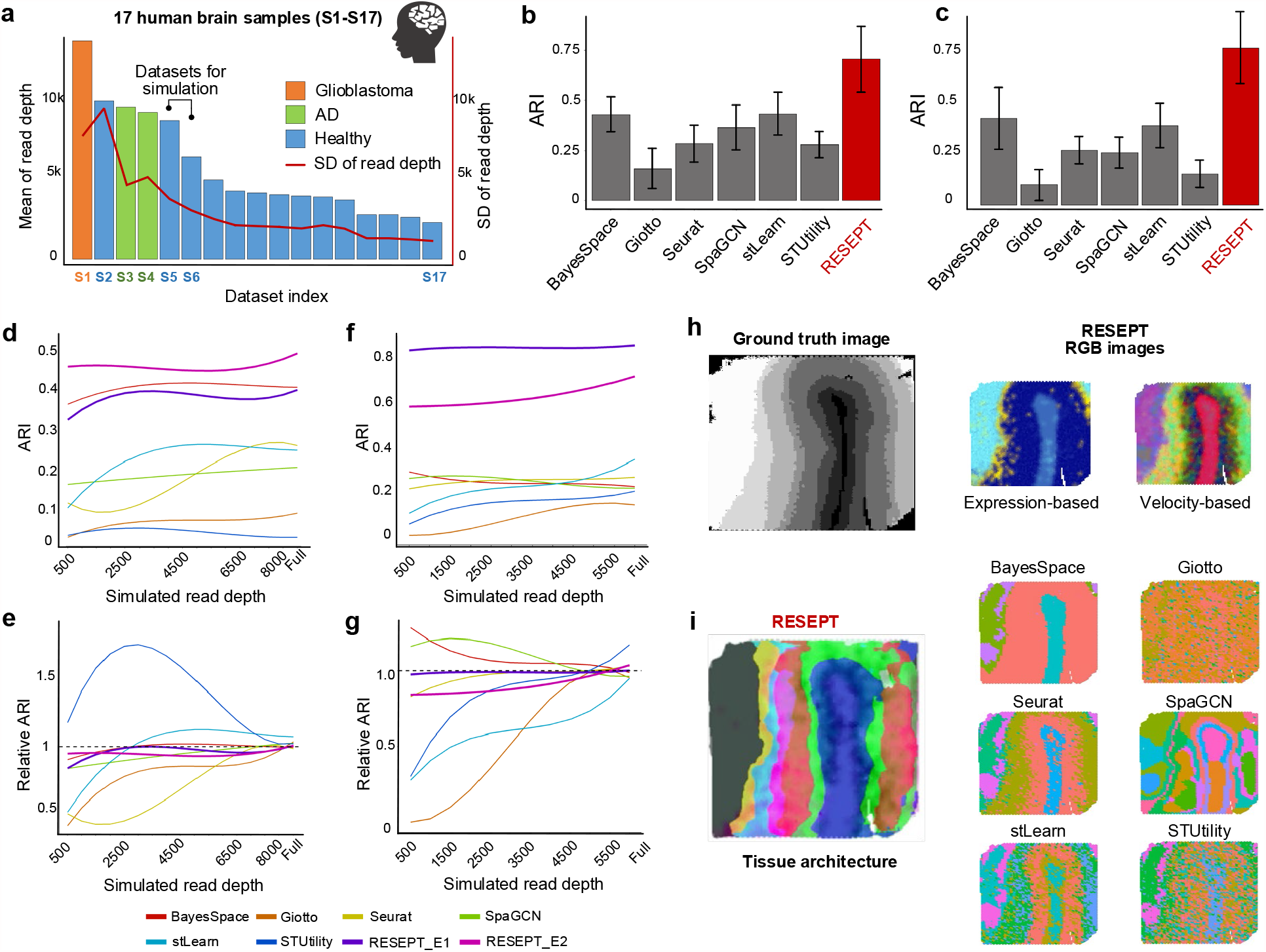
The RESEPT workflow and performance. (**a**) show mean and standard deviation of sequencing reads of 17 human brain datasets on 10x Visium platform. S2-S17 have manual annotations as the benchmark, S5 & S6 for simulation for high mean and low standard deviations of read depth, S1 & S4 for the case studies (more details in Supplementary Tables 1-2). (**b**) Performance of tissue architecture (with 7 clusters pre-defined) identification by six existing tools and RESEPT on criteria ARI. (**c**) Performance of tissue architecture (default parameters) identification by six existing tools and RESEPT on criteria ARI. (**d**) Stability of tissue architecture identification across sequencing depths on samples S5 using different tools. The Y-axis shows ARI performance, and the X-axis represents the sequencing depth with subsampling. The lines are smoothed by the B-Spline smooth method. (**e**) Normalized performance vs. sequencing depth on sample S5. Performance of full sequencing depth is set as 1.0. RESEPT_E1 using scGNN embedding, RESEPT_E2 using spaGCN embedding. (**f**) and (**g**) show the stability of ARI and normalized performance against grid sequencing depth for sample S6. (**h**) RGB image generated from RNA velocity reveals better architecture (Moran’s I = 0.920) than gene expression (Moran’s I = 0.787) on the AD sample S4. (**i**) Spatial domains on S4 detected by RESEPT, together with those identified by other tools.

RESEPT benefits from the representation power of the learned embedding from the spatially constrained GNN comparing with spaGCN and UMAP (**Supplementary Figs. 4-5**). The sufficiently diverse training images (**Supplementary Fig. 6**) and fine-gained visual features extracted from the extra deep CNN network also give strong discerning power to our segmentation model. We also validated the performance improvement with an increasing number of annotated training data (**Supplementary Fig. 7**). This improvement implied that as more annotated spatial transcriptomic data comes out, RESEPT will enhance its robustness accordingly.

**Fig. 4.**
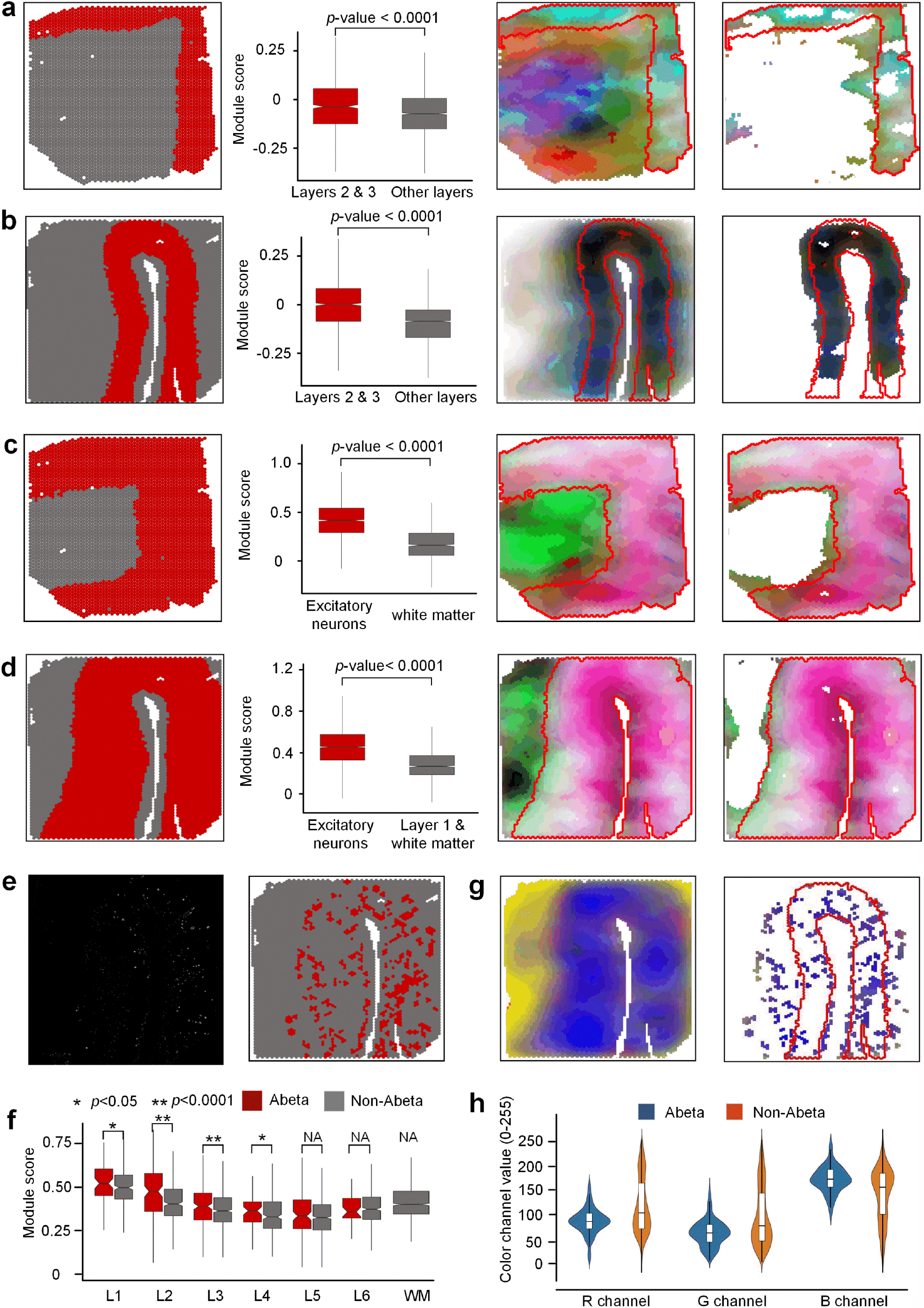
RESEPT identifies spatial cellular patterns in the human postmortem middle temporal gyrus (MTG). (**a**) The box plot shows the module score of the cortical layers 2 and 3 and other layers from Sample S3, where the x-axis shows layer categories and the y-axis represents scores. The second figure shows layer 2 and 3 architecture (red); the third figure shows an RGB image; the fourth figure is reconstructed by filtering out unrelated colors. (**b**) The box plot shows the module score of excitatory neurons from layers 2 to 6 and other layers. The second figure shows the ground truth of layers 2 to 6; the third figure shows an RGB image; the fourth figure is reconstructed by filtering out unrelated colors. (c) and (d) display the same layer architecture and cell type localization for sample S4. (**e**) The left figure was generated by immunofluorescence assay to show Aβ plaques location, and the right figure highlights the spots with the accumulation of Aβ plaques. (**f**) The box plot shows scores for the Aβ region and the non-Aβ region split by six layers and white matter. (**g**) The left figure shows the RGB image from the 20 genes embedding results, and the right figure shows the RGB image cropped according to the Aβ region and marked by layers 2&3 (encircled by the red line). (**h**) RGB channel shows the color value dispersion, where blue represents RGB values in the Aβ region and orange represents RGB values in the non-Aβ region.

### RESEPT interprets and discovers spatially related biological insights in AD

With our in-house AD brain samples^16^, human postmortem middle temporal gyrus (MTG) from an AD case (Sample S4) was spatially profiled on the 10x Visium platform, and RESEPT successfully identified the main architecture of the MTG comparing with the manual annotation as the ground truth (S3 ARI = 0.474; S4 ARI=0.409). With the RGB image generated from specific gene expression, we distinguish cortical layers 2 & 3 from other layers and identified regions enriched with excitatory neurons and amyloid-beta (Aβ) plaques. For the AD sample on cortical layers 2 & 3 (ground truth^16^ as **Fig. 4a-b**), well-defined marker genes (C1QL2, RASGRF2, CARTPT, WFS1, HPCAL1 for layer 2, and CARTPT, MFGE8, PRSS12, SV2C, HPCAL1 for layer 3) from the previous study^19^ were embedded and transformed to an RGB image instead of using whole transcriptomes (a full gene list in **Supplementary Table 2**). To validate the spatial specificity, module scores from Seurat^10^ showed that these marker genes are statistically significantly enriched only on cortex layers 2 & 3 among all the layers (*p*<0.0001 by Wilcoxon signed-rank test). Furthermore, RESEPT visually provided consistent colors for cortical layers 2 & 3. These spatial patterns were strengthened by filtering unrelated colors. More RGB images from other layer-specific marker genes can be found in **Supplementary Fig. 8**. To reveal critical cell-type distribution (*i.e*., <u>excitatory neuron</u>) associated with selective neuronal vulnerability in AD^20^, five well-defined excitatory neuron marker genes (SLC17A6, SLC17A7, NRGN, CAMK2A, and SATB2) in the cortex were obtained from our in-house database scREAD^21^ (other cell-type marker genes in **Supplementary Table 2**). The module score and optimized RGB image (**Fig. 4c-d**) showed statistically significant enrichment of excitatory neuron marker genes in cortical layers 2-6 (*p*<0.0001 by Wilcoxon signed-rank test), and the original and improved RGB image also localized the excitatory neurons (other cell types can be found in **Supplementary Fig. 9**). Moreover, the RGB image can reflect an important AD pathology-associated region, i.e., Aβ plaques-accumulated region. We conducted an immunofluorescence staining of Aβ on the adjacent AD brain section (see details in **Methods**) and identified the brain region with Aβ plaques^16^ (**Fig. 4e**). Among the gene module containing 57 Aβ plaque-induced genes discovered from the previous study^2^, we validated those 20 upregulated genes showed the specific enrichment in the Aβ region compared to the non-Aβ region in terms of layers 2 & 3 (*p*<0.0001 by Wilcoxon signed-rank test, **Fig. 4f**). By comparing the color in Aβ region-associated spots with the RGB image (**Fig. 4g**), we observed Aβ region-associated spots behaved a consistent color in layers 2 & 3. To evaluate RGB value variation quantitatively, we investigated the value range of channels R, G, and B for the Aβ region and non-Aβ region (**Fig. 4h**). The result showed that the Aβ region had a tight dispersion compared to the non-Aβ region, which proved the RGB image can be potentially used to indicate the pathological regions with Aβ plaques. Overall, with the evidence of images generated from hallmark panel genes, RESEPT can confidently reflect layer-specific, cell-type-specific, and pathological region-specific architecture, with well-studied marker genes and disease-associated genes. These results indicate significant potentials and strong applicative power of RESEPT to localize and present important spatial architecture contributing to AD pathology.

### The clinical and prognostic applications of RESEPT in cancer

To demonstrate the clinical and prognostic applications of RESEPT in the oncology field, we analyzed a glioblastoma dataset published by 10x Genomics using the Visium platform (**Fig. 5a**, Sample S1). Glioblastoma, a grade IV astrocytic tumor with a median overall survival of 15 months^22^, is characterized by heterogeneity in tissue morphologies which range from highly dense tumor cellularity with necrosis to other areas with single tumor cell permeation throughout the neuropil. Assessment of tissue architecture represents a key diagnostic tool for patient prognosis and diagnosis. RESEPT identified eight segments (**Fig. 5b**-**c, Supplementary Fig. 10**) and distinguished tumor-enriched, non-tumor, and regions of neuropil with infiltrating glioblastoma cells. These segmented areas show similarities to secondary structures of Scherer^23^. Based on the morphological features of Segment 3 in the Hematoxylin-Eosin (H&E) image (**Fig. 5c**), we observed cells with large cytoplasm and nuclei with prominent nucleoli, a morphology consistent with cortical pyramidal neurons, and many tumor cells located in this segment showing neuronal satellitosis. Differentially expressed gene (DEG) analysis demonstrated that a pre-defined glioblastoma marker CHI3L1^24,25^ was highly expressed in most of the spots in Segment 3 (**Fig. 5d, differentially expressed gene of each segment can be found Supplementary Data 6**). By exploring the H&E image of Segment 6, we found this prominent area of the segment with erythrocytes, likely representing an area of acute hemorrhage during the surgical biopsy. This morphological observation was in line with the GO enrichment analysis, where DEGs were enriched in blood functionality pathways (**Fig. 5e**). Most interestingly, from the morphological features of Segment 7, we observed that this segment belongs to infiltrating glioblastoma cells characterized by elongate nuclei admixed with non-neoplastic brain cells. Glioblastoma cells showing elongated nuclei are characteristic of invasion along white matter tracts^23^. Comparing DEGs with pre-defined infiltrating markers^26^, we found that infiltrating tumor marker genes KCNN3 and CNTN1 were expressed specifically in Segment 7 (**Fig. 5f**). Overall, RESEPT successfully recognized tumor architecture, non-tumor architecture, and infiltration tumor architecture. This tool augments the morphological evaluation of glioblastoma by enabling an improved understanding of glioblastoma heterogeneity. This objective characterization of the heterogeneity will ultimately improve oncological treatment planning for patients.

**Fig. 5.**
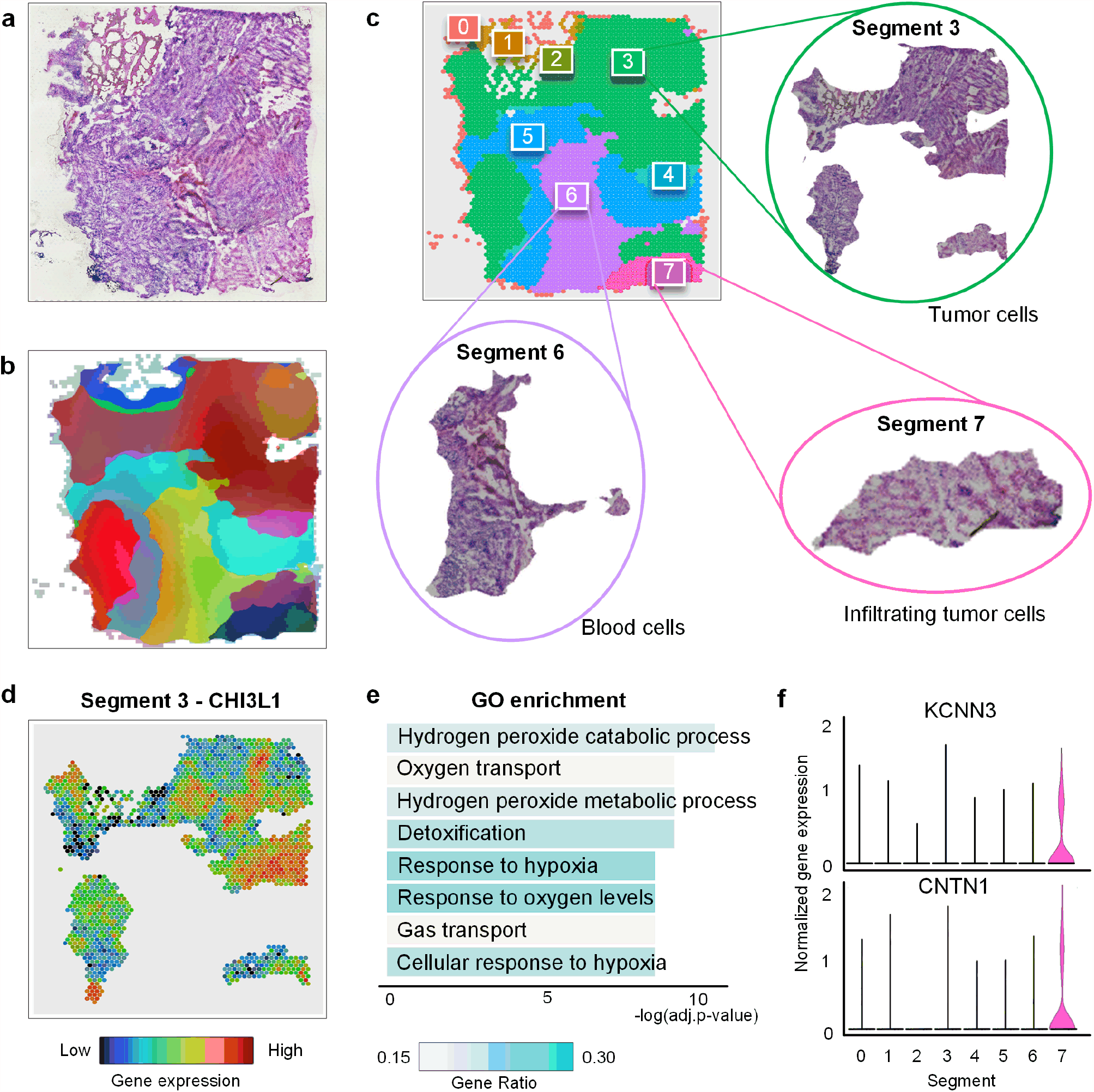
RESEPT identifies tumor regions in glioblastoma samples (Sample S1). (**a**) Original H&E staining image from the 10x Genomics. (**b**) RGB image generated from the RESEPT pipeline. (**c**) Labeled segmentation by RESEPT and Segments 3, 6, and 7 are cropped according to the segmentation result. Based on morphological features, our physiologist found Segment 3 contains large tumors from morphological features; Segment 6 contains a large number of blood cells; Segment 7 contains infiltrating tumor cells. (**d**) Glioblastoma marker gene CHI3L1 is highly and broadly expressed in Segment 3 based on the logCPM normalization value. (**e**) Bar plot shows the results of GO enrichment analysis, indicating Segment 6 having a large proportion of blood cells with blood signature genes for gas transport. (**f**) Infiltrating glioblastoma signature marker genes KCNN3 and CNTN1 are highly expressed in Segment 7 based on the logCPM normalization.

## Conclusion and Discussion

Our results show RESEPT is a robust and high-performance tool, for spatial transcriptomics data analysis, visualization, and interpretation. Powered by representation learning with graph neural networks in a spatial spot-spot graph model, the spatial transcriptomics is visualized as an RGB image. RESEPT formulates the problem as image segmentation and uses a deep-learning model to detect the tissue architecture. It has the potential to provide specific spatial architectures in broader applications, including neuroscience, immuno-oncology, and developmental biology.

RESEPT allows taking one of the two types of input, gene expression or RNA velocity. An RGB image generated from RNA velocity may have a different biological meaning from gene expression but is appropriate for some contexts, such as well-differentiated architectures in the spatial slice. For example, our study suggests well-structured brain cortical datasets like AD samples may have better performance in RNA velocity as input than gene expression. We will investigate the guideline on how and when to choose RNA velocity as the input instead of gene expression.

Besides RGB channels as the default setting, RESEPT can be adjusted to most mixing color pallets in graphic design, such as CMYK (Cyan, Magenta, Yellow, and blacK), HSV (Hue, Saturation, and Value), and hexadecimal colors. These alternative color systems may provide a broader color spectrum and enough variation in hue and brightness to present complex embedding. With these styles of visualization layouts as options, tissue architectures might be more accessible and distinguishable in some cases.

In the future, RESEPT will expand the methodology from lattice-based sequencing technologies including 10x and ST platform to fluorescence *in situ* hybridization (FISH) technologies, such as seqFISH and multiplexed error-robust FISH. With the availability of spatial multi-omics, RESEPT will also integrate other modals of information as histology image pixels together with the spatial coordinates and gene expression.

Meanwhile, RESEPT will be colorblind accessible with a ‘colorblind safe’ mode in visualization, in which all output images will be replaced with predefined color-blind palettes to avoid problematic color combinations. For different types of color blindness, RESEPT will offer corresponding narrow-down palettes accordingly. In addition, different patterns/labels instead of colors can be mapped in the image to distinguish among clusters.

## Methods

## 1. RESEPT pipeline

RESEPT is implemented in two major steps: (i) reconstruction of an RGB image of spots using gene expression or RNA velocity from spatial transcriptomics sequencing data; (ii) implementation of a pre-trained image segmentation deep-learning model to recognize the boundary of specific spatial domains and to perform functional zonation. **Fig. 1** and **Fig. 2** demonstrate the pipeline with conceptual description and technical details, respectively.

### 1.1 Construct RGB image for spatial transcriptomics

An RGB image is constructed to reveal the spatial architecture of a tissue slice using three-dimensional embedding as the primary color channels. Besides gene expression, RESEPT can accept RNA velocity^17^ as the input. RNA velocity unveils the dynamics of RNA expression at a given time by distinguishing the ratio of unspliced and spliced mRNAs, reflecting the kinetics and potential influences of transcriptional regulations in the present to the future cell state. The original BAM file of human studies is often unavailable to public users due to ethical reasons, and hence, in most cases, we only refer to expression-derived RGB images in our study. The scGNN^13^ package is used to generate spatial embeddings for each spot based on the pre-processed expression matrix or RNA velocity matrix along with the corresponding meta-data. In practice, RESEPT can adapt any type of low dimensional representations, such as embedding from UMAP and spaGCN^9^. On benchmarks, scGNN embedding obtained better results in most cases, so RESEPT uses scGNN in default (**Supplementary Fig. 8**).

#### Dimensional Reduction

After log-transformed and normalized library size by CPM, the spatial transcriptome expression or raw RNA velocity as the input is dimensionally reduced by learning a low dimensional embedding through an autoencoder. Both the encoder and the decoder consist of two symmetrically stacked layers of dense networks followed by the ReLU activation function. The encoder learns embedding *X*′ from the input matrix *X*, and the encoder reconstructs the matrix 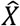 from the *X*, where *X* can be either gene expression or RNA velocity. Thus, 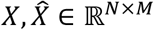 and *X*′ ∈ ℝ^*N*×*M*^′, where *M* is the number of input genes from the spatial transcriptome, *M*′ is the dimension of the learned embedding, and *M*′ < *M. N* is the number of spots of the spatial slide. The objective of the training is to achieve a maximum similarity between the original and reconstructed matrices through minimizing the mean squared error (**MSE**) 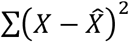 as the loss function. Positional encoding^27^ using Euclidean distance between spots on the tissue slice is also incorporated in reconstructing the input matrix.

#### Generating Spatial retained Spot Graph

The cell graph is a powerful mathematical model to formulate cell-cell relationships based on similarities between cells. In single-cell RNA sequencing (scRNA-seq) data without spatial information, the classical K-Nearest-Neighbor (KNN) graph is widely applied to construct such a cell-cell similarity network in which nodes are individual cells, and the edges are relationships between cells in the gene expression space. With the availability of spatial information in spots as the unit of observation arranged on the tissue slice, our in-house tool scGNN adopts spatial relation in Euclidean distance as the intrinsic edge in a spot-spot graph. Each spot in the spatial transcriptomics data contains one or more cells, and the captured expression or the calculated RNA velocity is the summarization of these cells within the spot. Only directly adjacent spots in contact in the 2D spatial plane have edges between them, and hence, the lattice of the spatial spots comprises the spatial spot graph. For the generated spot graph *G* = (*V, E*), *N* = |*V*| denoting the number of spots and *E* representing the edges connecting with adjacent neighbors. *A* is its adjacency matrix and *D* is its degree matrix, *i.e*., the diagonal matrix of number of edges attached to each node. The node feature matrix is the learned embedding *X*′ from the dimensional reduction autoencoder. In the 10x Visium platform, each spot has six adjacent spots, so the spatial retained spot graph has a fixed node degree of six for all the nodes. Similar to the KNN graph derived from scRNA-seq, each node in the graph contains *M*′ attributes.

#### Graph autoencoder

Given the generated spatial spot-spot graph, a graph autoencoder learns a node-wise three-dimensional representation to preserve topological relations in the graph. The encoder of the graph autoencoder composes two layers of graph convolution network (GCN) to learn the low dimensional graph embedding *Z* in *Eq*. (1).

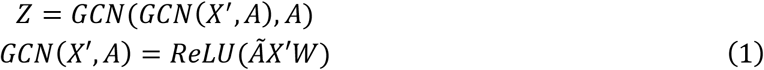

where *Ã* = *D*^−1/2^*AD*^−1/2^ is the symmetrically normalized adjacency matrix and *W* is a weight matrix learned from the training. The output dimensions of the first and second layers are set as 32 and 3, according to the 3 color channels as RGB, respectively. The learning rate is set at 0.001.

The decoder of the graph autoencoder is defined as an inner product between the graph embedding *Z*, followed by a sigmoid activation function:

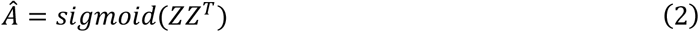

where *Ã* is the reconstructed adjacency matrix of *A*.

The goal of graph autoencoder learning is to minimize the cross-entropy *L* between the input adjacency matrix *A* and the reconstructed matrix *Ã*.

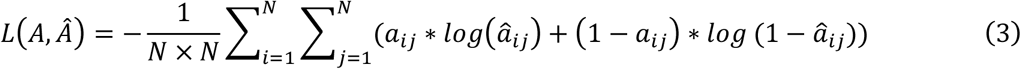

where *a*_*ij*_ and *â*_*ij*_ are the elements of adjacency matrix *A* and *Ã*,1 ≤ *i* ≤ *N*, 1 ≤ *j* ≤ *N*. As there are *N* nodes as the number of spots in the slide, *N* × *N* is the total number of elements in the adjacency matrix.

#### Reconstruct RGB Image

The learned embedding *Z* ∈ ℝ^*N*×3^ is capable of representing and preserving the underlying relationships in the modeled graph from spatial transcriptomics data. Meanwhile, the three-dimensional embedding can also be intuitively mapped to Red, Green, and Blue channels in the RGB space of the image. Normalized to an RGB color space accordingly to a full-color spectrum (pixel range from 0 to 255) as Eq. (4), the embedding of each spot is assigned a unique color for exhibiting the expression or velocity pattern in space.

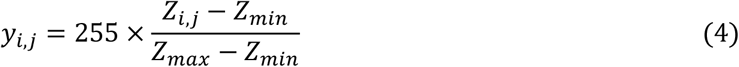

where *y* ∈ ℝ^*N*×3^ and *y*_*i,i*_ is its transformed color of the *i*-th spot in the *j*-th channel, 1 ≤ *i* ≤ *N, j* ∈ {*R, G, B*}. *Z*_*min*_ and *Z*_*min*_ represent the maximum and minimum of all embedding values in the RGB channels, respectively. With their coordinates and diameters at the full resolution provided from 10x Visium, we are able to plot all spots with their synthetic colors on a white drawing panel and reconstruct a full-size RGB image explicitly describing the spatial expression or velocity properties in the original spatial coordinate system. For the spatial transcriptomic data sequenced in lattice from other techniques as ST platform, RESEPT allows users to specify a diameter to capture appropriate relations between spots in the RGB image accordingly.

### 1.2 RGB image segmentation model

The RGB image makes the single-cell spatial architecture perceptible in human vision. With the constructed image, we treat the potential functional zonation partition as a semantic segmentation problem, which automatically classifies each pixel of the image into a spatially specific segment. Such predictive segments reveal the functional zonation of spatial architecture.

#### Image segmentation model architecture

We trained an image-segmentation model based on a deep architecture DeepLabv3+ ^28,29^, which includes a backbone network, an encoder module, and a decoder module (**Fig. 2**).

#### Backbone network

The backbone network provides dense visual feature maps for the following semantic extraction by any deep convolutional network. Here, ResNet-101^30^ is selected as the underlying model for the backbone network, which consists of a convolutional layer with 64-channels in 7 × 7 size of filters and 33 residual blocks, each of which stacks one convolutional layer with multi-channel (including 64, 128, 256, and 512) in 3 × 3 size of filters and two convolutional layers with multi-channel (including 64, 128, 256, 512, 1024 and 2048) 1 × 1 size of filters. The generated RGB image is mapped into a *c* -channel feature map by the first convolutional layer and gradually fed into the following residual blocks to produce rich visual feature maps for describing the image from different perspectives. Here, *c* equals 64. In each residual block, the feature map generated from the previous block *y* ∈ ℝ^*N*×3^ is updated to *y* ∈ ℝ^*N*×*c*^ in Eq. (5).

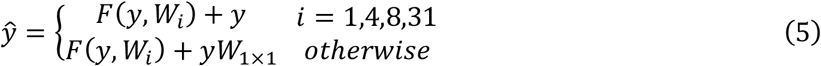

where

- *F*(*) is the activation function, and we use ReLU ^31^ in this study.
- *W*_*i*_ represents the learning convolutional weights in the *i*^*th*^ block,1 ≤ *i* ≤ 33.
- *W*_1×1_ represents the learning weights of the convolutional layer with 1×1 kernel size.

Element-wise addition operation *F* + *y* in Eq. (5) enables a direct shortcut to avoid the vanishing gradient problem in this deep network. In the 1^st^, 4^th^, 8^th^, and 31^st^ blocks of the 33 residual blocks, their input and output dimensions do not match up due to different filter settings from their previous layers. Accordingly, the projection shortcut with an additional 1×1 convolution in Eq. (5) is used to align dimensions in these blocks, which are also named identity blocks. The rest blocks stacked on the previous blocks with the same filter settings employ a direct shortcut. We leveraged ResNet-101 as a basic visual feature provider and sent the most informative feature maps from the last convolutional layer before logits to the following encoder module.

#### Encoder module

The aim of the encoder module is to capture multi-scale contextual information based on the dense visual feature maps from the backbone. To achieve the multi-scale analysis, atrous convolution^28^ is adopted in the encoder to extend the size of the respective field. For the generated RGB image with width *i* and length *n*, the total number of spots *N* = *i* × *n*. Given the input signal from Eq. (5) as *y* ∈ ℝ^*m*×*m*×*c*^ with a *c*′-channel filter *e* ∈ ℝ^*K*×*K*×*c*^′, the output feature signal *y*′ ∈ ℝ^*m*×*m*×*c*^′ is defined as follows:

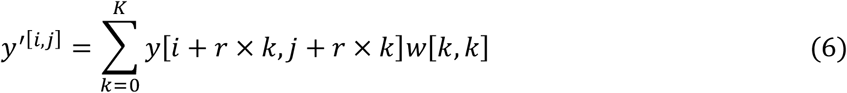

where

- *y*[*i, j*] represents the input signal at the location *(i, j)* with *c-*channel values. 0 ≤ *i* ≤ *i*, 0 ≤ *j* ≤ *n* . *e* is the stride rate in atrous convolution.
- *e*[*k, k*] represents the convolutional weights with *c’*-channel values, 0 ≤ *k* ≤ *K* . *K* is the kernel size of the convolutional filter.
- *y*′[*i, j*] represents the output signal at the location *(i, j)* with *c’-*channel values.

Compared to the standard convolution, the atrous convolution samples the input signal *y* with the stride *r* rather than using direct neighbors inside the convolutional kernel. Therefore, the standard convolution is a special case of atrous convolution with *r = 1*. By using multiple rate value settings (rate = 1, 6, 12 and 18), we separately apply one standard convolutional layer with 256-channel 1 × 1 size of filters (i.e., the atrous convolutional layer with rate = 1), three atrous convolutional layers with 256-channel 3 × 3 size of filters and an additional average pooling layer to produce high-level multi-scale features. These semantic features are then merged into the decoder module.

#### Decoder module

In the decoder, the input high-level features are bilinearly up-sampled and concatenated with the basic visual features for recovering the segment boundaries and spatial dimension. A standard convolutional layer with 256-channel 3 × 3 size of filters is applied to outweigh the importance of the merged features and obtain sharper segmentation results. Eventually, an additional bilinear up-sampling operation forms the output of the decoder to a *m* × *n* × 256 matrix, where *m* and *n* denote the width and height of the input image, respectively. The following convolution layer with *d*-channel 1 × 1 size of filters squeezes the feature matrix along the channel axis to *m* × *n* × *d* shape, where *i* is the pre-defined maximum number of categories. The softmax^32^ function is then applied to generate its predictive segmentation map, which takes a matrix with the same size of the input image recording the segment category of each pixel on it. The pixels falling into a certain category in the segmentation map point to a segmented spatial region. Our modeling objective is to minimize the cross-entropy^33^ between the predictive segmentation map *Ŝ* and labeled spatial functional regions *S*:

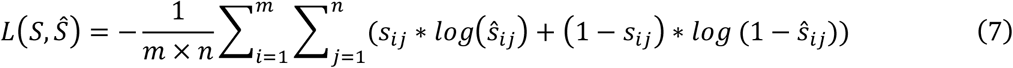

where *i*_*ij*_ and ŝ_*ij*_ are the segment categories of the pixel at the *i*-th row and the *j*-th column for the input images with *m* × *n* pixels. *s*_*ij*_ ∈ [1, *i*], ŝ_*ij*_ ∈ [1, *d*].

#### Training set data preparation

We performed scGNN using various autoencoder dimensions (*M′*= 3, 10, 16, 32, 64, 128, and 254) and multiple positional encoding intensity parameters (*PE α* = 0.1, 0.2, 0.3, 0.5, 1.0, 1.2, 1.5, and 2.0), resulting in 56 embeddings used to generate diverse RGB images for each sample in the training set (see image results on https://github.com/OSU-BMBL/RESEPT). In this study, we performed 16-fold Jackknife cross-validation, each of which formed all but one observation as the training set. The one sample was left to evaluate the trained model in each fold.

#### Model training

We implemented the training procedure on the MMSegmentation platform^34^, which is an open-source semantic segmentation toolbox based on PyTorch. The weights of DeepLabv3+ were initialized by the pre-trained weights from the Cityscapes dataset provided by MMSegmentation. To introduce diversity to the training data and improve the generalization of our model, we applied transforms defined in MMSegmentation, including the random cropping, rotation and photometric distortions, to augment the training RGB images. *400 × 400* sized patches are randomly cropped to provide different regions of interest from the whole RGB images. A random rotation (range from -180 degrees to 180 degrees) was further conducted to fit the potential irregular layout of spatial architectures. Some photometric distortions such as brightness, contrast, hue, and saturation changes were also utilized to training samples when loading to MMSegmentation. Stochastic gradient descent (SGD)^35^ was chosen as the optimization algorithm, and its learning rate was set to 0.01. The training procedure iterated 30 epochs, and the checkpoint among all epochs with the best Moran’s I autocorrelation index^36^ on the testing data was selected as the final model.

#### Image segmentation inference

Once a model completes training, it is capable of predicting the functional zonation on the tissue from its RGB images. On the inference, RESEPT performs scGNN with the same parameter combinations with the training settings resulting in 56 candidate RGB images for each input sample. RESEPT infers all the segmentation maps on these 56 images and scores them using the Moran’s I metric (details in **Supplementary Fig. 9**) to assess the quality of segmentations. The segmentation maps of 5-top ranked images in terms of Moran’s I are returned for user selection. We found that such a quality assessment protocol results in segmentation results with higher accuracy than the default one and enhances the robustness of RESEPT.

## 2 Data analysis

### 2.1 Experiment preparation, data generation, and processing

#### Experiment preparation and data generation

Four postmortem human brain samples of the middle temporal gyrus^16^ were obtained from the Arizona Study of Aging and Neurodegenerative Disorders/Brain and Body Donation Program at Banner Sun Health Research Institute^37^ and the New York Brain Bank at Columbia University Medical Center^38^. Two of them are from non-AD cases at Braak stage I-II, namely Samples S2 and S5 in the study, and the other two are from early-stage AD cases at Braak stage III-IV, namely Samples S4 and S3 in the study. The region of AD cases was chosen based on the presence of Aβ plaques and neurofibrillary tangles. The 10x Genomics Visium Spatial Transcriptome experiment was performed according to the User Guide of 10x Genomics Visium Spatial Gene Expression Reagent Kits (CG00239 Rev D). All the sections were sectioned into 10 µm thick and mounted directly on the Visium Gene Expression (GE) slide for H&E staining and the following cDNA library construction for RNA-Sequencing. Besides the section mounted on the GE slide, one of the adjacent sections (20 µm away from GE section) from AD samples persevered for the Aβ immunofluorescence staining. The method of immunofluorescence staining of Aβ on persevered section was the same as previously described^20^. The image of Aβ staining was used as the ground truth and was aligned to H&E staining on GE slides using the “Transform/Landmark correspondences” plugin in ImageJ^39^.

#### FASTQ generation, alignment, and count

BCL files were processed by sample with the SpaceRanger (v.1.2.2) to generate FASTQ files via spaceranger *mkfastq*. The FASTQ file was then aligned and quantified based on the reference GRCh38 Reference-2020-A via spaceranger count. The functions spaceranger *mkfastq* and spaceranger *count* were used for demultiplexing sample and transcriptome alignment via the default parameter settings.

### 2.2 Data preprocessing

To standardize the raw gene expression matrix and spot metadata, the different spatial transcriptomics data were preprocessed as follows.

For the 10x Visium data (**Table 1**), the filtered feature-barcode matrix (HDF5 file) was reshaped into a two-dimensional dense matrix in which rows represent spots and columns represent genes. The dense matrix was further added with spots’ spatial coordinates by merging them with the ‘tissue_positions_list’ file, containing tissue capturing information, row, and column coordinates. The mean color values of the RGB channels for each spot’s circumscribed square and annotation label were also added to the dense matrix after processing the Hematoxylin-Eosin (H&E) image. The gene expression as part of the dense matrix was stored in a sparse matrix format. Other information describing the spots’ characteristics was stored as individual metadata.

For the HDST data, the expression matrix and spots’ coordinates were reshaped into the dense matrix, which was similar to 10x Visium preprocessing. The expression matrices from dense matrices were formed into the individual sparse matrices, and other information was stored as metadata.

For the ST data, the expression matrix was reshaped into the two-dimensional dense matrix, and spots’ spatial coordinates were added to the dense matrix by merging with the spot_data_selection file. The color values of each spot were added to the dense matrix after processing the H&E image (if available). The remaining steps were the same as for the 10x Visium data.

### 2.3 Data normalization and denoising

#### Data normalization

The raw read counts were used as formatted input to generate normalization matrices. Seven normalization methods were used in the study, including DEseq2^40^ (v.1.30.1), scran^41^ (v.1.18.5), sctransform^42^ (v.0.3.2), edgeR^43^ (v.3.32.1), transcripts per million (TPM), reads per kilobase per million reads (RPKM), and log-transformed counts per million reads^44^ (logCPM). We used Seurat (v.4.0.1) to generate the sctransform and the logCPM normalized matrices. edgeR was used to generate TMM^43^ normalized matrices. The gene length was used for calculating TPM, and RPKM was obtained from biomaRt (v.2.46.3) by using *useEnsemble* function and parameters setting as dataset=“hsapiens_gene_ensembl” and GRCh=38. All normalized matrices for whole transcriptomics were eventually calculated via the following default settings and converted into sparse matrices. RNA velocity was calculated for the whole transcriptomics via velocyto^17^ (v.0.17.17) and scVelo^18^ (v.0.1) followed by their default settings. RNA velocity matrices were converted into sparse matrices.

#### Missing spots imputation

In practice, several spots may have missing expression in some tissue slices due to imperfect technology, which leads to blank tiles at the locations of these spots on the RGB images. Such blank tiles as incompatible noises may skew the following boundary recognition of spatial architecture. We assume the near neighbors are more likely to have similar values to the missing spot and impute these missing spots by applying the weighted average to the pixels of their valid six neighboring spots. Since these missing spots are colored while in default as the same with the background out of tissue, we need to distinguish them from all-white pixels according to a topological structural analysis^45^. Firstly, all contours (including outer contours of tissue and inner contours caused by missing spots) of tissue are detected from the border following procedure^45^. The contour with the largest area is determined as the outer contour of tissue. Then, all pixels in white inside the tissue contour are replaced by imputation from their neighbors. Given missing spot coordinates, we search their nearest *k* valid spots *s*_*i*_ *(i =*1, 2,…, *k)* to calculate the imputation value *x* _*s*_ of target missing spot *s* as:

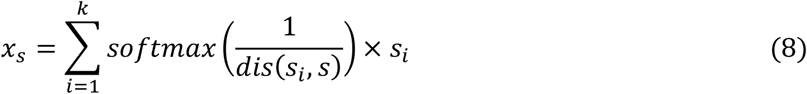

where *dis(s*_*i*_, *s*) represents the Euclidean distance between target spot *s* and a certain neighbor *s*_*i*_ in spatial space. The softmax function normalizes all distance reciprocals of *s* and its *k* (we set *k=*6 by default) neighbors *s*_*i*_ to the weights ranging from 0 to 1. The imputation of *s* is the weighted average on all *s*_*i*_. If a tissue slice is detected without missing spots, RESEPT skips this imputation process.

#### Parameter setting

Parameters in scGNN to generate embedding are referred to in the previous study^13^. In the case study of the AD sample, in analysis on cortical layers 2 & 3, the expressions of 8 well-defined marker genes were log-transformed and embedded by spaGCN with 0.65 resolution. In the analyses of cortical layer 2 to layer 6, PCA (*n.PCs=3*) was firstly utilized to extract the principal components of their expressions of marker genes for highlighting the dominant signals, and then they were embedded by spaGCN with 0.65 resolution. In the exploration of tumor regions in glioblastoma samples, their marker gene expressions were preprocessed by logCPM normalization and PCA (*n.PCs=50*). The processed data was embedded by spaGCN with 0.35 resolution. In the analyses of AD-associated critical cell types, marker gene expressions were preprocessed by log-transform and PCA (*n.PCs=3*) as well and then embedded by spaGCN with 0.65 resolution. For investigating Aβ pathological regions, log-transform to the expressions of validated 20 upregulated genes was applied, and their embedding was generated by spaGCN with 0.65 resolution.

## 3 Benchmarking evaluation

All the benchmarking tasks were run on a Red Hat Enterprise Linux 8 system with 13 T storage, 2x AMD EPYC 7H12 64-Core Processor, 1TB RAM 1TB DDR4 3200MHz RAM, and 2x NVIDIA A100 GPU with 40GB RAM. The usage of the existing tools and their parameter settings in our benchmarking evaluation were described below.

*Seurat (v.4.0.1)* identifies tissue architecture based on graph-based clustering algorithms (e.g., Louvain algorithm). Creating Seurat object, identification of highly variable features, and scaling of the data was performed using default parameters. The PCs were set to 128 to match our framework default setting. The *FindNighbors* and *FindClusters* functions with default parameters were used for tissue architecture identification. To further evaluate the robustness of the combination of the different parameters, we used 16 samples and selected three important parameters, including the number of PCs (dims = 10, 32, and 64), the value of *k* for the *FindNeighbor* function (*k.parm* = 20, 50 and 100), and the resolution in the *FindClusters* function (*res* = 0.1 to 1, step as 0.1).

*BayesSpace (v.1.0.0)* identifies tissue architecture based on the Gaussian mixture model clustering and Markov Random Field at an enhanced resolution of spatial transcriptomics data. Creating the *SingleCellExperiment* object is implemented to the following analysis by loading normalized expression data and position information for barcodes. Then, we set 128 as the number of PCs in *spatialPreprocess* function and parameter *log.normalize* was set FALSE due to the normalized data input. Lastly, tissue architecture was identified by running *qTune* and *spatialCluster* functions. We followed the official tutorial and adopted k-means as initial methods while other parameters were from the default based on prior information. In the process of assessing the robustness of BayesSpace, we set the cluster number as seven, the parameter *n.PCs* in spatialPreprocess function (*n.PCs* = 10, 64, and 128), and the parameter *nrep* in spatialCluster function (*nrep* = 5000, 10000, and 150000) for 16 samples.

*SpaGCN (v.0.0.5)* can integrate gene expression, spatial location, and histology to identify spatial domains and spatially variable genes by graph convolutional network. SpaGCN was used to generate 3D embedding and tissue architecture and includes three procedures, including loading data, calculating adjacent matrix, and running SpaGCN. In the first step, both expression data and spatial location information were imported. Second, adjacent matrices were calculated using default parameters. Lastly, we selected 128 PCs, the initial clustering algorithm as Louvain, and other parameters used default settings. To evaluate the robustness of the parameters and enable comparison with other tools, three parameters, the number of PCs (*num_pcs* = 20, 30, 32, 40, 50, 60, 64), the value of k for the k-nearest neighbor algorithm (*n_neighbors* = 20, 30, and 40), and the resolution in the Louvain algorithm (res = 0.2, 0.3, and 0.4) for 16 samples were adjusted.

*stLearn (v.0.3.2)* is designed to comprehensively analyze ST data to investigate complex biological processes based on Deep Learning. stLearn highlights innovation to normalize data. Therefore, we input expression data, location information as well as images. stLearn consists of two steps, i.e., preparation and run stSME clustering. In preparation, loading data, filtering, normalization, log-transformation, preprocessing for spot image, and feature extraction were implemented. In the following module, PCA dimension reduction was set to 128 PCs, applying stSME to normalize log-transformed data and Louvain clustering on stSME normalized data using the default parameters. To evaluate the robustness of the parameters and enable comparison with other tools, three parameters were considered to be adjusted for 16 samples, the number of PCs (*n_comps* = 10, 20, 30, 32, 40, and 50), the value of k for the kNN algorithm (*n_neighbors* = 10, 20, 30, 40, and 50), and the resolution in the Louvain algorithm (resolution = 0.7, 0.8, 0.9 and 1).

*STUtility (v0.1.0)* can be used for the identification of spatial expression patterns alignment of consecutive stacked tissue images and visualizations. We implemented STUtility as a tissue architecture tool based on the Seurat framework. *RunNMF* was carried out as the dimension reduction method. The number of factors was set to 128 for matching our framework default setting. *FindNeighbors* and *FindClusters* were used to identify tissue architecture. To further evaluate the robustness of the combination of the different parameters, we used 16 samples and selected three important parameters for tuning, including the number of factors (*nfactors* = 10, 32, and 64), the value of *k* for *FindNeighbor* function (*k.parm* = 20, 50, 100, 200, and 250), and the resolution in *FindClusters* function (res =0.05, 0.1, 0.2, 0.3, 0.5, and 0.7, 0.9).

*Giotto (v.1.0.3)* is a comprehensive and multifunction computational tool for spatial data analysis and visualization. We implemented Giotto as the issue architecture identification tool in this study via using default settings. Giotto first identified highly variable genes via calculateHVG function, then performed PCA dimension reduction using 128 PCs, constructed the nearest neighbor network via *createNearestNetwork*, and eventually identified tissue architecture via *doLeidenCluster*. To further evaluate the robustness of the combination of the different parameters, we used 16 samples and selected three important parameters for tuning, including the number of PCs (*npc* = 10, 32, and 64), the value of k for *createNearestNetwork* function (*k* = 20, 50 and 100), and the resolution in *doLeidenCluster* function (*resolution* = 0.1, 0.2, 0.3, 0.4, and 0.5).

### Downsampling simulation for read depth

Comparing the mean and standard deviation of 16 10x Visium datasets, samples S5 and S6 were selected to generate simulation data with decreasing sequencing depth. Let matrix *G* be the *N* × *M* expression count matrix, where *N* is the number of spots and *M* is the number of genes. Define the spot-specific sequencing depths 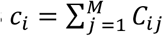 i.e., the column sums of *G*. Thus, the average sequencing depth of the experiment is 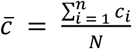 Let 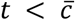 be our target downsampled sequencing depth and let C^*^ be the *N* × *M* downsampled matrix. We perform the downsampling as follows:

For each spot *i* = 1, …, *N*:

1. Define the total counts to be sampled in the spot i as 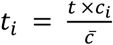
2. Construct the character vector of genes to be sampled as 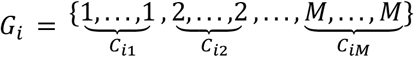
3. Sample t_*i*_ elements from *G*_*i*_ without replacement and define *N*_*i*_ as the number of times gene *j* was sampled from *G*_*i*_ for *j* = 1, …, *M*.
4. Let 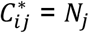.

Using this method, the average downsampled sequencing depth is:

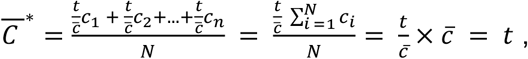

as desired. Note also that this method preserves the relative total counts of each spot, i.e., spots that had higher sequencing depths in the original matrix have proportionally higher depths in the downsampled matrix.

## 4 Evaluation metrics

### 4.1 Benchmark performance evaluation criteria

Adjusted Rand Index (ARI), Rand index (RI), Fowlkes–Mallows index (FM), and Adjusted mutual information (AMI) are used to evaluate the performances between the ground truth and predicted results.

Adjusted Rand index (ARI) measures the agreement between two partitions. Given a set *S* consisting of *n* elements, ℱ_1_ = {*X*_1_, *X*_2_, …, *X*_*r*_} and ℱ_2_ = {*Y*_1_, *Y*_2_, …, *Y*_*i*_} are two partitions of *S*; that is, *S* =∪_*i*_ *X*_*i*_ and *X*_*i*_ ∩ *X*_*i*_ = ∅, so does ℱ_2_. *X*_*i*_ can be interpreted as a cluster generated by some clustering method. In this way, ARI can be described as follow:

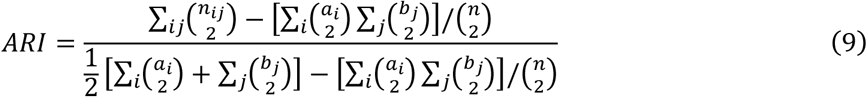

where *n*_*ij*_ = *X*_*i*_ ∩ *Y*_*j*_, denotes the number of objects in common between *X*_*i*_ and *Y*_*j*_; *a*_*i*_ = ∑_*i*_ *n*_*ij*_ and *b*_*i*_ = ∑_*i*_ *n*_*ij*_. Besides, *ARI* ∈ [−1, 1], the higher *ARI* reflects the higher consistency. The bs function of the splines package (v.4.0.3) was used for smoothing ARI generated from grid effective sequencing depth data via default settings.

### Rand index (RI)

is also a measure of the similarity between two data clustering results. If the ground truth is available, the *R* can be used to evaluate the performance of one cluster method by calculating *R* between the clustering produced by this method and the ground truth. Let S be a set containing *n* elements, which represents *n* barcodes in this paper, and two partitions of *S*, ℱ_1_ = {*X*_1_, *X*_2_, …, *X*_*r*_}, ℱ_2_ = {*Y*_1_, *Y*_2_, …, *Y*_*i*_}; that is, *S* =∪_*i*_ *X*_*i*_ and *X*_*i*_ ∩ *X*_*i*_ = ∅; so does ℱ_2_. *X*_*i*_ and Y_j_ are the subset of *S*, representing one cluster produced by some clustering method and the ground truth, respectively. *R* can be computed using the following formula:

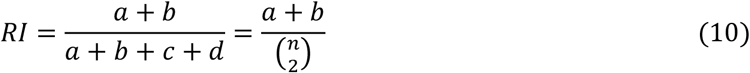

where:

- *a, b, c, i* denote the number of pairs of elements in *S* in the same subset in ℱ_1_ and in the same subset in ℱ_2_, in different subsets in ℱ_1_ and in different subsets in ℱ_2_, in the same subset in ℱ_1_ and in different subsets in ℱ_2_, and in different subsets in ℱ_1_ and in the same subset in ℱ_2_, respectively.
- ^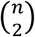^ is the binomial coefficient. In addition, the range of *RI* is [0,1], and the higher *RI*, the higher similarity of two partitions is.

### The Fowlkes–Mallows index (FM)

is an external evaluation method, which can measure the results’ consistency of two cluster algorithms. Not only can FM be implemented on two hierarchical clusterings, but also the clusters and the benchmark classifications. For the set S of n objects, *A*_1_ and *A*_2_ denote two clustering results (generated by two cluster algorithms or one for cluster algorithm, one for the ground truth). In this paper, A_1_ is produced by a clustering algorithm while the ground truth contributes A_2_. If the clustering algorithm performs well, then *A*_1_ and *A*_2_ should be as similar as possible. The calculation of FM can be described as:

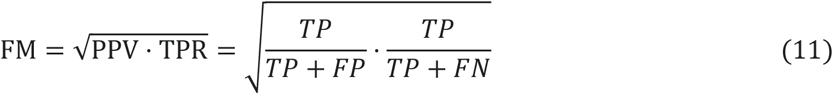

where

- TP is the number of true positives, representing the number of pair objects that are present in the same cluster in both *A*_1_ and *A*_2_.
- FP is the number of false positives, representing the number of pair objects that are present in the same cluster in *A*_1_ but not in *A*_2_.
- TN is the number of false negatives, representing the number of pair objects that are present in the same cluster in *A*_2_ but not in *A*_1_.
- PPV is so-called precision while *TPR* refers to recall. In addition, FM ∈ [0, 1]. Therefore, in our cases, the closer it is to 1, the better the clustering algorithm will be.

### Adjusted mutual information (AMI)

is driven from probability theory and information theory and can be used for comparing clustering results. To introduce adjusted mutual information, the preliminary is necessary to present two conceptions mutual information (MI) and entropy. Given a set S = {s_1_, s_2_, …, s_n_}, ℱ_1_ = {*X*_1_, *X*_2_, …, *X*_*r*_} and ℱ_2_ = {*Y*_1_, *Y*_2_, …, *Y*_*i*_} are two partitions of *S*, that is, *S* =∪_*i*_ *X*_*i*_ and *X*_*i*_ ∩ *X*_*i*_ = ∅, so does ℱ_2_. MI between partition ℱ_1_ and ℱ_2_ is defined as:

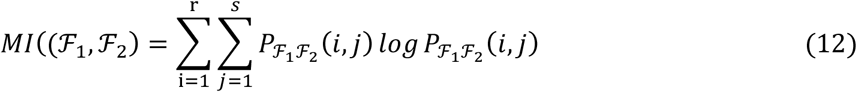

where

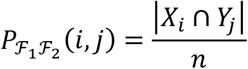

measures the probability of one object belonging to *X*_*i*_ and *Y*_*i*_ simultaneously. The entropy associated with the partitioning ℱ_1_ is defined as:

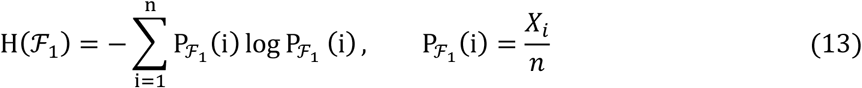

where

- 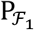 (i) refers to the probability that the object falls into the cluster *X*_*i*_.
- H(ℱ_2_) and 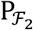 (*j*) have analogous definitions.

The following formula shows the expected mutual information between two random clustering results:

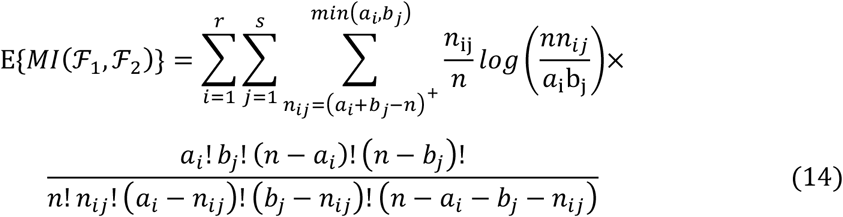

where *a*_*i*_ + *b*_*i*_ − *n* = *max*1, *a*_*i*_ + *b*_*i*_ − *n*; *a*_*i*_ = ∑_*i*_ *n*_*ij*_ and *b*_*j*_ = ∑_*j*_ *n*_*ij*_, *n*_*ij*_ = *X*_*i*_ ∩ *Y*_*i*_, represents the number of objects in common between *X*_*i*_ and *Y*_*j*_. Finally, AMI can be obtained by

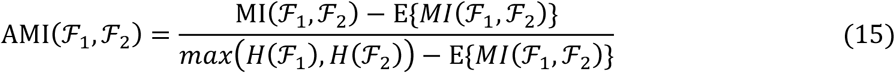

It should be pointed out that AMI ∈ [0, 1], the similarity between the two clusterings increases with the augment of AMI.

### 4.2 RGB image and 3D embedding evaluation

We modified the metric peak signal-to-noise ratio (PSNR)^46^, whose original version is commonly used to measure the reconstruction loss of image compression, to assess the similarity between the color distribution of an RGB image and its corresponding labeled segmentation map. We re-used its basic concept to calculate the PSNR from each labeled segment, and then applied weighted sum to the PSNRs from all *p* segments according to their area:

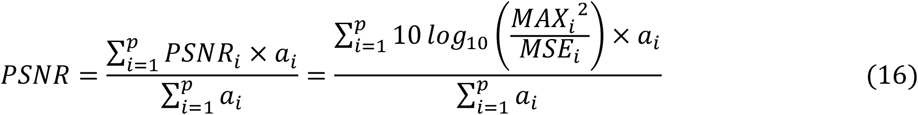

where:

- *a*_*i*_ is the number of pixels located in the *i*^*th*^ segment, 1 ≤ *i* ≤ *p*
- *MAX*_*i*_ is the maximum pixel-value of the *i*^*th*^ segment, 0 ≤ *MAX* ≤ 255
- *MSE*_*i*_ is the pixel-wise mean squared error of the *i*^*th*^ segment.

The larger PSNR implies the better the RGB image can indicate the labeled spatial architectures, and further demonstrates the better quality its corresponding 3-dimensional embeddings achieve.

### 4.3 Predicted Segmentation Map Quality assessment

Differed from the Moran’s I auto-correlation index^36^ using for revealing a single gene’s spatial auto-correlation, we modified Moran’s I in Geo-spatiality^47^ to evaluate a predictive segmentation map without known ground truth. The metric analyzes the heterogeneity of predictive inter-segments by measuring the pixel contrast cross any two predicted adjacent segments per channel:

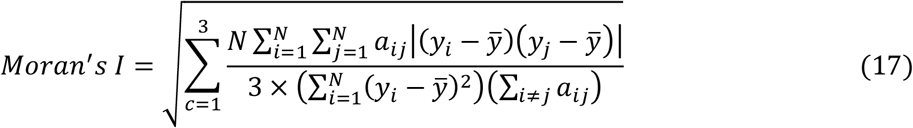

where

- *a*_*ij*_ is the binary spatial adjacency of the *i*^*th*^ segment and *j*^*th*^ segment . 1 ≤ *i* ≤ *N*, 1 ≤ *j* ≤ *N*
- *y*_*i,c*_ ∈ ℝ^3^ denotes the mean pixel values at *c*^*t*h^ channels in Red, Green and Blue of the *i*^*th*^segment, 1 ≤ *c* ≤ 3,
- 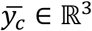 denotes the mean pixel values at channel Red, Green and Blue of the whole image.

### 4.4 Module score calculation and differential expression analysis

The module score for specific marker genes was calculated based on the Seurat function *AddModuleScore*, which calculated the average expression levels of genes for specific spot groups. The DEG analysis was conducted by the Seurat function FindAllMarkers based on RESEPT predicted seven segments via default settings. Based on the identified DEGs, the enrichment analyses of GO terms (Biological Process) and KEGG were performed via the R package clusterProfile (v.3.18.0) using the functions of enrichGO and enrichKEGG. The enrichment analysis results were filtered out if the adjusted p-value was greater than 0.05. For KEGG analysis, gene database Org.Hs.eg.Db was used for transferring SYMBOL to ENREZID via function bitr. R package ggplot2 (v.3.3.2) was used for the visualizations.

## Data availability

The 10x Visium datasets (10 from Spatial Gene Expression 1.0.0; 14 from Spatial Gene Expression 1.1.0, 13 from Spatial Gene Expression 1.2.0; including S1) can be accessed from https://www.10xgenomics.com/products/spatial-gene-expression. Our own AD datasets (S2-S5) are available from Dr. Hongjun Fu upon request. The datasets (S6-S17) used for training model and benchmarking can be accessed via endpoint “jhpce#HumanPilot10x” on Globus data transfer platform at http://research.libd.org/globus/. The HDST datasets are available as accession number SCP420 in the Single Cell Portal via link https://singlecell.broadinstitute.org/single_cell. The ST and 10x Visium data (squamous cell carcinoma) can be accessed from the GEO database with an accession number GSE144239. More details of datasets can be found in Supplementary Table 1.

## Code availability

RESEPT is freely available as an open-source Python package at https://github.com/OSU-BMBL/RESEPT.

## Contributions

Conceptualization: D.X. and Q.M.; methodology: F.H, J.W., Y.C., Q.M. and D.X.; software coding: F.H, Y.C, J.L, Y.Y, L.S., J.W. and L.Y.; data collection and investigation: Y.C., S.C. and L.S.; data generation: S.S.; data analysis and visualization: Y.C., F.H, J.L., Y.Y., J-X.L, L.S., S.C., Y.L. and A.M.; AD result interpretation: H.F.; Glioblastoma result interpretation: J.O.; software testing and tutorial: Y.Y.; Simulation: C.A. and D.C.; manuscript writing, review, and editing: J.W., Y.C., F.H., B.L., C.A., D.C., Z.L., D.X., C.A., D.C. and Q.M.

## Acknowledgements

This work was supported by awards R35-GM126985 and R01-GM131399 from the National Institute of General Medical Sciences and awards K01-AG056673 and R56-AG066782-01 from the National Institute on Aging of the National Institutes of Health. The work was also supported by award NSF1945971 from the National Science Foundation and the award of AARF-17-505009 from the Alzheimer’s Association. We thank Hua Li and Shiyuan Chen from Stowers Institute, Liangping Li from the Ohio State University for helpful discussion, and Paul Toth from Ohio State University for polishing the manuscript. Human de-identified brain tissues were kindly provided by the Banner Sun Health Research Institute Brain and Body Donation Program, supported by NIH grants U24-NS072026 and P30-AG19610 (TGB), the Arizona Department of Health Services (contract 211002, Arizona Alzheimer’s Research Center), the Arizona Biomedical Research Commission (contracts 4001, 0011, 05-901 and 1001 to the Arizona Parkinson’s Disease Consortium) and the Michael J. Fox Foundation for Parkinson’s Research and the New York Brain Bank at Columbia University Medical Center. This work used the high-performance computing infrastructure at the Ohio State University and the University of Missouri, as well as the Extreme Science and Engineering Discovery Environment (XSEDE), which is supported by the National Science Foundation grant number ACI-1548562.

