## Supplementary Figures and Tables for "Define and visualize pathological architectures of human tissues from spatially resolved transcriptomics using deep learning"

***Supplementary Fig. 1:*** *Abstract of RGB images & architecture detected by RESEPT based on LogCPM normalization.*

***Supplementary Fig. 2:*** *RGB images of Sample S6 with different read depth.*

***Supplementary Fig. 3:*** *RGB images of data in 10x, ST, and HDST platforms without manual annotation.*

***Supplementary Fig. 4:*** *ARI metrics from scGNN and spaGCN on 16 samples.*

***Supplementary Fig. 5:*** *QA metrics from scGNN, spaGCN, and UMAP on 16 samples.*

***Supplementary Fig. 6:*** *QA metrics for different parameter settings in 16 samples.*

***Supplementary Fig. 7:*** *ARI error bar changes according to different training sets.*

***Supplementary Fig. 8:*** *RGB image of other layer-specific markers on the AD sample S4 (update using non-normalization data).*

***Supplementary Fig. 9:*** *Other cell-specific markers on the AD sample S4.*

***Supplementary Fig. 10:*** *Other 4 section morphological features and transcriptional features on glioblastoma sample S1.*

**Supplementary Tables**

***Supplementary Table 1:*** *Summary of Datasets used in the study.*

***Supplementary Table 2:*** *Marker genes in AD.*

**Supplementary Data (individual files)**

***Supplementary Data 1:*** *ARI and tuned parameter usage of tools for generating exact 7 clusters.*

***Supplementary Data 2:*** *Time and memory cost for all tools based on expression and LogCPM prediction results.*

***Supplementary Data 3:*** *Evaluation scores on all methods using all normalization methods on all benchmarks*

***Supplementary Data 4:*** *RGB images from different read count down samplings.*

***Supplementary Data 5:*** *RGB images and architecture detected by all methods using all normalization methods on all benchmarks.*

***Supplementary Data 6:*** *DEGs of the predictive seven segments.*

### Supplementary Figures

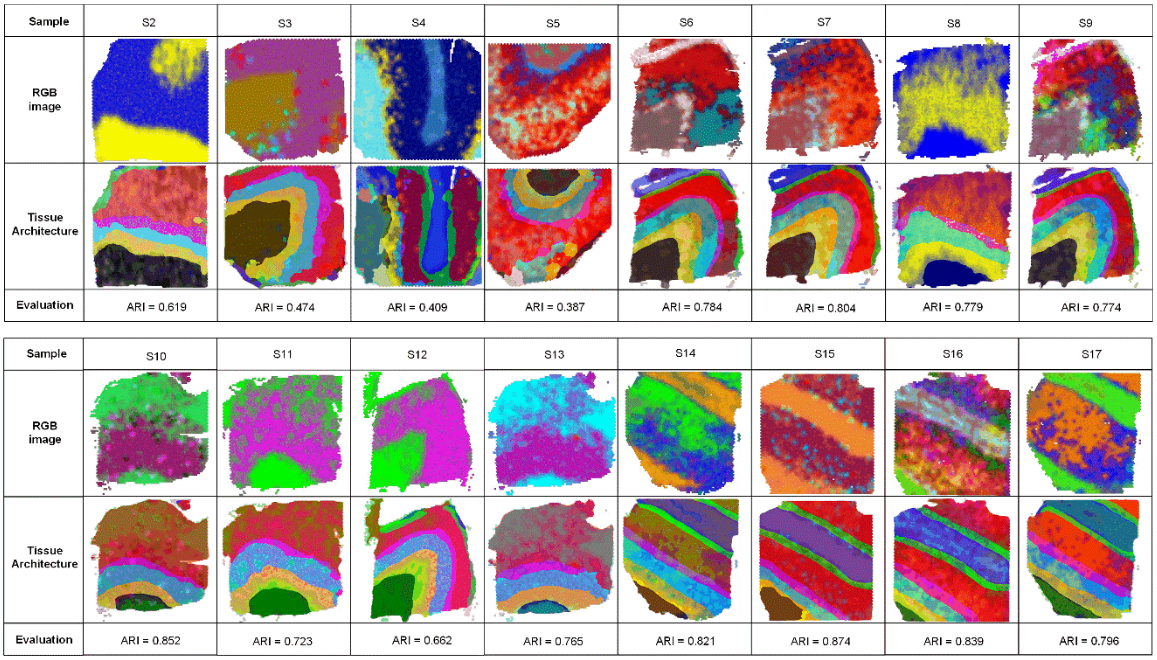


***Supplementary Fig. 1:*** *Abstract of RGB images & architecture detected by RESEPT based on LogCPM normalization.* The figures show RGB image, tissue architecture, and ARI identified by RESEPT for 16 samples (from S2 to S17).

| | | | | |
|---|---|---|---|---|
| **RGB image** | 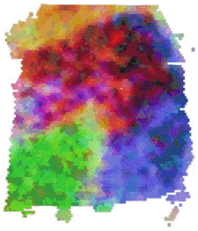 | 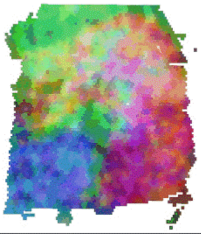 | 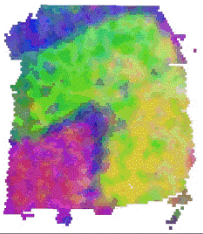 | 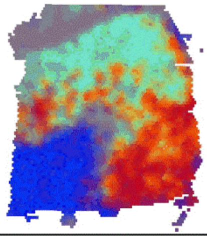 |
| **Effective read depth** | 500 | 1000 | 1500 | 2000 |
| **Evaluation** | ARI = 0.802 | ARI = 0.797 | ARI = 0.811 | ARI = 0.813 |
| **RGB image** | 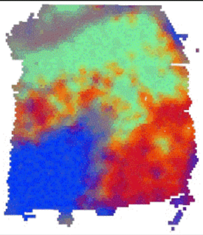 | 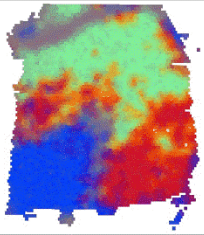 | 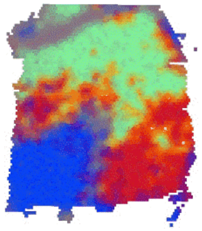 | 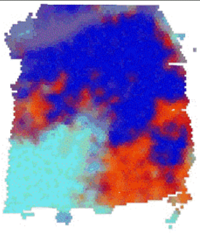 |
| **Effective read depth** | 2500 | 3000 | 3500 | 4000 |
| **Evaluation** | ARI = 0.811 | ARI = 0.815 | ARI = 0.807 | ARI = 0.807 |
| **RGB image** | 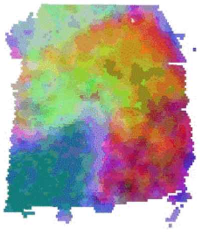 | 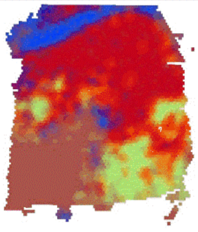 | 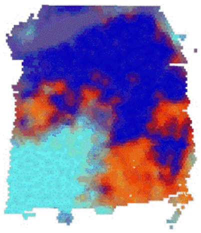 | 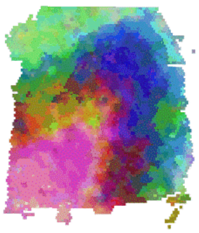 |
| **Effective read depth** | 4500 | 5000 | 5500 | 5908 |
| **Evaluation** | ARI = 0.804 | ARI = 0.811 | ARI = 0.816 | ARI = 0.818 |

***Supplementary Fig. 2:*** *RGB images of Sample S6 with different read depth.* The figure and ARI show the changes of RGB images across different effective read depths (from 500 to full read depth).

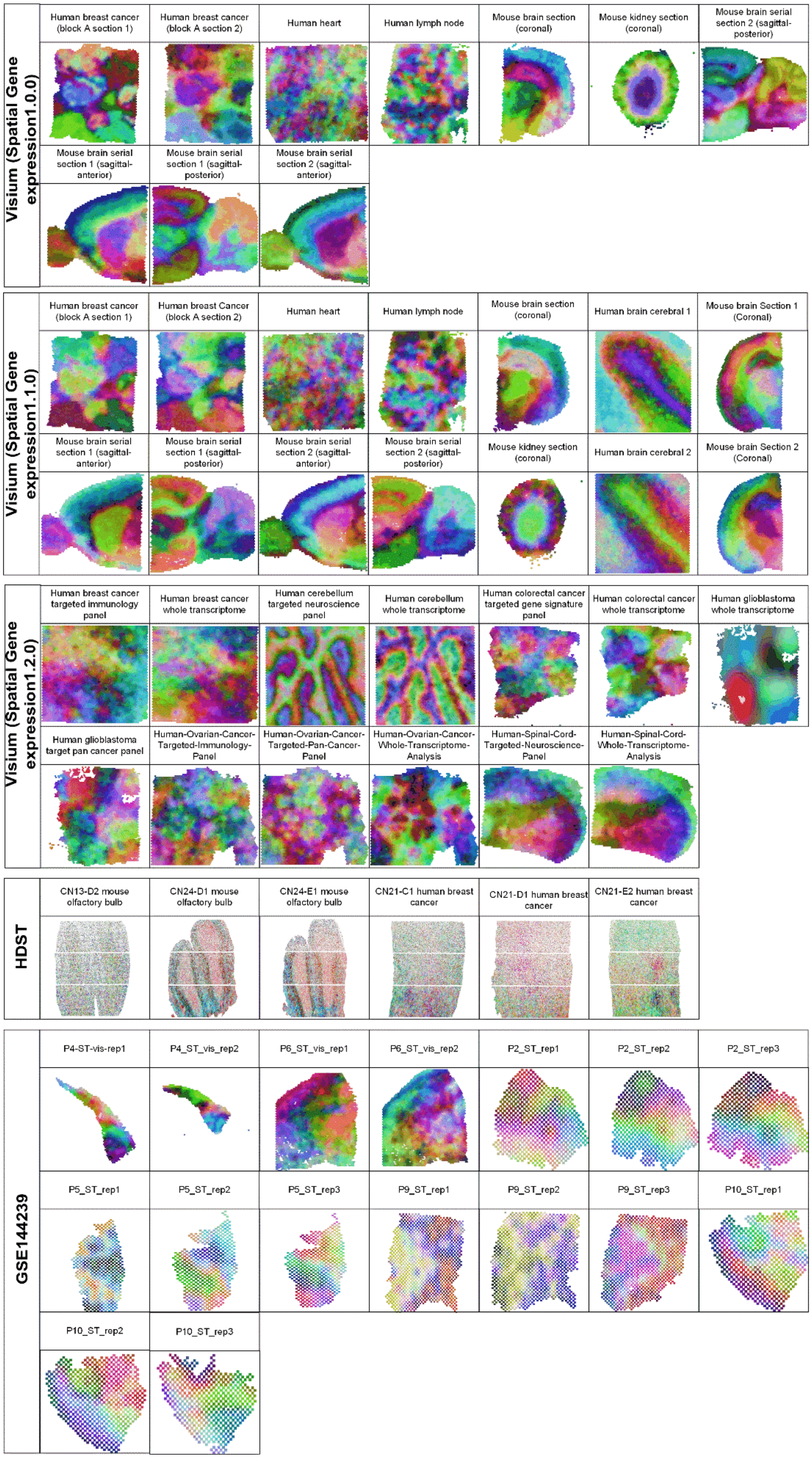
Visium (Spatial Gene expression1.0.0)
Human breast cancer (block A section 1)
Human breast cancer (block A section 2)
Human heart
Human lymph node
Mouse brain section (coronal)
Mouse kidney section (coronal)
Mouse brain serial section 2 (sagittal-posterior)
Mouse brain serial section 1 (sagittal-anterior)
Mouse brain serial section 1 (sagittal-posterior)
Mouse brain serial section 2 (sagittal-anterior)
Visium (Spatial Gene expression1.1.0)
Human breast cancer (block A section 1)
Human breast Cancer (block A section 2)
Human heart
Human lymph node
Mouse brain section (coronal)
Human brain cerebral 1
Mouse brain Section 1 (Coronal)
Mouse brain serial section 1 (sagittal-anterior)
Mouse brain serial section 1 (sagittal-posterior)
Mouse brain serial section 2 (sagittal-anterior)
Mouse brain serial section 2 (sagittal-posterior)
Mouse kidney section (coronal)
Human brain cerebral 2
Mouse brain Section 2 (Coronal)
Visium (Spatial Gene expression1.2.0)
Human breast cancer targeted immunology panel
Human breast cancer whole transcriptome
Human cerebellum targeted neuroscience panel
Human cerebellum whole transcriptome
Human colorectal cancer targeted gene signature panel
Human colorectal cancer whole transcriptome
Human glioblastoma whole transcriptome
Human glioblastoma target pan cancer panel
Human-Ovarian-Cancer-Targeted-Immunology-Panel
Human-Ovarian-Cancer-Targeted-Pan-Cancer-Panel
Human-Ovarian-Cancer-Whole-Transcriptome-Analysis
Human-Spinal-Cord-Targeted-Neuroscience-Panel
Human-Spinal-Cord-Whole-Transcriptome-Analysis
HDST
CN13-D2 mouse olfactory bulb
CN24-D1 mouse olfactory bulb
CN24-E1 mouse olfactory bulb
CN21-C1 human breast cancer
CN21-D1 human breast cancer
CN21-E2 human breast cancer
GSE144239
P4-ST-vis-rep1
P4_ST_vis_rep2
P6_ST_vis_rep1
P6_ST_vis_rep2
P2_ST_rep1
P2_ST_rep2
P2_ST_rep3
P5_ST_rep1
P5_ST_rep2
P5_ST_rep3
P9_ST_rep1
P9_ST_rep2
P9_ST_rep3
P10_ST_rep1
P10_ST_rep2
P10_ST_rep3

***Supplementary Fig. 3:*** *RGB images of data in 10x, ST, and HDST platforms without manual annotation.* The figure shows the application of RESEPT on other data via default parameter settings. For 10x Visium, ten samples from spatial gene expression 1.0.0 were used; 14 samples from spatial gene expression 1.1.0 were used; 13 samples from spatial gene expression 1.2.0 were used. For HDST data, six samples were used. For the GSE144239 dataset (squamous cell carcinoma), 4 Visium samples and 12 ST samples were used.

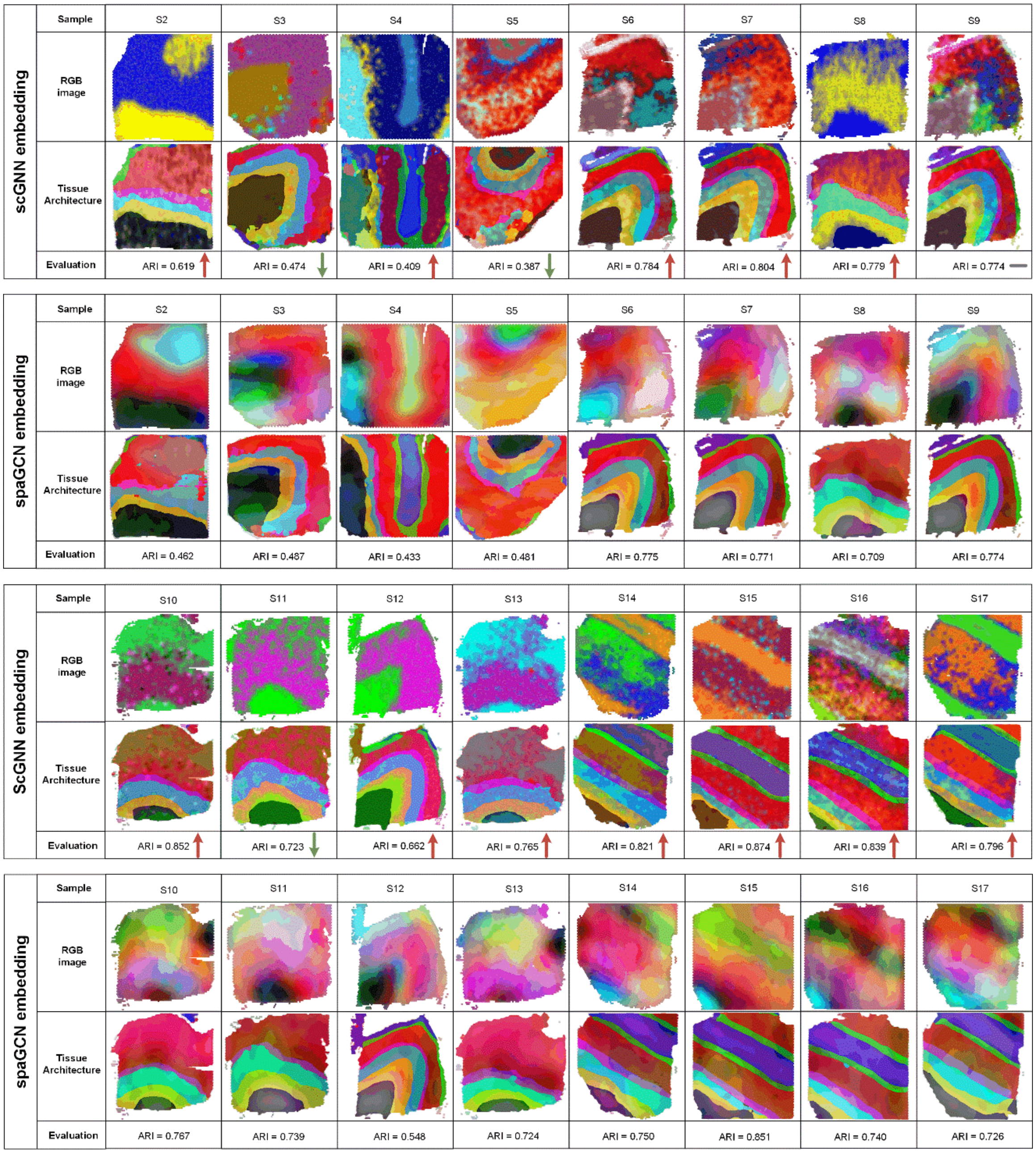


***Supplementary Fig. 4:*** *ARI metrics from scGNN and spaGCN on 16 samples.* The figure shows RGB image, tissue architecture, and ARI identified by RESEPT via using two different embedding methods, including scGNN and spaGCN. The red color arrow in the ARI panel indicates a higher ARI score for prediction results based on the scGNN embedding method, and the green color indicates a lower ARI score for the scGNN embedding method.

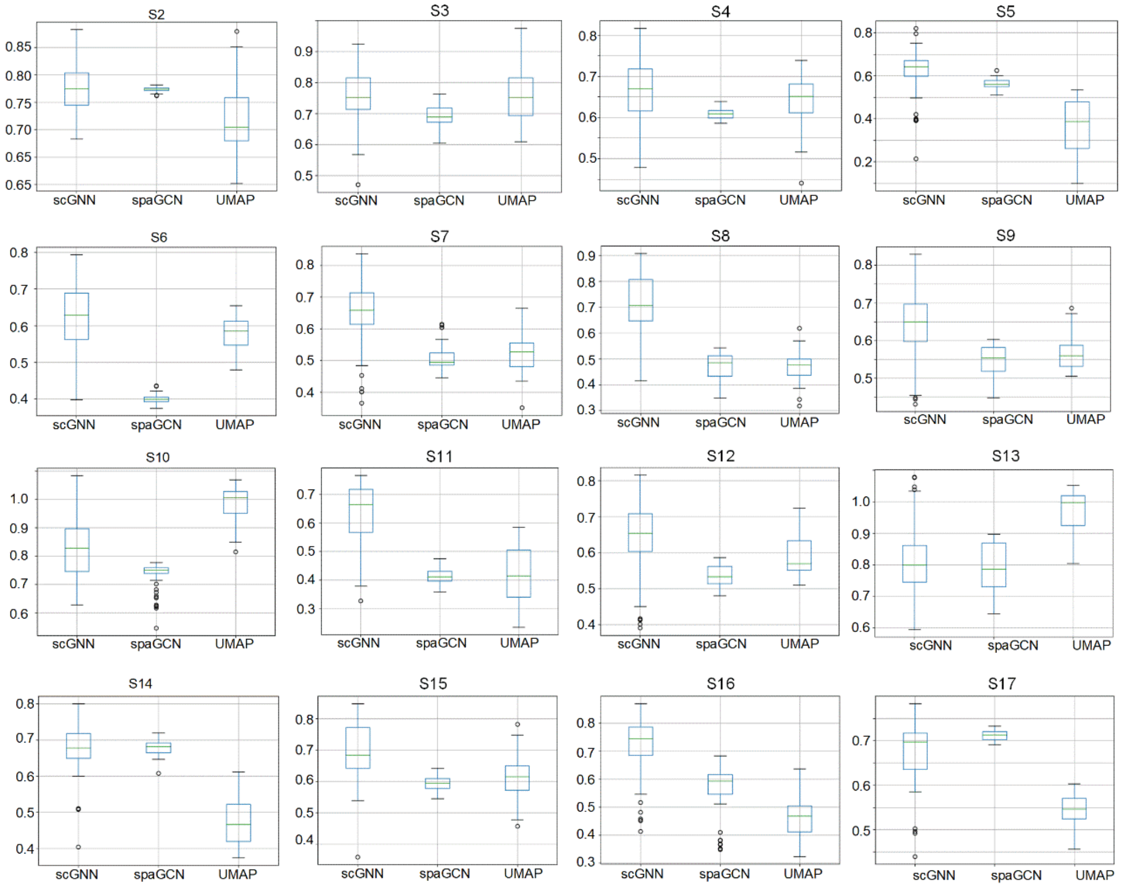


***Supplementary Fig. 5:*** *QA metrics from scGNN, spaGCN, and UMAP on 16 samples.* The 16 boxplots show Moran's I (MI) score for three embedding methods based on multiple parameter combinations. The X-axis represents three embedding methods, including scGNN, spaGCN, and UMAP. The Y-axis represents the MI score being calculated by multiple parameter settings.

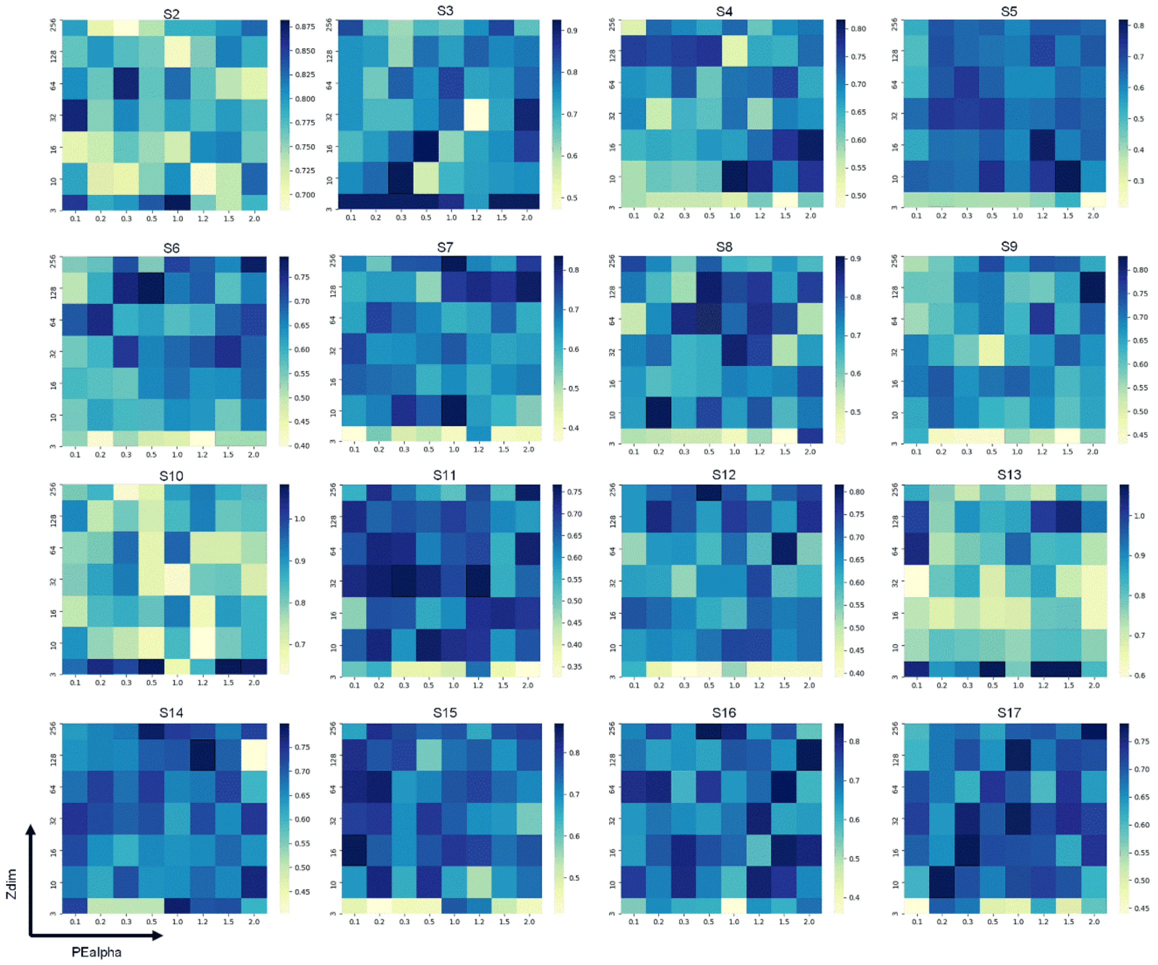


***Supplementary Fig. 6:*** *QA metrics for different parameter settings in 16 samples.* The heatmap shows the MI score calculated by two major parameters' combinations based on scGNN embedding. Zdim represents the number of principal components selected from 3, 10, 16, 32, 64, 128, and 256). PEalpha (described in Methods) of scGNN were selected from 0.1, 0.2, 0.3, 0.5, 1.0, 1.2, 1.5, and 2.0. The blue color indicates a higher MI score.

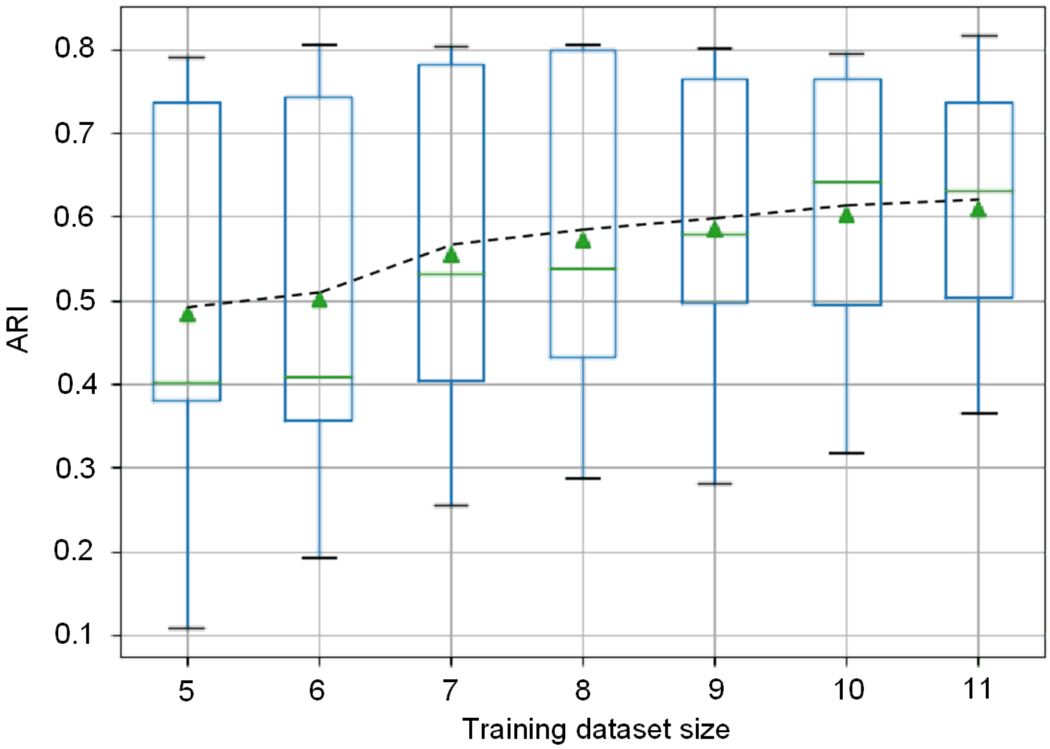


***Supplementary Fig. 7:*** *ARI error bar changes according to different training sets.* The boxplot shows ARI changes across different training datasets from 5 to 11 (select from S2, S3, S5, S6, S7, S8, S10, S12, S13, S16, and S17). The line presents mean ARI changes of five testing sets, including S4, S9, S11, S14, and S15.

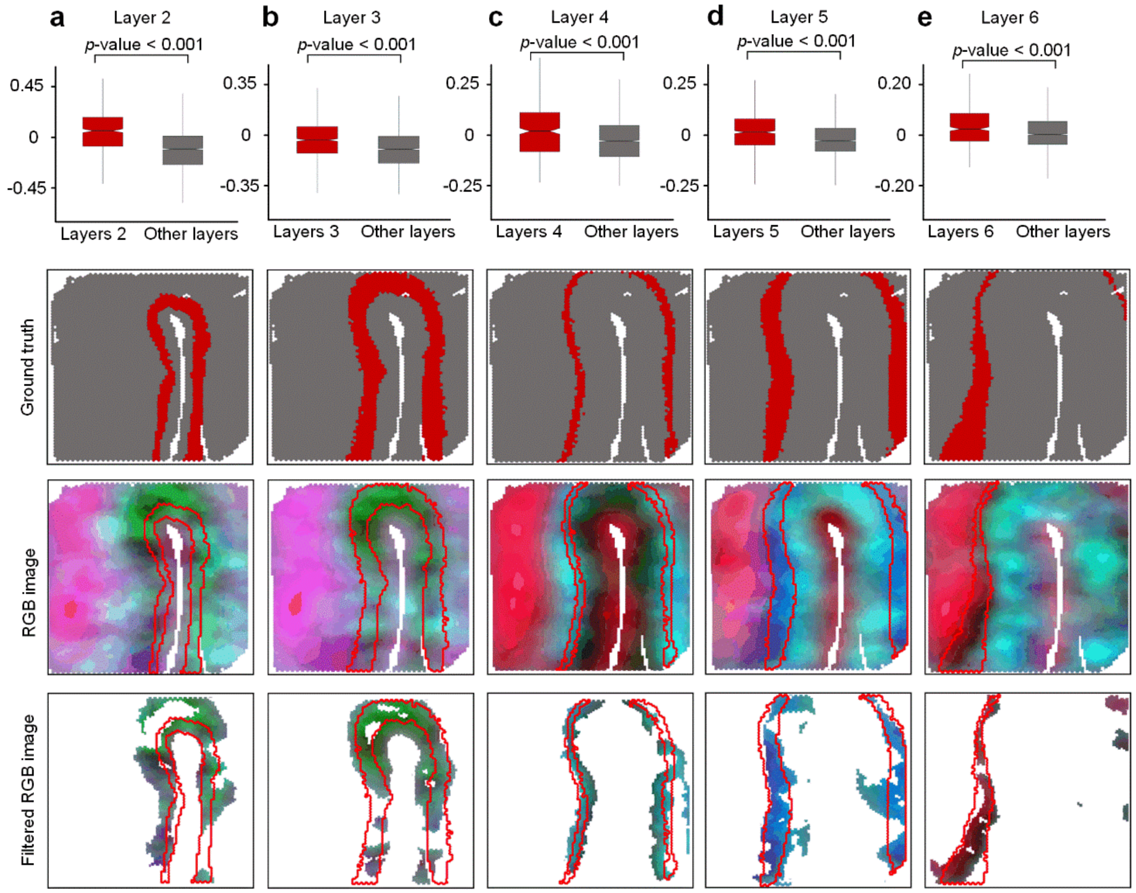


***Supplementary Fig. 8:*** *RGB image of other layer-specific markers on the AD sample S4 (update using non-normalization data).* (a) The box plot shows the module score of the cortical layers 2 from the AD sample (S4), in which the x-axis shows layer category, and the y-axis represents score value. The second figure shows the ground truth; the third figure shows an RGB image in which the red line shows ground truth; the fourth figure is reconstructed by filtering out unrelated colors. (b) The panel shows the layer 3 module score, ground truth, RGB image, and filtered RGB image. (c) The panel shows the layer 4 module score, ground truth, RGB image, and filtered RGB image. (d) The panel shows the layer 5 module score, ground truth, RGB image, and filtered RGB image. (d) The panel shows the layer 6 module score, ground truth, RGB image, and filtered RGB image.

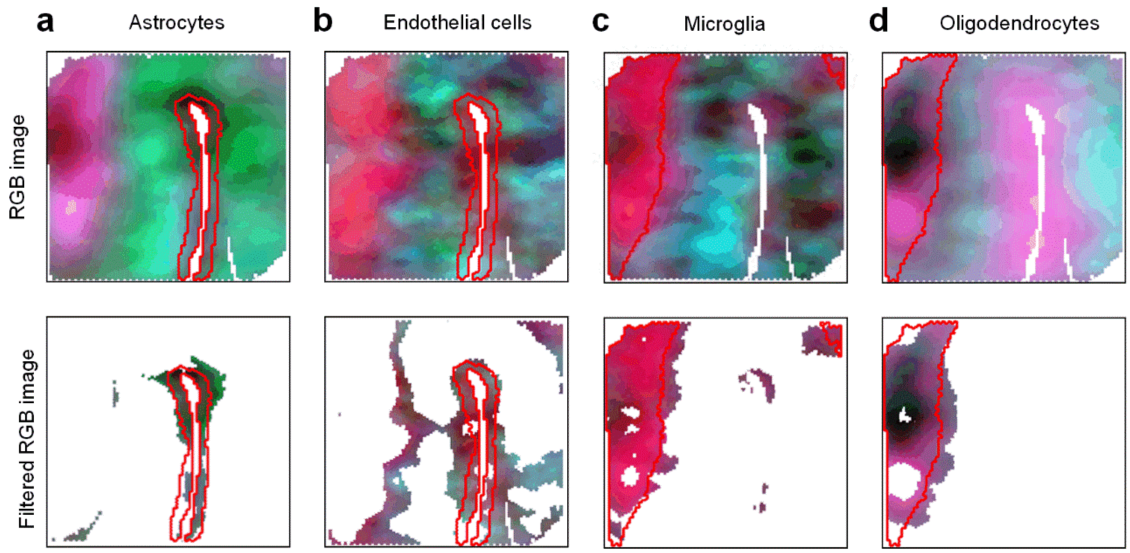


***Supplementary Fig. 9:*** *Other cell-specific markers on the AD sample S4.* (a) The RGB image and filtered RGB image show the astrocytes distribution reconstructed from the astrocytes marker genes. (b) The RGB image and filtered RGB image show the distribution of the endothelial cells reconstructed from the endothelial cell marker genes. (c) The RGB image and filtered RGB image show the distribution of microglia cells reconstructed from the microglia cell marker genes. (d) The RGB image and filtered RGB image show the oligodendrocytes distribution reconstructed from the oligodendrocytes marker genes.

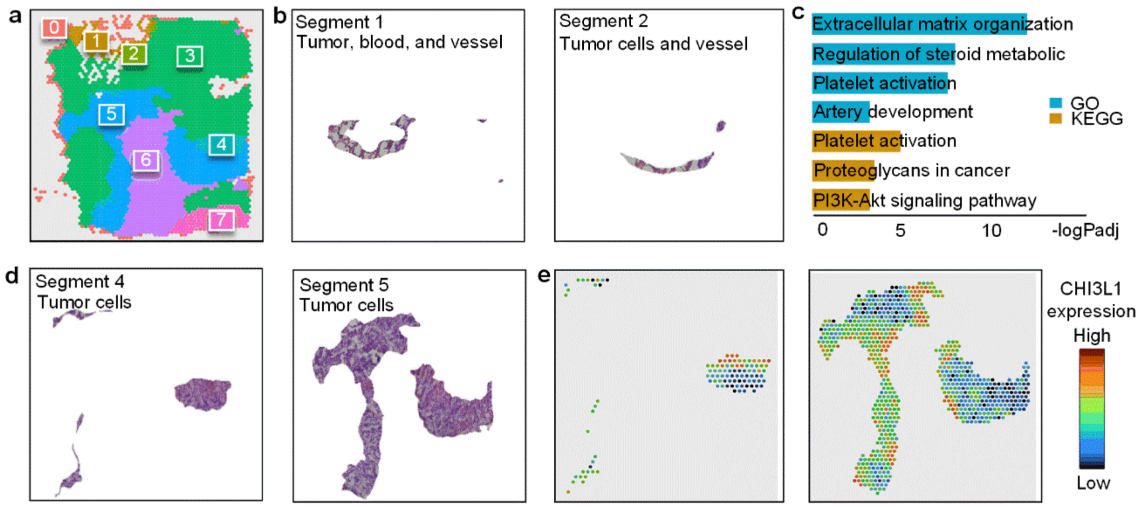


***Supplementary Fig. 10:** Other 4 section morphological features and transcriptional features on Glioblastoma sample S1.* (a) The figure shows the RESEPT prediction results, including seven segments. (b) Two figures show the HE image cropped based on segmentation results, including Segments 1 and 2. Our physiologist found Segment 1 and 2 tumors, blood cells, and vessels based on morphological features. (c) The pathway enrichment result from GO and KEGG shows platelet activation for segment1 and segment2. Furthermore, regulation of steroid metabolic, proteoglycans regulation, and P13K-Akt signaling pathway was reported to associate glioblastoma in previous studies. (d) Two figures show the HE image cropped based on segmentation results, including segments 4 and 5, containing many tumor cells. (e) Heatmap shows glioblastoma marker CHI3L1 expression on segments 4 and 5.

### Supplementary Tables

***Supplementary Table 1:*** *Summary of datasets used in the study.* The table displays all data and its meta-information using in this study, including benchmarking and training dataset (S1- S17) and other application datasets. The Training column indicates whether the data were used for training and benchmarking. The Annotation column indicates whether data were fully annotated. The Tech column indicates a spatially resolved transcriptomics platform. The Tissue column cat1 and the Tissue cat2 column indicate health conditions and tissue information. The Species column indicates the species information. The Disease status column indicates whether the data belong to disease tissue. The last column indicates the number of samples in each data. Sixteen samples (S2- S17) were from Training set and Test data in the first row. Case study (S1) was in the sixth row. *Abbreviations: Spatial Transcriptomics (ST), high-definition spatial transcriptomics, Alzheimer's disease (AD).*

| Training | Annotation | Tech | Tissue cat1 | Tissue cat2 | Species | Disease status | # of samples |
|---|---|---|---|---|---|---|---|
| Training set | Labeled | Visium | Health | Brain | Human | No | 12 |
| Test data | Labeled | Visium | Health | Brain | Human | No | 2 |
| Test data | Labeled | Visium | AD | Brain | Human | Yes | 2 |
| Application | Unlabeled | Visium | Tumor | Breast | Human | Yes | 5 |
| Application | Unlabeled | Visium | Tumor | Colon | Human | Yes | 1 |
| Application | Unlabeled | Visium | Tumor | Glioblastoma | Human | Yes | 1 |
| Application | Unlabeled | Visium | Tumor | Skin cancer | Human | Yes | 4 |
| Application | Unlabeled | ST | Tumor | Skin cancer | Human | Yes | 12 |
| Application | Labeled | HDST | Health | Brain | Mouse | No | 3 |
| Application | Labeled | HDST | Tumor | Breast | Human | Yes | 3 |
| Application | Unlabeled | Visium | Health | Brain | Mouse | No | 16 |
| Application | Unlabeled | Visium | Health | Heart | Mouse | No | 2 |
| Application | Unlabeled | Visium | Health | Lymph node | Human | No | 2 |
| Application | Unlabeled | Visium | Health | Kidney | Mouse | No | 2 |
| Application | Unlabeled | Visium | Health | Spin cord | Human | No | 2 |
| Application | Unlabeled | Visium | Tumor | Glioblastoma | Human | Yes | 1 |
| Application | Unlabeled | Visium | Tumor | Ovarian | Human | Yes | 3 |
| Application | Unlabeled | Visium | Tumor | Colon | Human | Yes | 1 |
| Application | Unlabeled | Visium | Tumor | Breast | Human | Yes | 1 |

***Supplementary Table 2:*** *Marker genes in AD.* The table shows the signature gene summarized from publications. The first table lists markers identified from Maynard's study, indicating layer-specific markers for the human brain dorsolateral prefrontal cortex. The second table lists cell-type-specific markers from scREAD database. The third table lists Amyloid-beta-associated genes from Chen's study from the mouse.

| By layers | |
|---|---|
| Layer 1 | RELN |
| Layer 2 | C1QL2, RASGRF2, CARTPT, WFS1, HPCAL1 |
| Layer 3 | CARTPT, MFGE8, PRSS12, SV2C, HPCAL1 |
| Layer 4 | RORB, PDYN, CHRNA3, KCNIP2, KCNIP1, CYP39A1 |
| Layer 5 | PCP4, HTR2C, VAT1L, ETV1, CRIM1, NEFH, OPN3, OMA1, FOXO1, S100A10, LIX1, LDB2, CNTN6, SYT9 |
| Layer 6 | OPRK1, SYNPR, ANXA1, PCDH17, TLE4, SYT6, TH, FOXP2, LXN, AKR1C2, AKR1C3, NR4A3, SEMA3C, SYT10, NPY2R, PPP1R1B |
| White matter | MBP, MOBP |
| **By cell type** | |
| Astrocytes | GFAP, EAAT1, AQP4, GJA1, SLC1A2, FGFR3, NKAIN4, AGT, PLXNB1, SLC1A3 |
| Endothelial cells | CLDN5, ITM2A, VWF, BMX, CDH5 |
| Excitatory neurons | SLC17A6, SLC17A7, NRGN, CAMK2A, SATB2 |
| Inhibitory neurons | SLC32A1, GAD1, GAD2 |
| Microglia | P2RY12, CSF1R, CX3CR1, C3, HEXB, AIF-1, TMEM119 |
| Oligodendrocyte | OLIG2, MBP, MOBP, PLP1, MYRF, ERMN, MAG |
| Oligodendrocyte precursor cells | VCAN, SOX8, SOX10 |
| **Abeta_gene_mouse** | |
| Abeta | CTSH, FCGR2A, FCGR2B, LY86, LYZ, TYROBP, AXL, C1QA, C1QB, C1QC, CTSS, CYBA, GRN, HEXB, LAPTM5, OLFML3, CLU, CST3, GFAP, IGFBP5 |

**Supplementary Data (individual files)**

***Supplementary Data 1:*** *ARI and tuned parameter usage tools for generating 7 clusters.* The first table shows the ARI score of predicting tissue architecture for seven tools. The second table (parameter matching to the first table) shows the parameter usage for generating seven clusters. NA in the first and second table means that the software cannot find seven clusters via different parameter combinations.

***Supplementary Data 2:*** *Time and memory cost for all tools based on expression and LogCPM prediction results.*

The table shows the time efficiency (unit: Second) and memory (unit: GB) cost of seven tools based on LogCPM and raw expression matrix.

***Supplementary Data 3:*** *Evaluation scores on all methods using all normalization methods on all benchmarks*

The table shows the four matrices evaluation score, including ARI, RI, FM, and AMI. For each evaluation score, 17 samples were used for performing tissue architecture identification based on raw count matrix and seven normalization method, including TPM, TMM, scTransform, scran, RPKM, LogCPM, expression, and DESeq2. Seven tools were used for benchmarking, including STUtility, spaGCN, Seurat, RESEPT, Giotto, stlearn, and BayesSpace. Stlearn was only used based on raw expression data due to SME normalization method requiring raw express data as input. NA in the score represents an error to perform the software.

***Supplementary Data 4:*** *RGB images from different read count down samplings.*

The data are included in four folders, including S5_scGNN, S5_spaGCN, S6_scGNN, and S6_spaGCN. S5_scGNN folder contains 15 images generating from 500 effective read depth to full read depth based on Sample S5 and scGNN embedding method. S5_scGNN folder contains 17 images. S5_spaGCN contains 17 images. S6_scGNN folder contains 12 images generating from 500 effective read depth to full read depth based on Sample S6 and scGNN embedding method. S6_spaGCN folder contains 12 images.

***Supplementary Data 5:*** *RGB images and architecture detected by all methods using all normalization methods on all benchmarks.*

The data contains two folders, including 'Other six tools' and 'RESEPT.' In the 'Other six tools' folder, eight subfolders indicate eight input methods (DESeq2, TMM, TPM, expression, LogCPM, scran, sctransform, and RPKM) and contain visualization results generated from 16 samples and six benchmarking tools. The ‘RESEPT’ folder contains ‘RGB image’ subfolder, providing RGN images generating from seven input methods (DESeq2, TMM, TPM, expression, scran, sctransform, and RPKM). The other subfolder (‘Tissue architecture’)provides image segmentation results for visualizing tissue architecture.

***Supplementary Data 6:*** *DEGs for predictive seven segments.*

The table shows the differentially expressed gene analysis for seven predictive segments from RESEPT. The P_val column indicates p-value calculated from the Wilcoxon Rank Sum test. The Avg_logFC column indicates average log-foldchange. The pct.1 column indicates the number of percentages for a gene expressed in group 1 spots. The pct.1 column indicates the number of percentages for a gene expressed in group 2 spots. The cluster column indicates the segments category. The gene column indicates the gene symbol name.
