## Supplementary figures and images for "Define and visualize pathological architectures of human tissues from spatially resolved transcriptomics using deep learning"

### S2_6_tools_DESeq2.jpeg

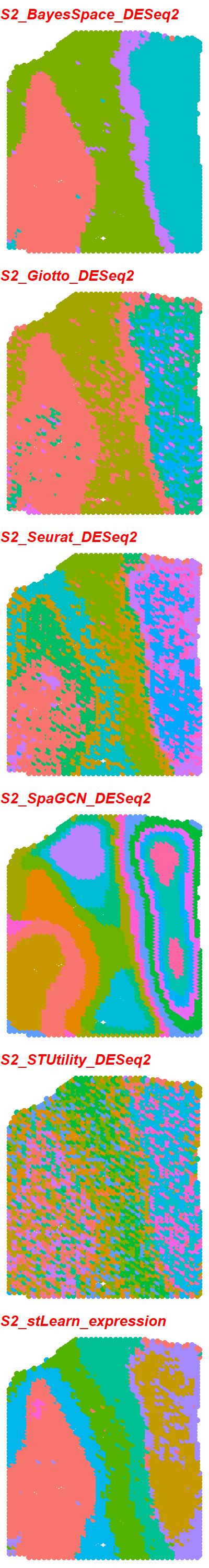

### S2_6_tools_expression.jpeg

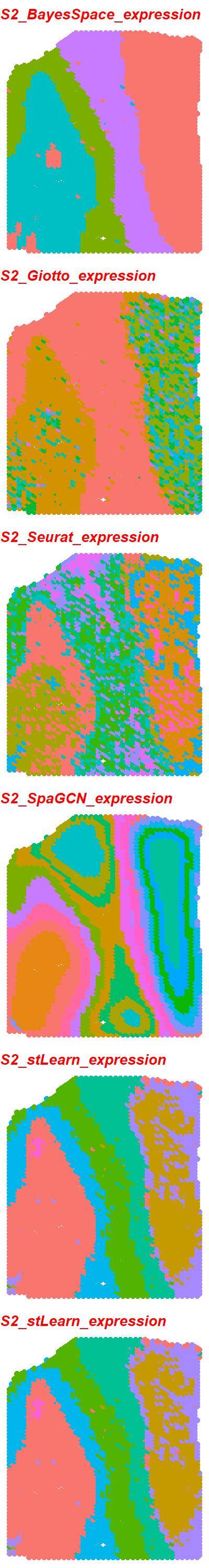

### S2_6_tools_LogCPM.jpeg

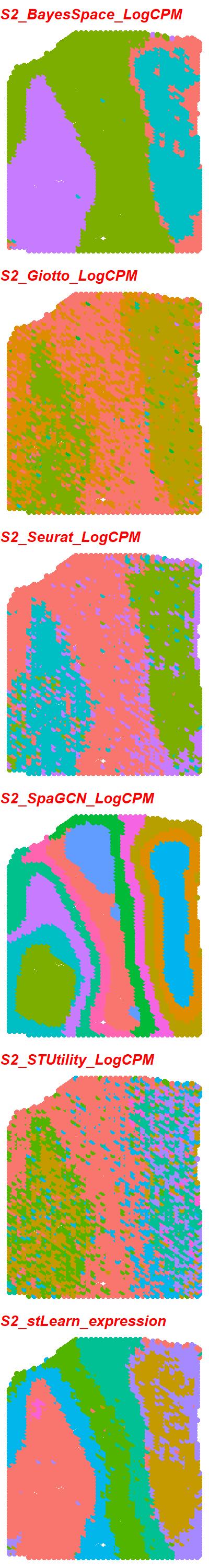

### S3_6_tools_DESeq2.jpeg

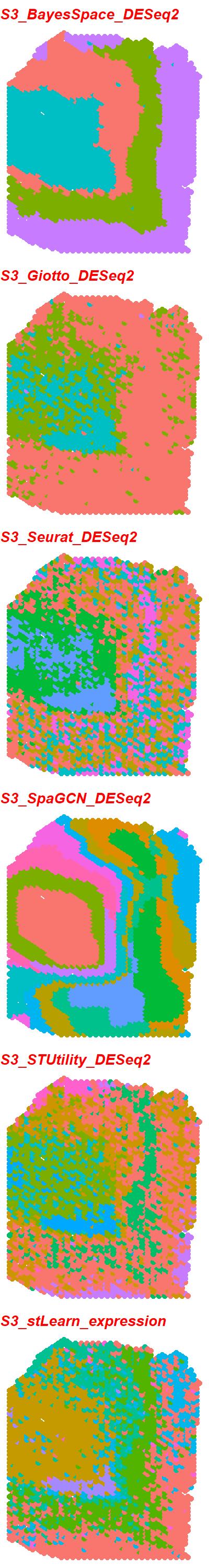

### S3_6_tools_expression.jpeg

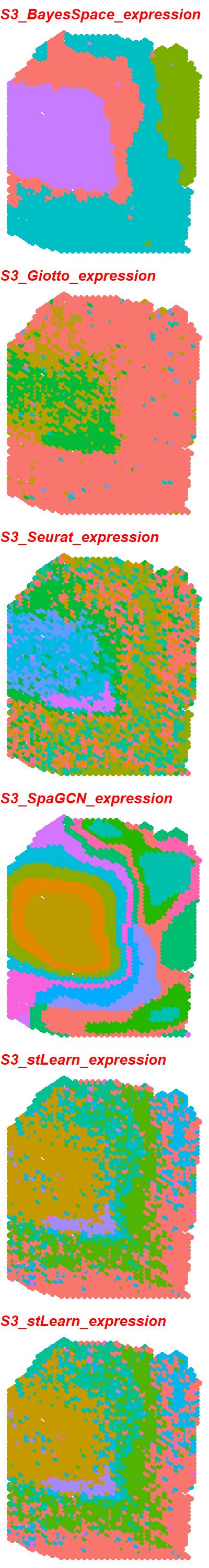

### S3_6_tools_LogCPM.jpeg

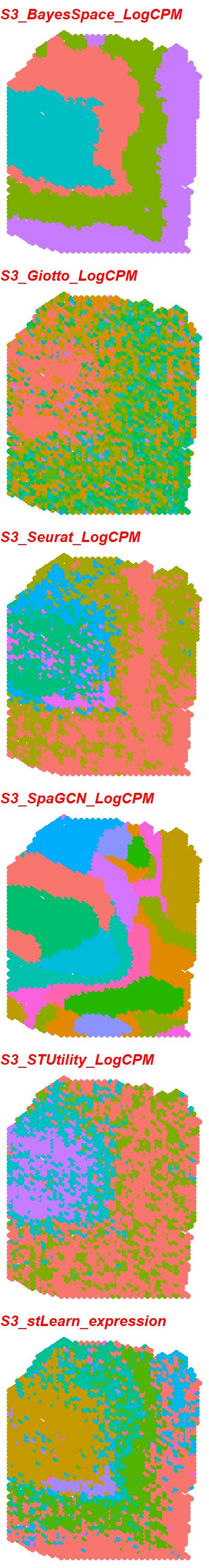

### S4_6_tools_DESeq2.jpeg

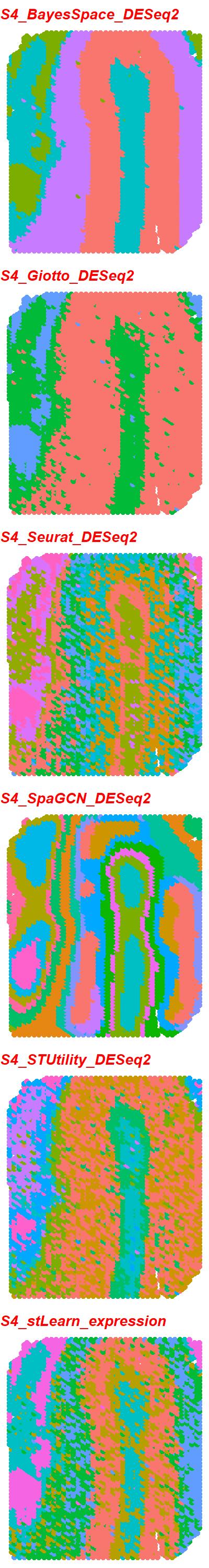

### S4_6_tools_expression.jpeg

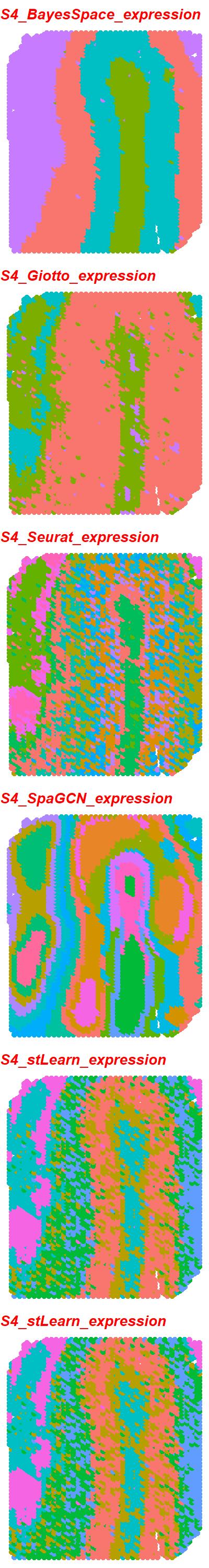

### S5_6_tools_DESeq2.jpeg

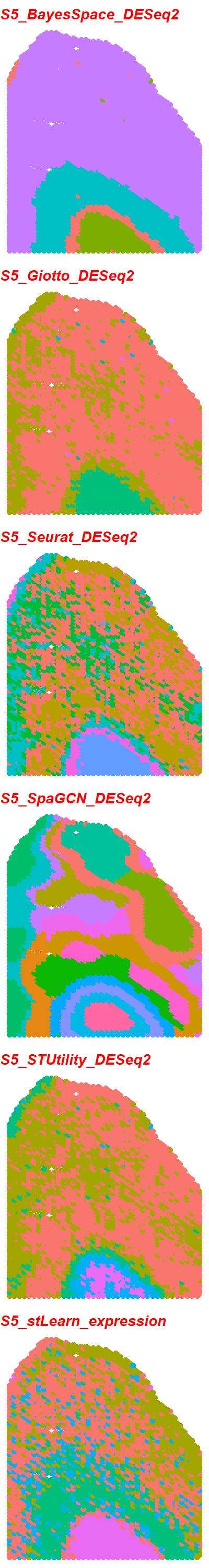

### S5_6_tools_expression.jpeg

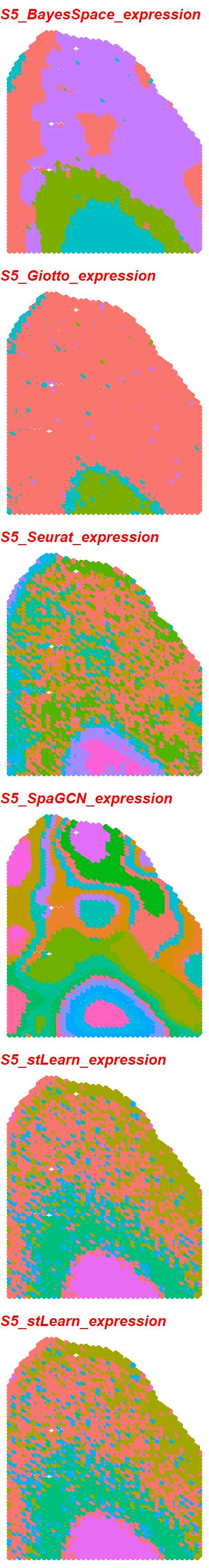

### S5_scGNN_logcpm_PEalpha0.1_zdim3_downsampling1000.png

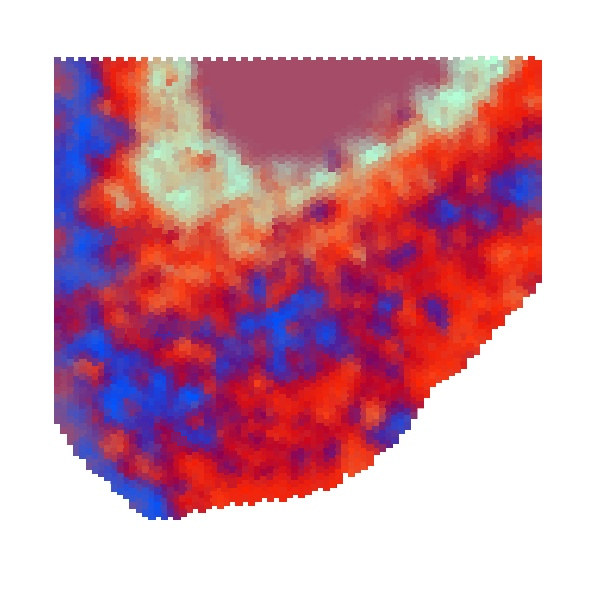

### S5_scGNN_logcpm_PEalpha0.1_zdim3_downsampling1500.png

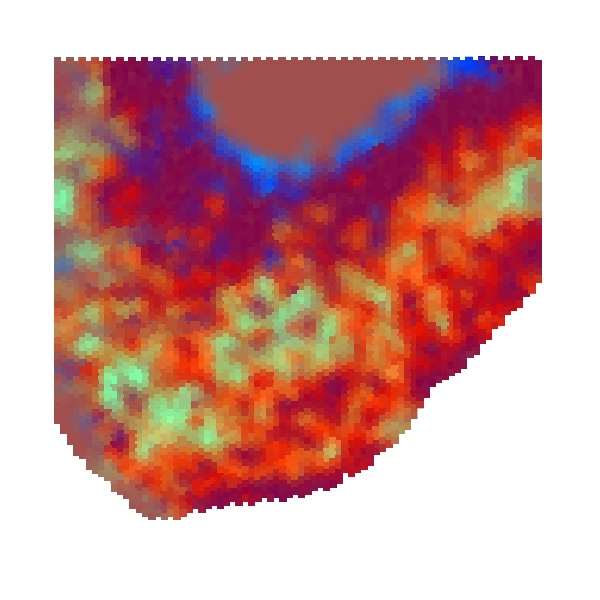

### S5_scGNN_logcpm_PEalpha0.1_zdim3_downsampling5500.png

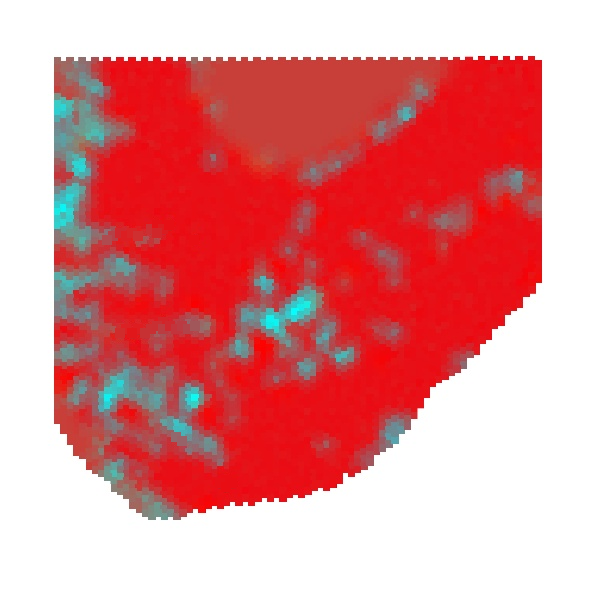

### S5_scGNN_logcpm_PEalpha0.2_zdim3_downsampling6500.png

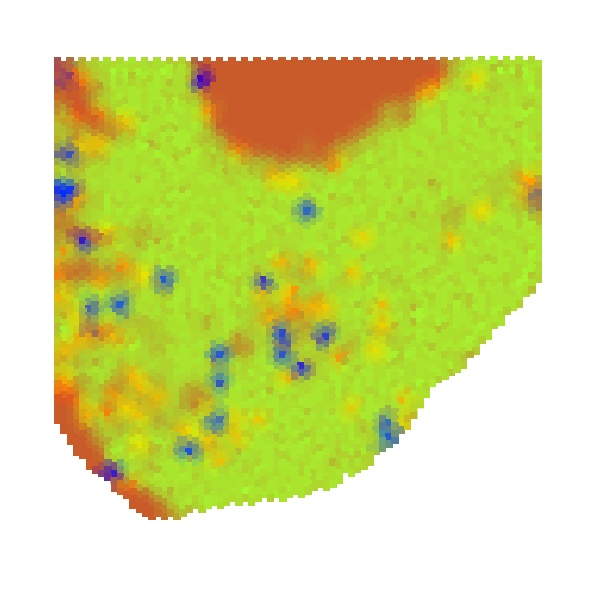

### S5_scGNN_logcpm_PEalpha0.2_zdim3_downsampling7000.png

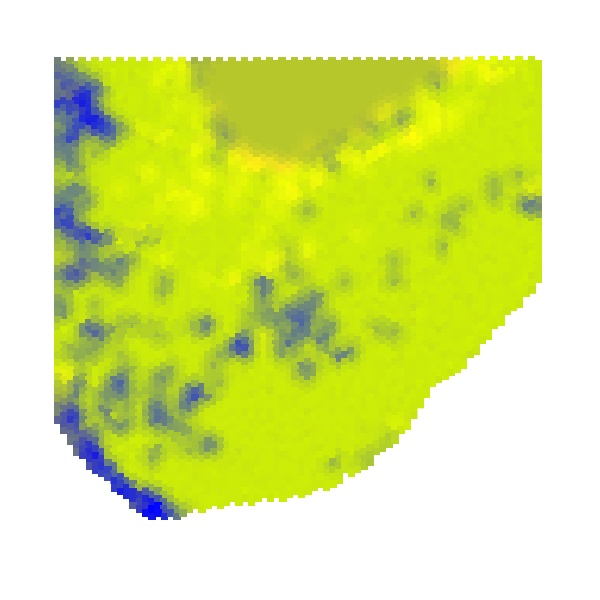

### S5_scGNN_logcpm_PEalpha0.2_zdim256_downsampling500.png

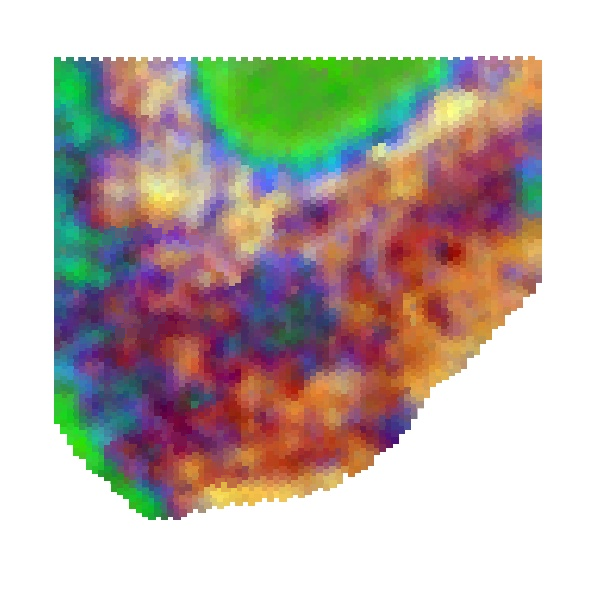

### S5_scGNN_logcpm_PEalpha0.3_zdim3_downsampling2500.png

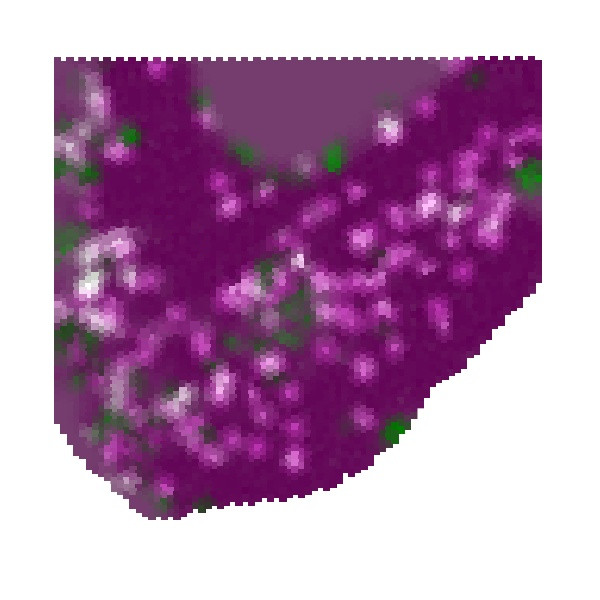

### S5_scGNN_logcpm_PEalpha1.0_zdim3_downsampling2000.png

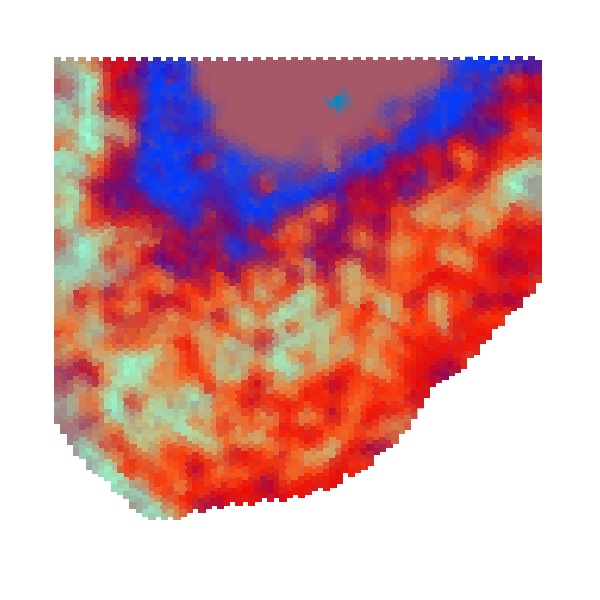

### S5_scGNN_logcpm_PEalpha1.0_zdim3_downsampling3000.png

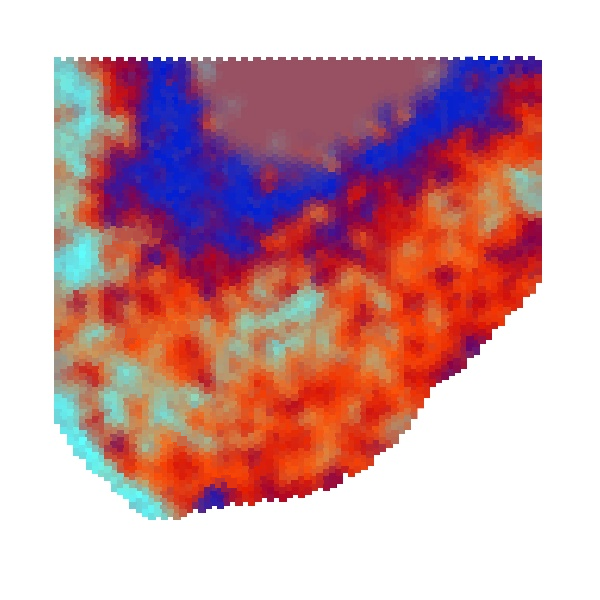

### S5_scGNN_logcpm_PEalpha1.0_zdim3_downsampling4000.png

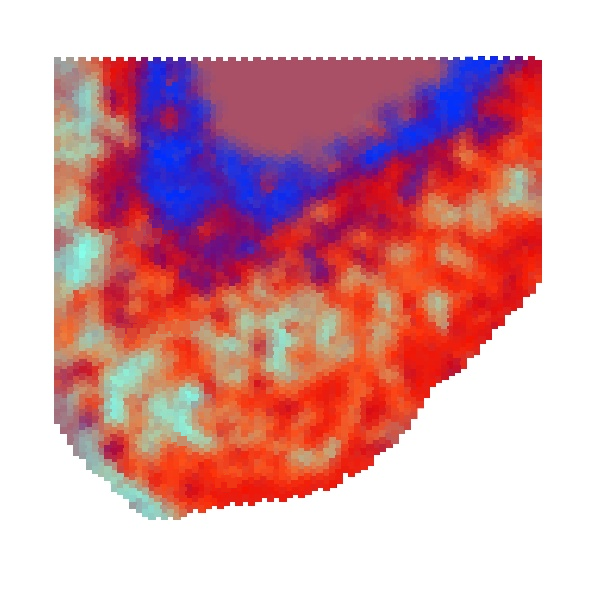

### S5_scGNN_logcpm_PEalpha1.2_zdim3_downsampling3500.png

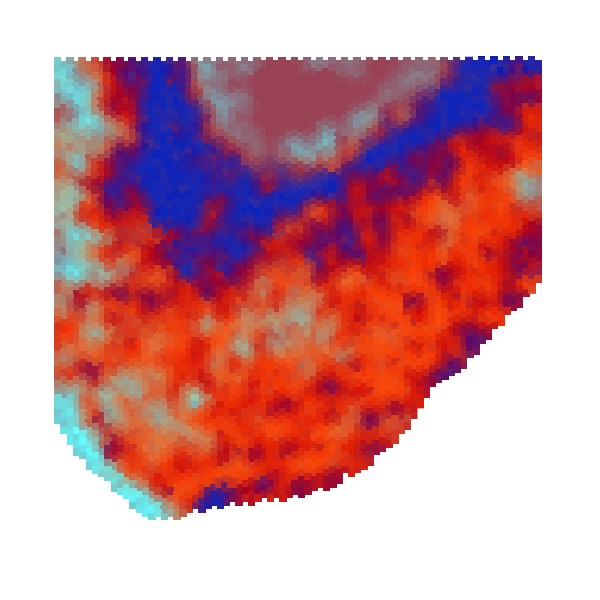

### S5_scGNN_logcpm_PEalpha1.2_zdim3_downsampling4500.png

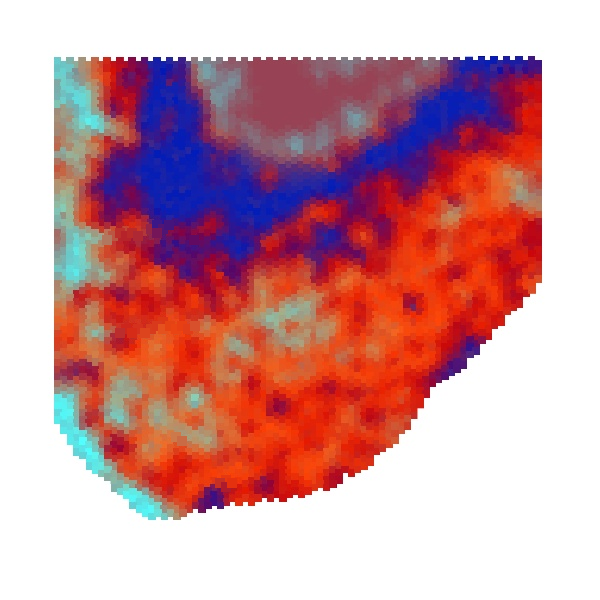

### S5_scGNN_logcpm_PEalpha1.2_zdim128_downsampling6000.png

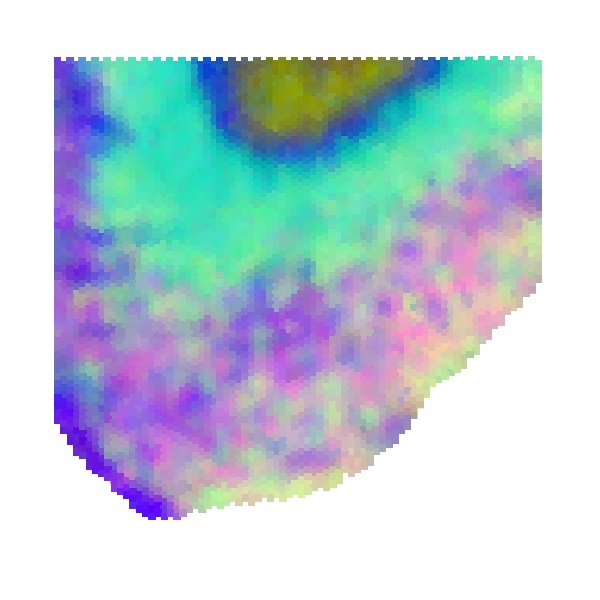

### S5_scGNN_logcpm_PEalpha1.5_zdim3_downsampling5000.png

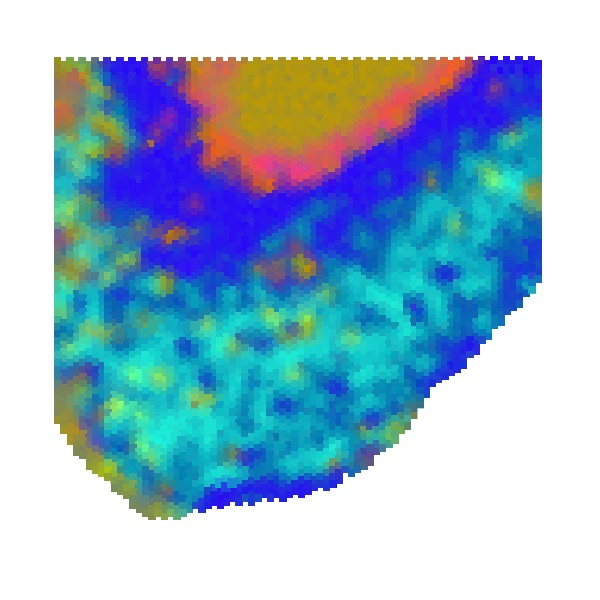

### S5_scGNN_logcpm_PEalpha1.5_zdim3_full.png

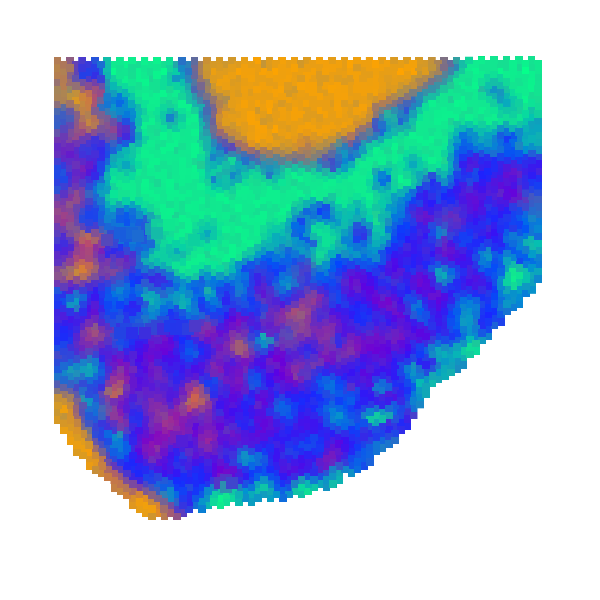

### S5_scGNN_logcpm_PEalpha2.0_zdim3_downsampling7500.png

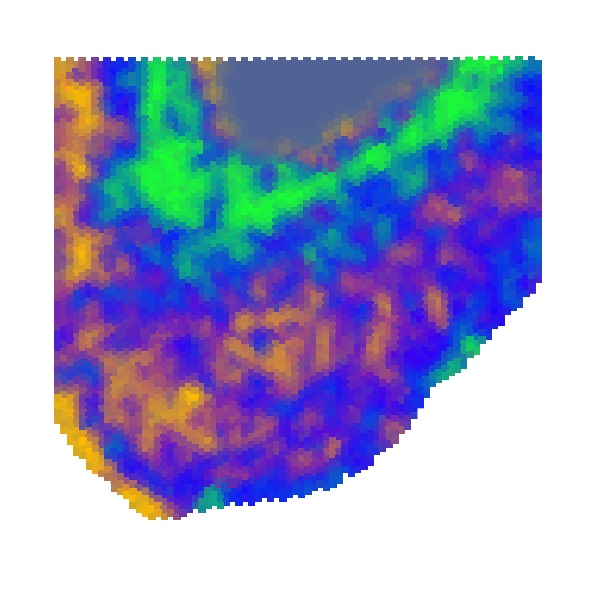

### S5_scGNN_logcpm_PEalpha2.0_zdim3_downsampling8000.png

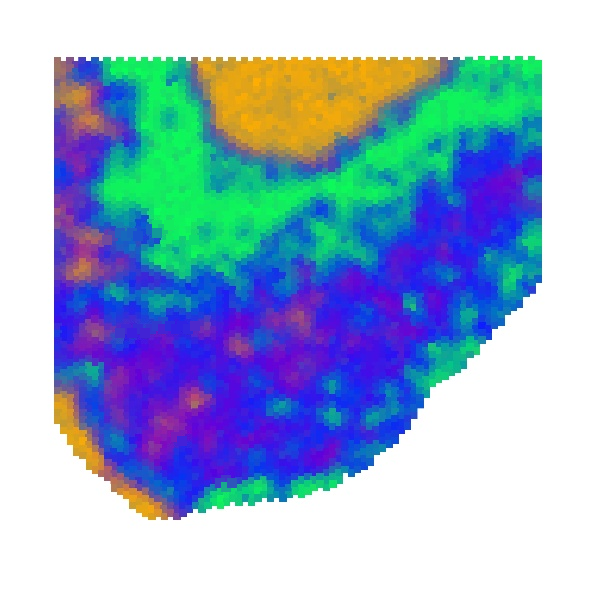

### S5_spaGCN_logcpm_pca32_res0.2_downsampling3500.png

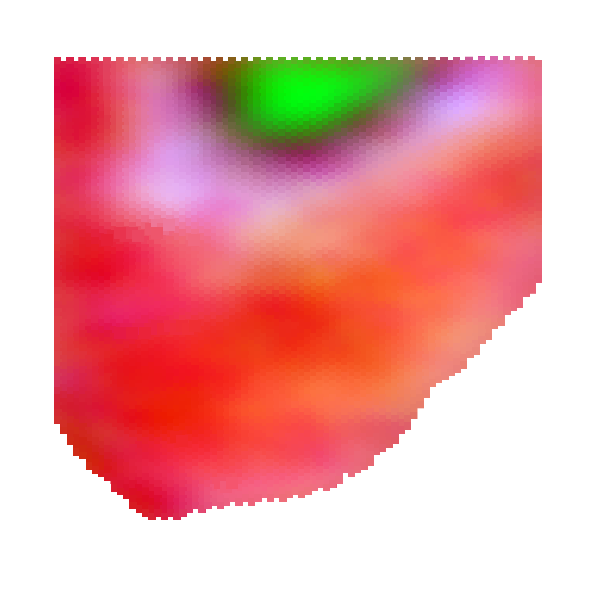

### S5_spaGCN_logcpm_pca32_res0.2_downsampling4000.png

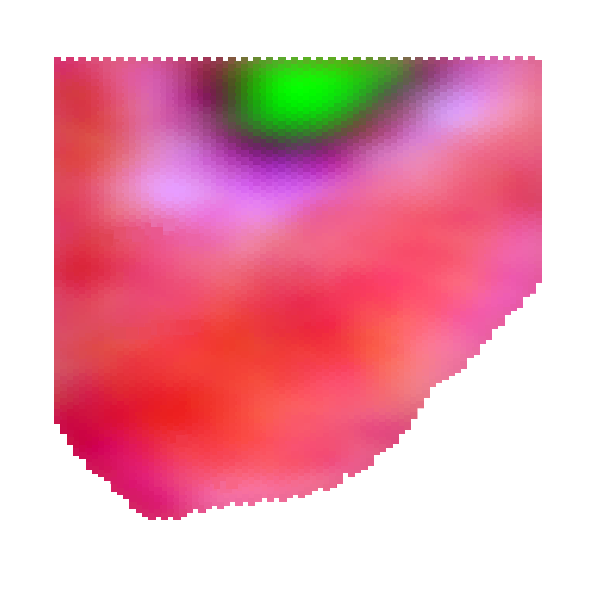

### S5_spaGCN_logcpm_pca32_res0.4_downsampling2500.png

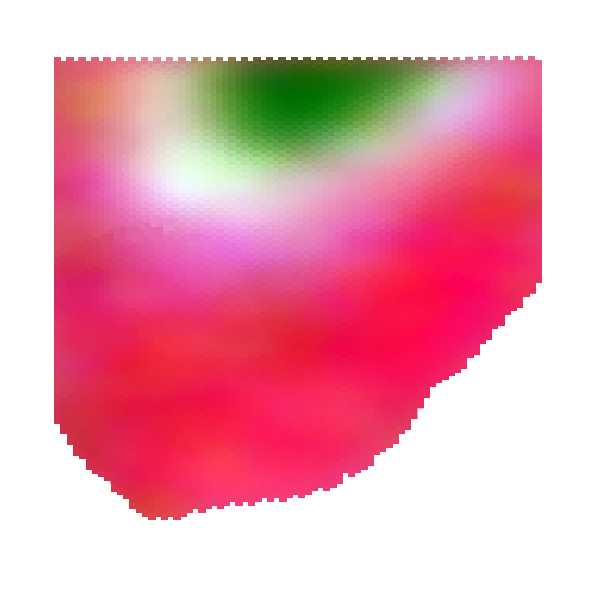

### S5_spaGCN_logcpm_pca32_res0.5_downsampling1500.png

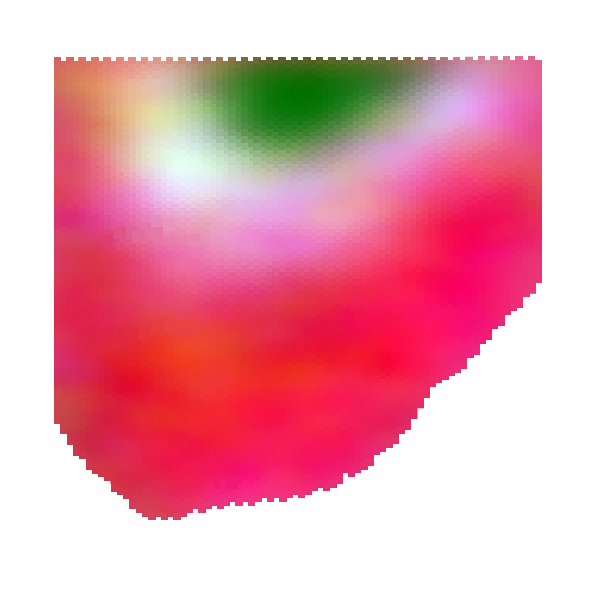

### S5_spaGCN_logcpm_pca32_res0.25_downsampling5500.png

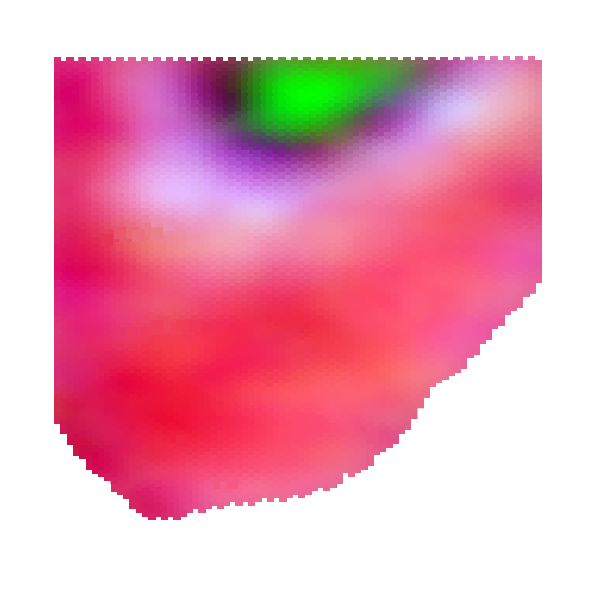

### S5_spaGCN_logcpm_pca32_res0.45_downsampling2000.png

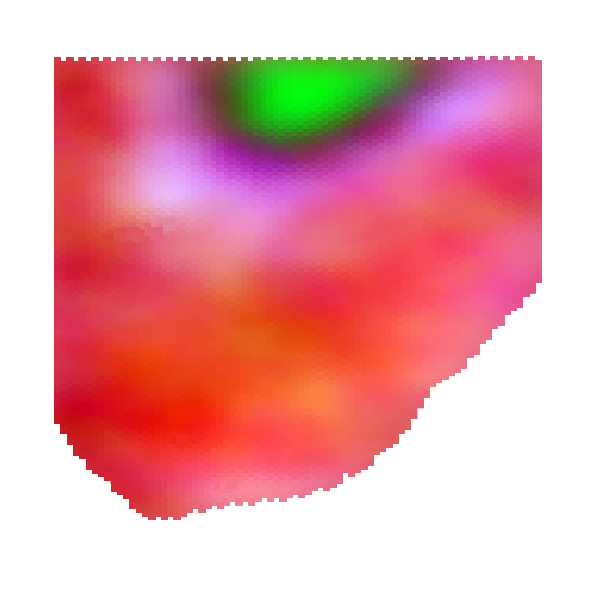

### S5_spaGCN_logcpm_pca32_res0.55_full.png

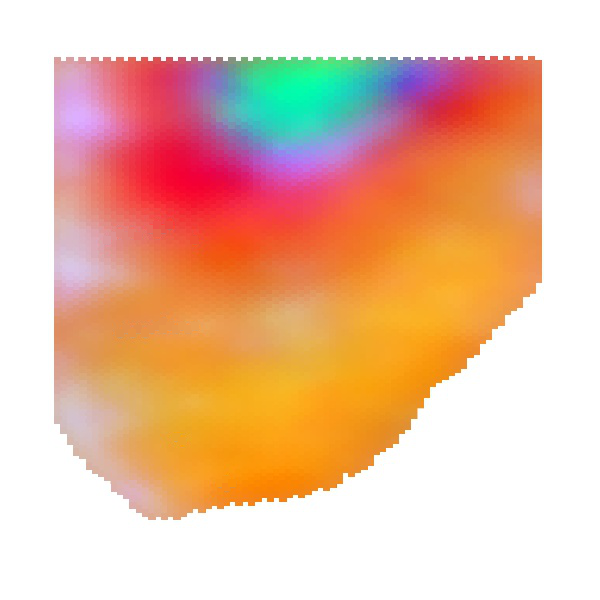

### S5_spaGCN_logcpm_pca50_res0.35_downsampling500.png

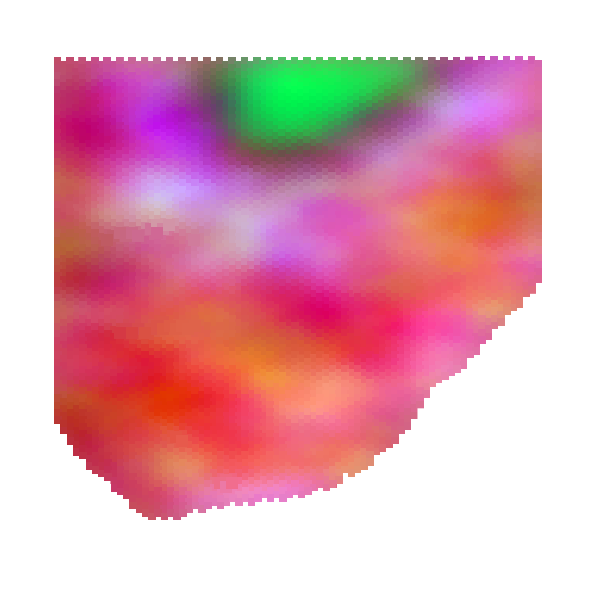

### S5_spaGCN_logcpm_pca64_res0.4_downsampling1000.png

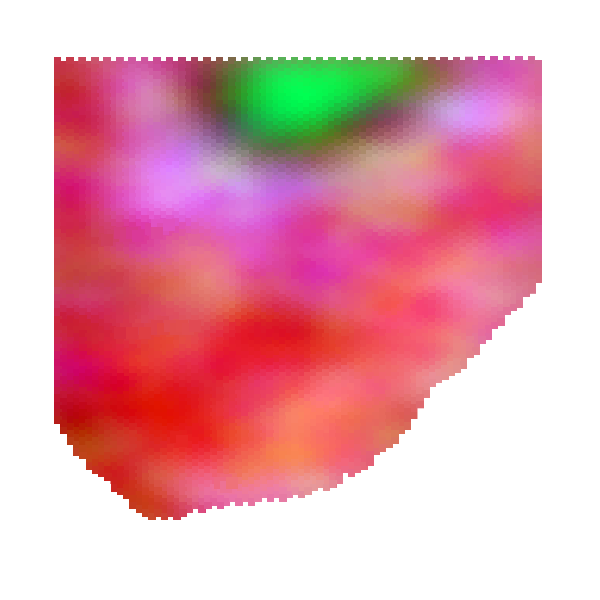

### S5_spaGCN_logcpm_pca64_res0.35_downsampling7000.png

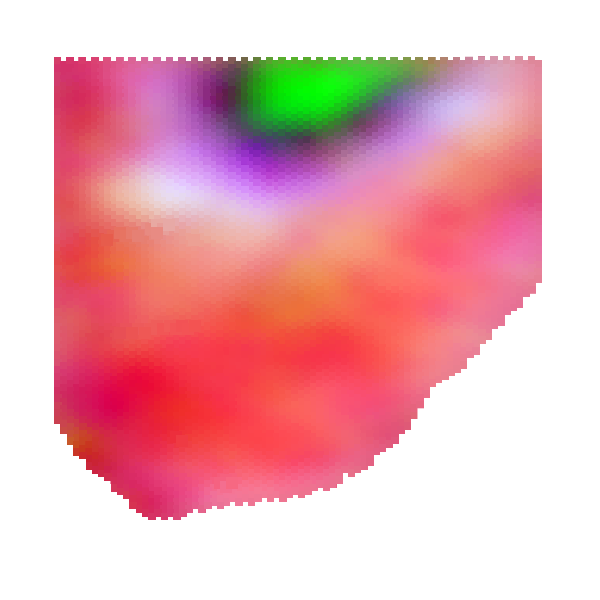

### S5_spaGCN_logcpm_pca64_res0.55_downsampling5000.png

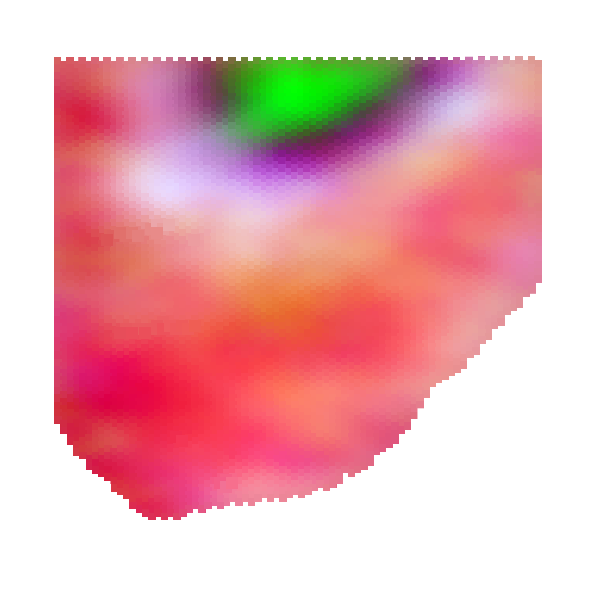

### S5_spaGCN_logcpm_pca128_res0.6_downsampling3000.png

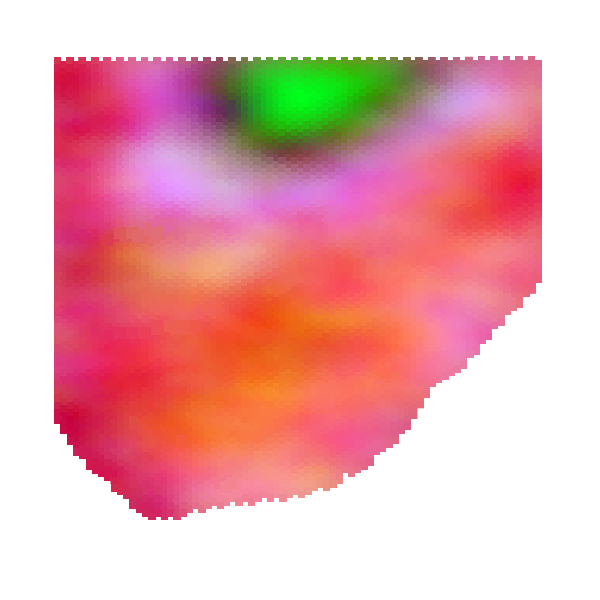

### S5_spaGCN_logcpm_pca128_res0.6_downsampling6500.png

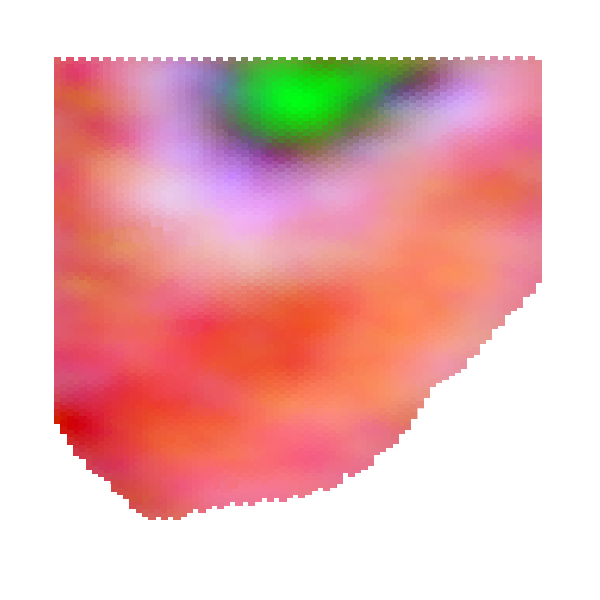

### S5_spaGCN_logcpm_pca128_res0.35_downsampling4500.png

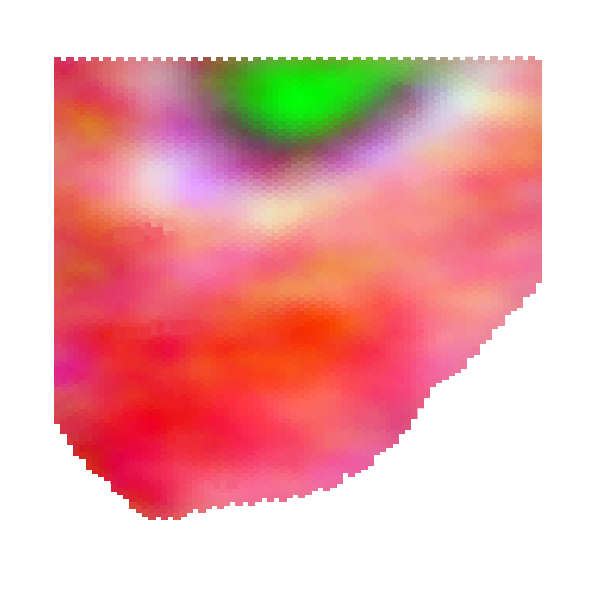

### S5_spaGCN_logcpm_pca1024_res0.3_downsampling8000.png

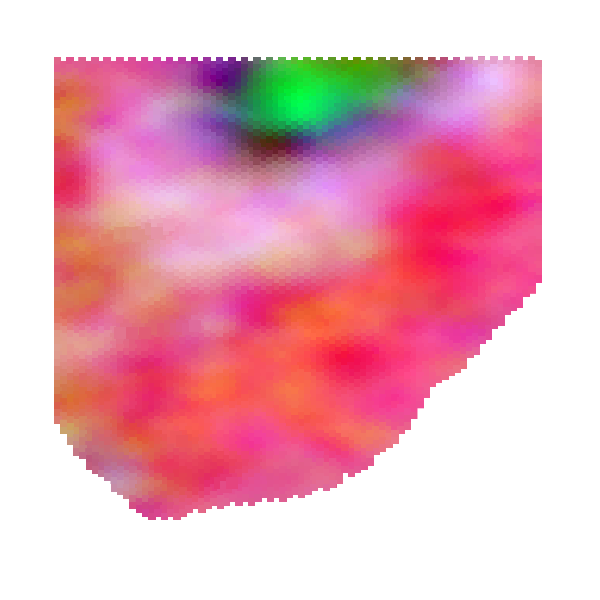

### S5_spaGCN_logcpm_pca1024_res0.4_downsampling6000.png

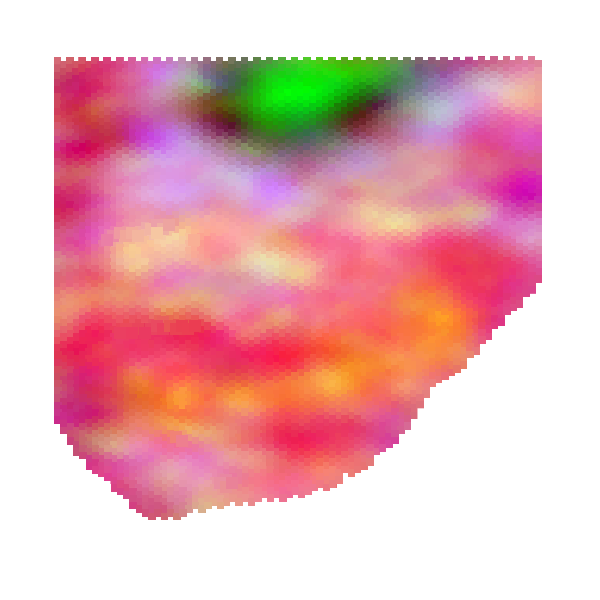

### S5_spaGCN_logcpm_pca1024_res0.25_downsampling7000.png

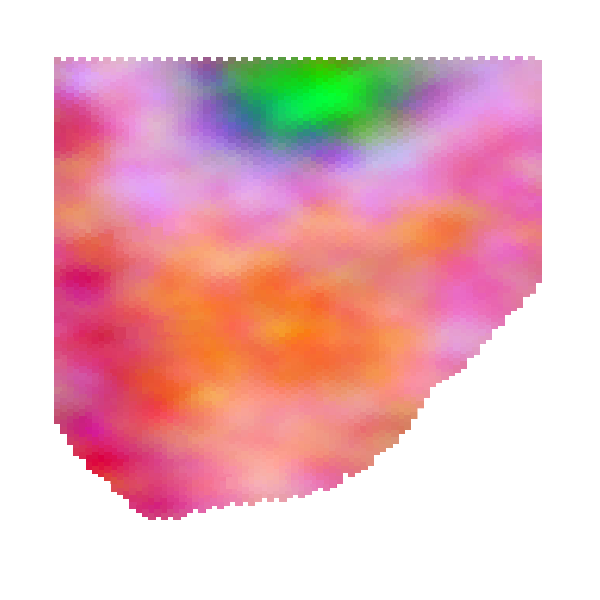

### S6_6_tools_DESeq2.jpeg

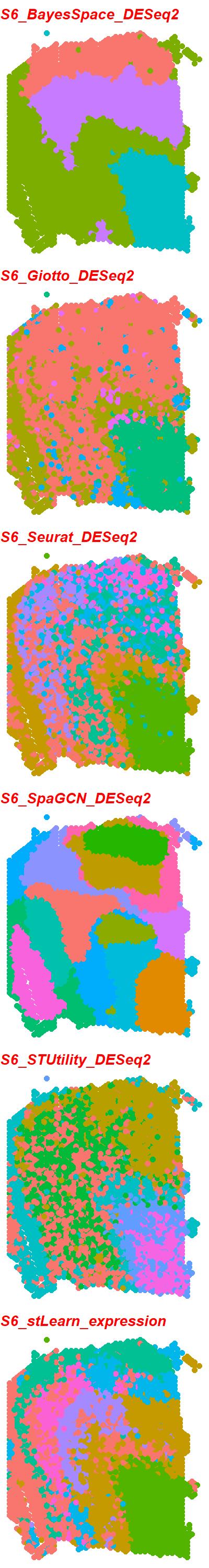

### S6_6_tools_expression.jpeg

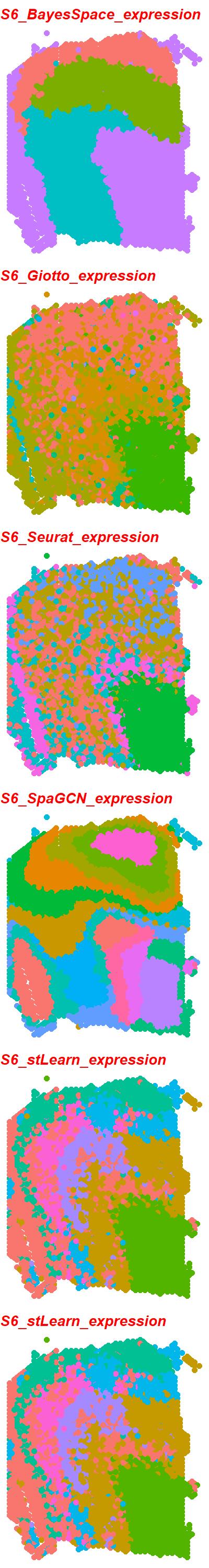

### S6_scGNN_logcpm_PEalpha0.1_zdim3_downsampling3000.png

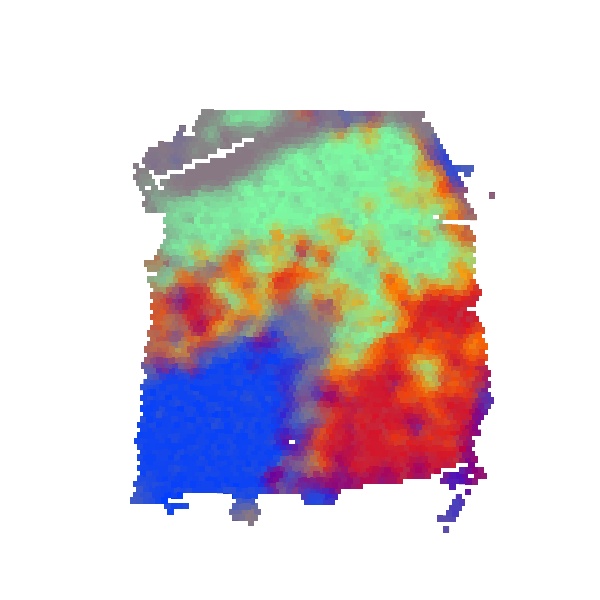

### S6_scGNN_logcpm_PEalpha0.1_zdim3_downsampling5000.png

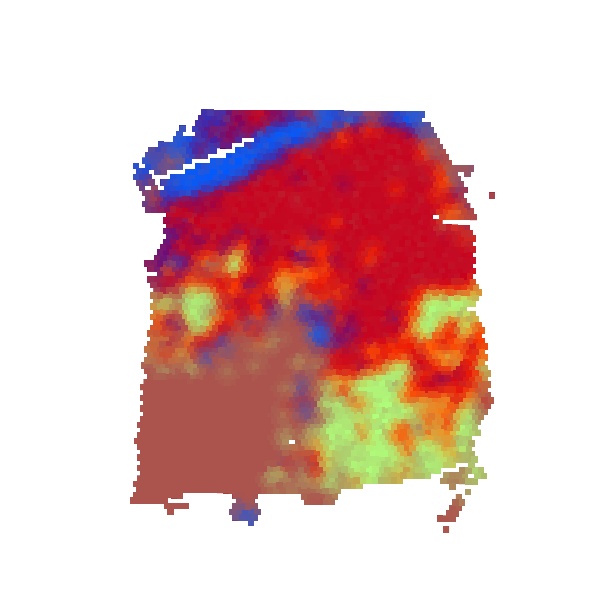

### S6_scGNN_logcpm_PEalpha0.1_zdim64_downsampling500.png

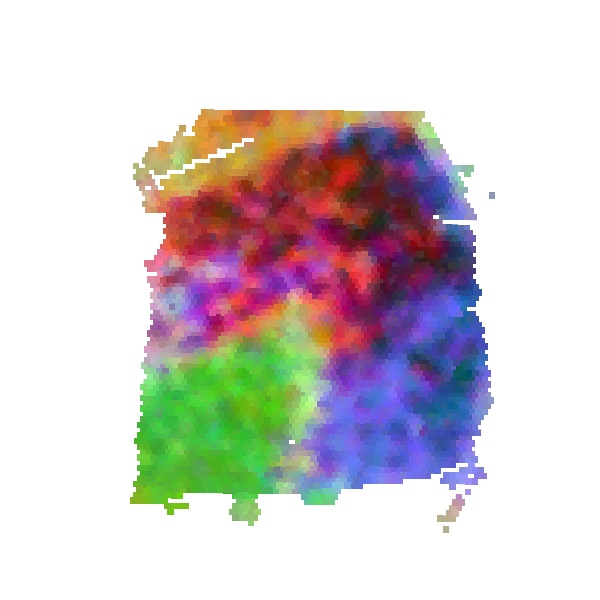

### S6_scGNN_logcpm_PEalpha0.2_zdim3_downsampling2500.png

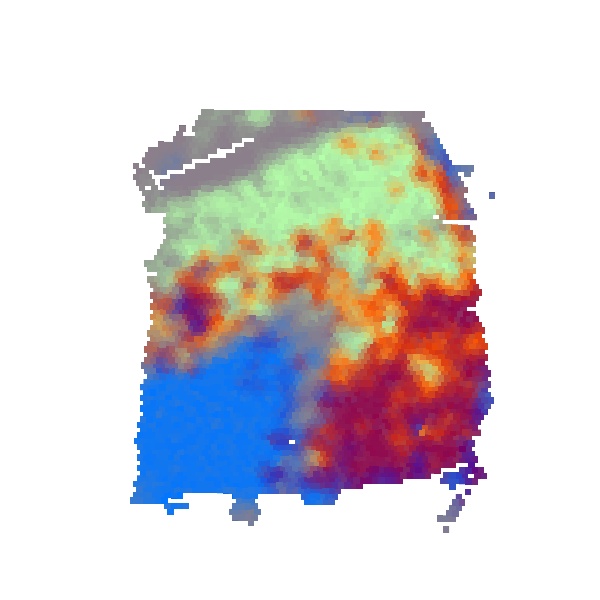

### S6_scGNN_logcpm_PEalpha0.2_zdim3_downsampling3500.png

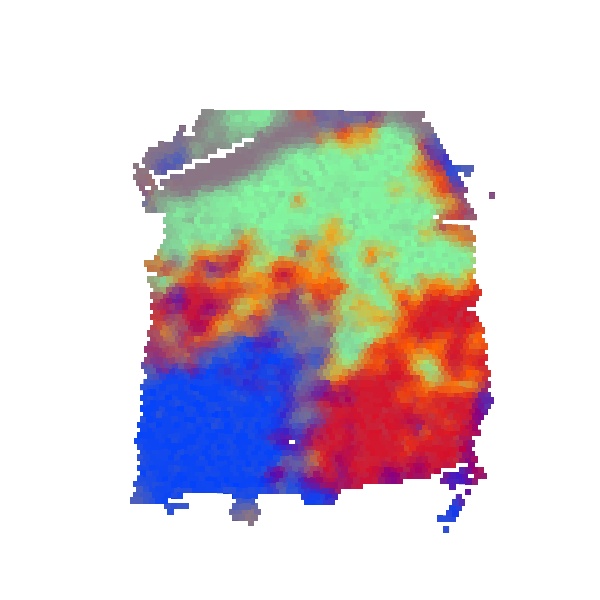

### S6_scGNN_logcpm_PEalpha0.3_zdim3_downsampling2000.png

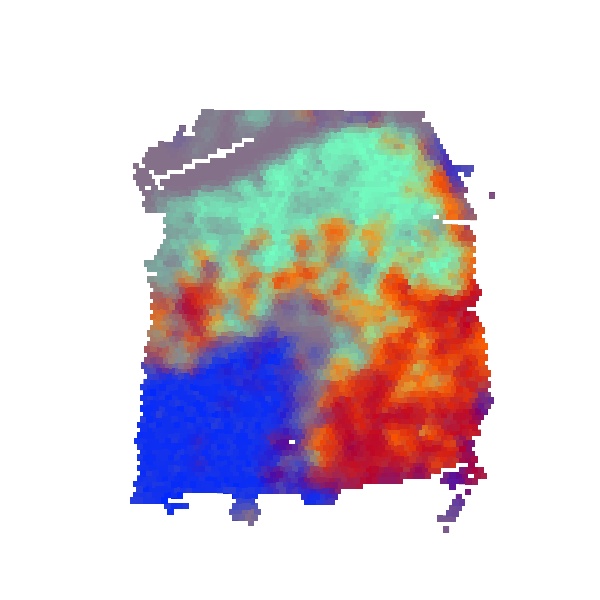

### S6_scGNN_logcpm_PEalpha0.5_zdim32_downsampling1000.png

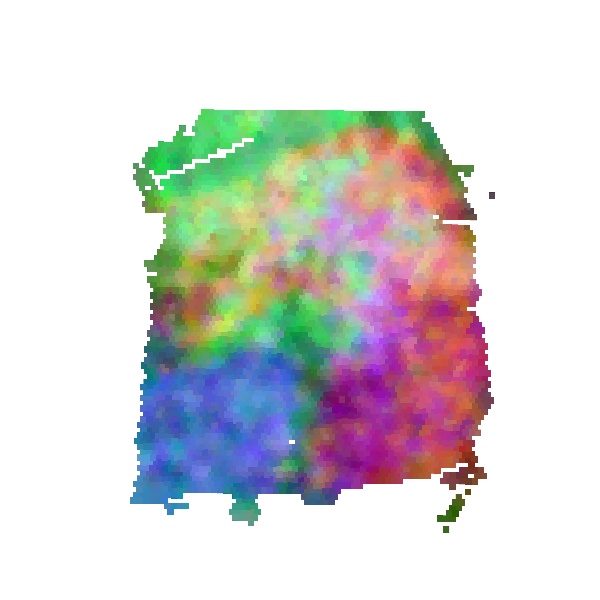

### S6_scGNN_logcpm_PEalpha1.0_zdim3_downsampling4000.png

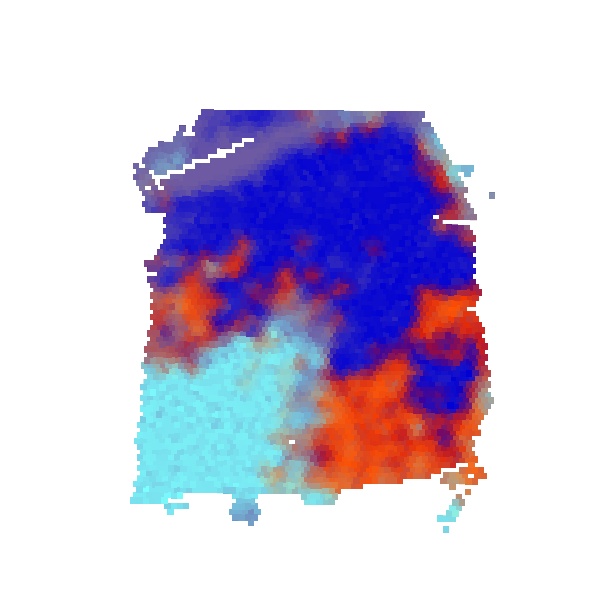

### S6_scGNN_logcpm_PEalpha1.0_zdim3_downsampling5500.png

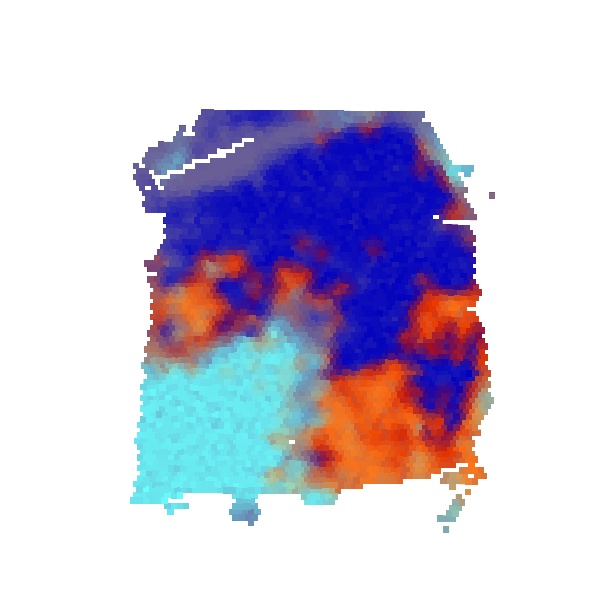

### S6_scGNN_logcpm_PEalpha1.0_zdim128_downsampling4500.png

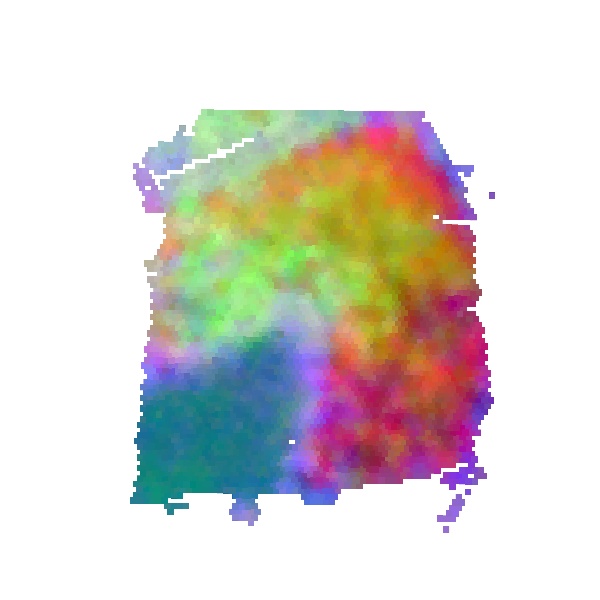

### S6_scGNN_logcpm_PEalpha1.2_zdim16_full.png

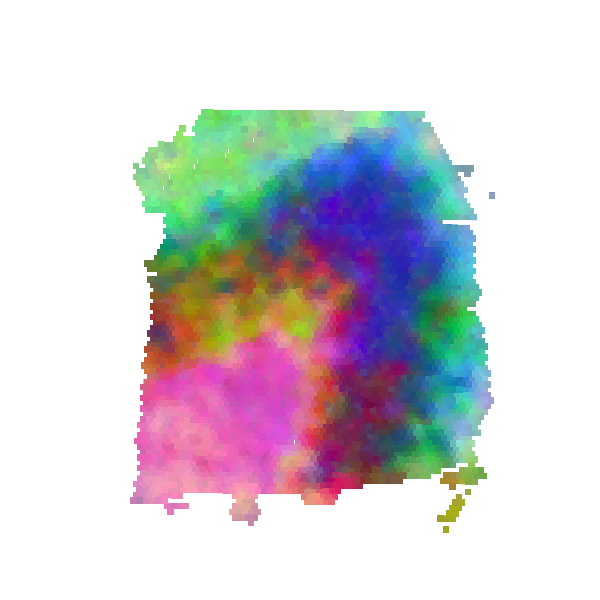

### S6_scGNN_logcpm_PEalpha1.2_zdim128_downsampling1500.png

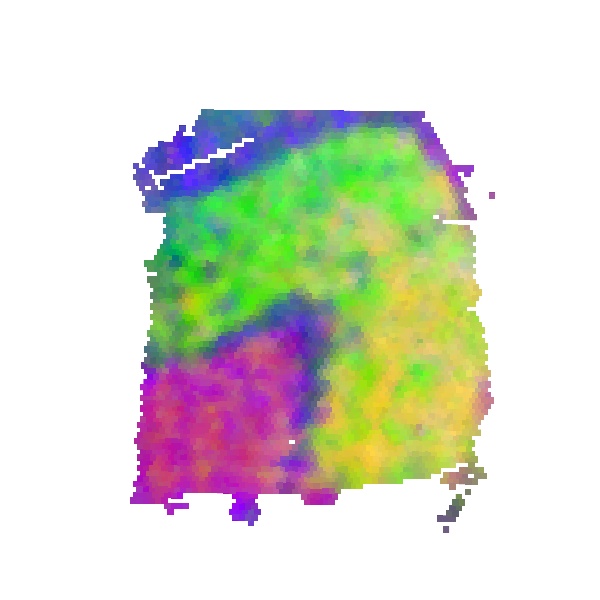

### S6_spaGCN_logcpm_pca32_res0.3_downsampling3500.png

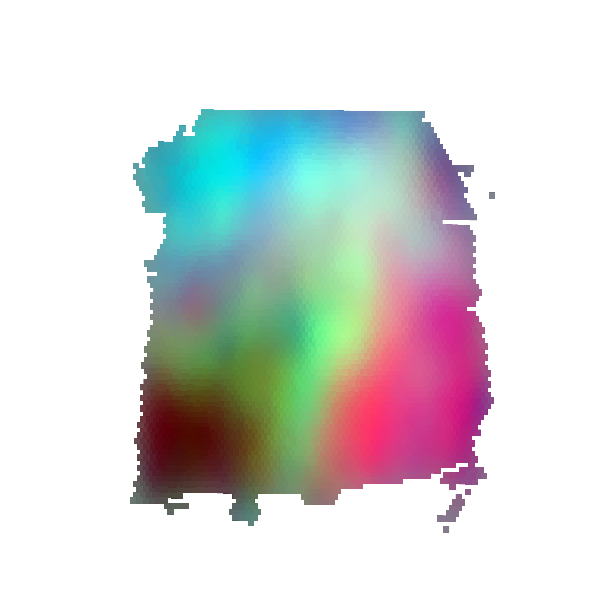

### S6_spaGCN_logcpm_pca32_res0.4_downsampling4000.png

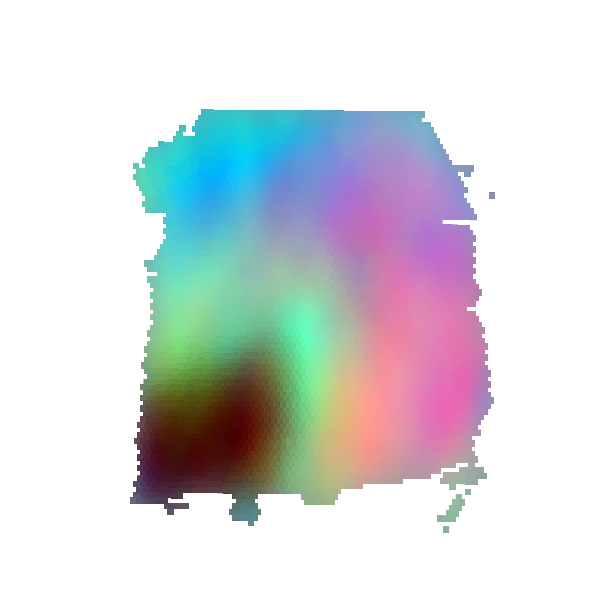

### S6_spaGCN_logcpm_pca32_res0.5_downsampling1500.png

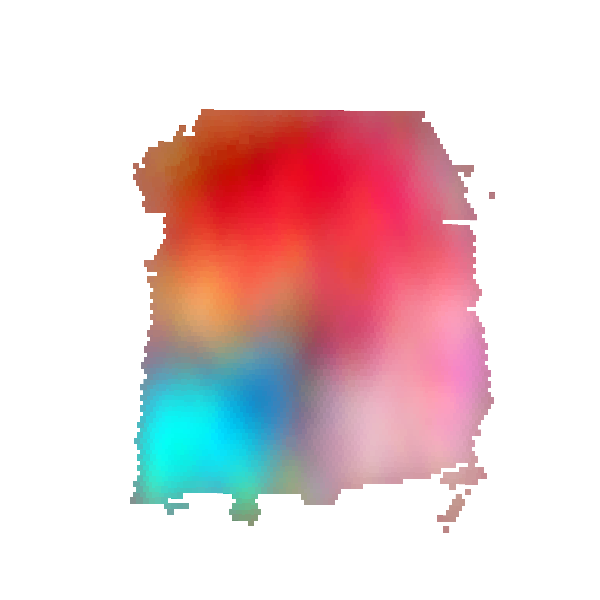

### S6_spaGCN_logcpm_pca32_res0.5_downsampling5500.png

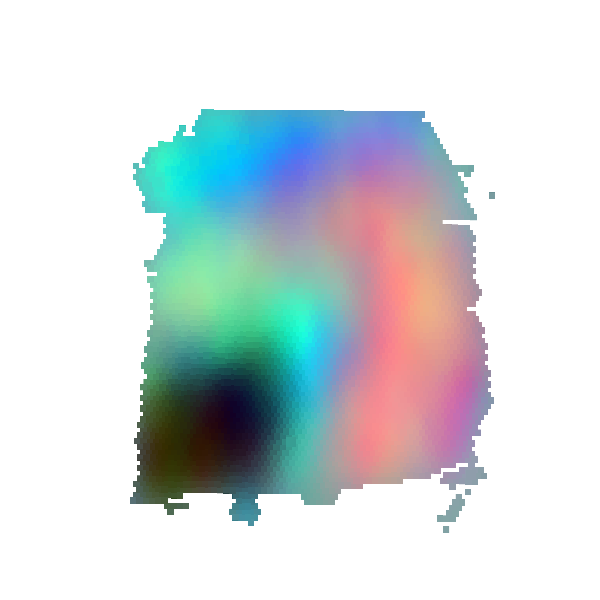

### S6_spaGCN_logcpm_pca32_res0.25_downsampling4500.png

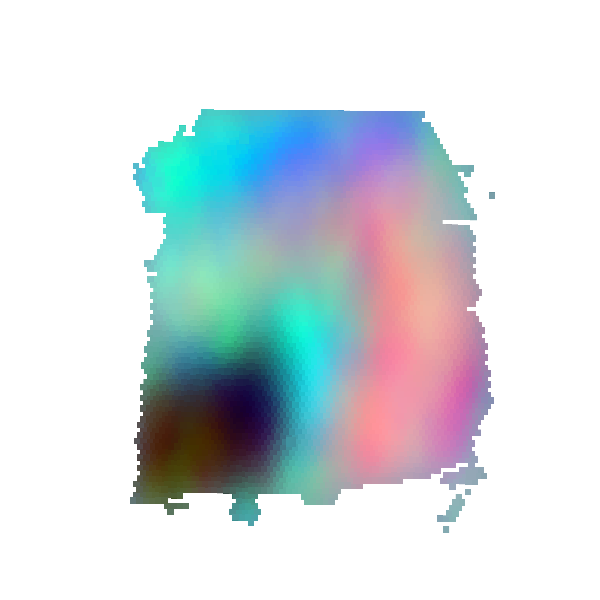

### S6_spaGCN_logcpm_pca50_res0.3_downsampling500.png

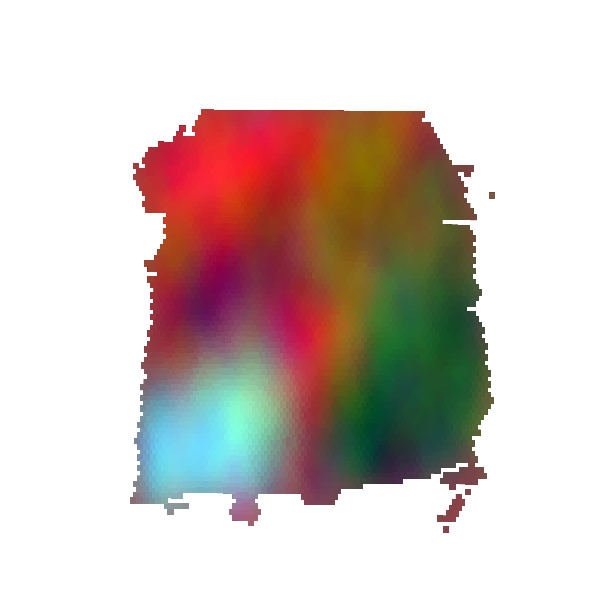

### S6_spaGCN_logcpm_pca50_res0.65_downsampling1000.png

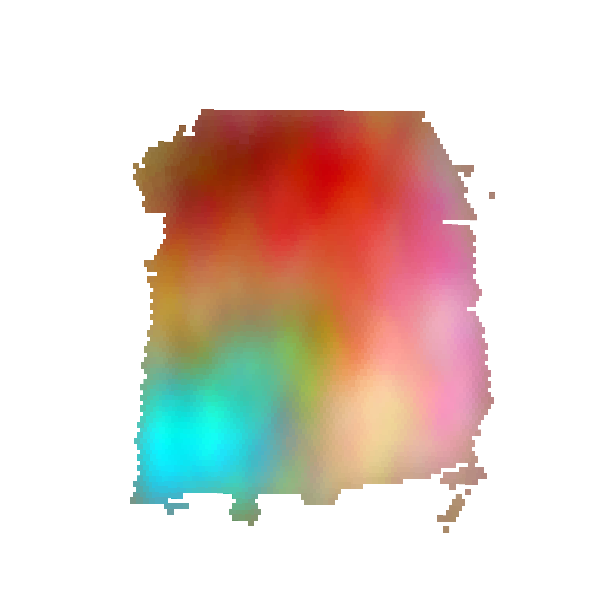

### S6_spaGCN_logcpm_pca64_res0.3_downsampling3000.png

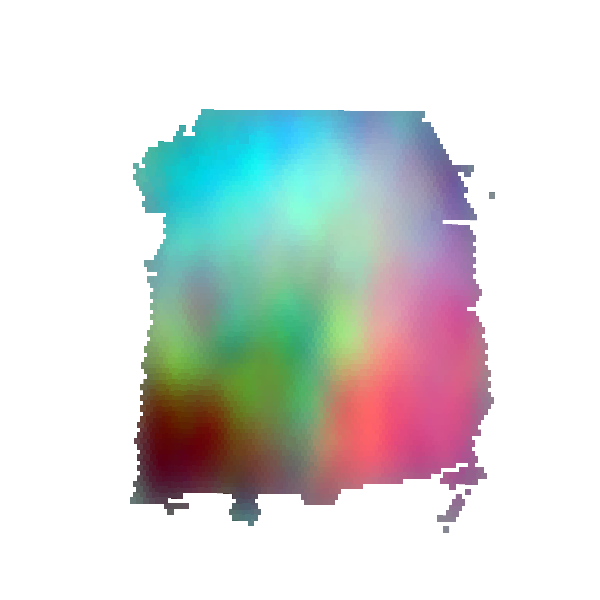

### S6_spaGCN_logcpm_pca128_res0.5_full.png

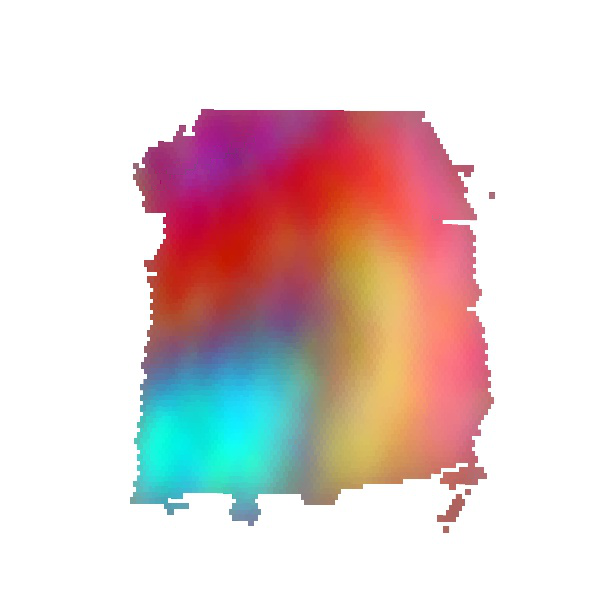

### S6_spaGCN_logcpm_pca128_res0.6_downsampling5000.png

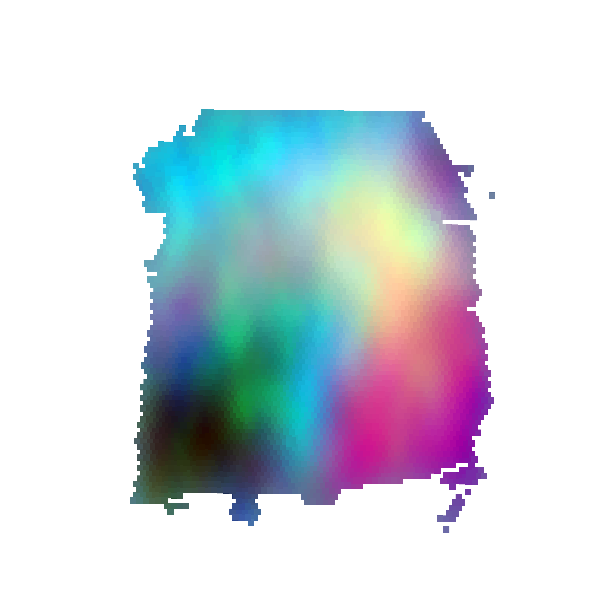

### S6_spaGCN_logcpm_pca256_res0.55_downsampling2500.png

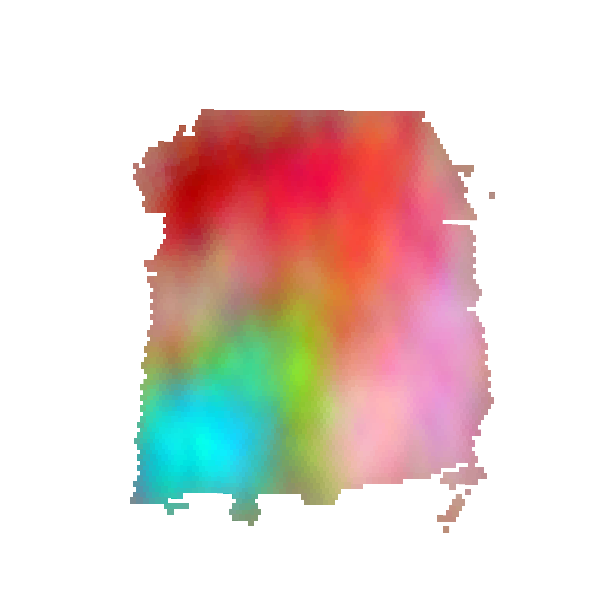

### S6_spaGCN_logcpm_pca1024_res0.35_downsampling2000.png

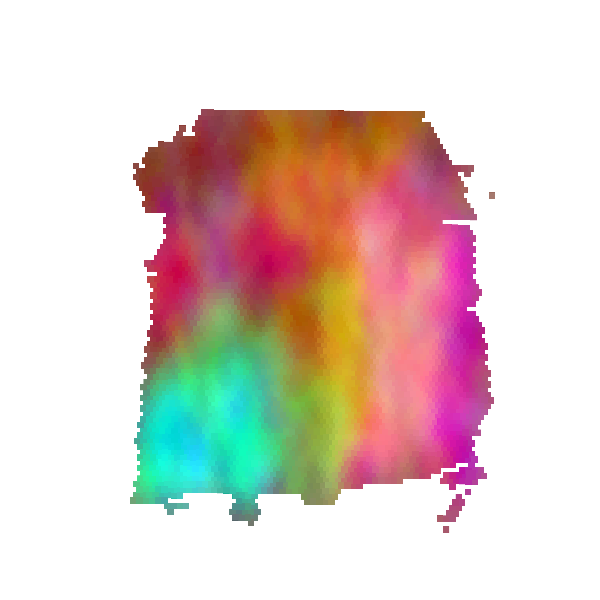

### S7_6_tools_DESeq2.jpeg

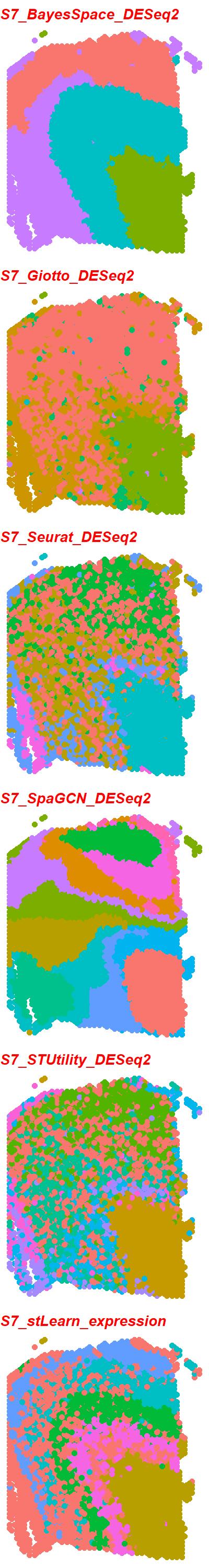

### S7_6_tools_expression.jpeg

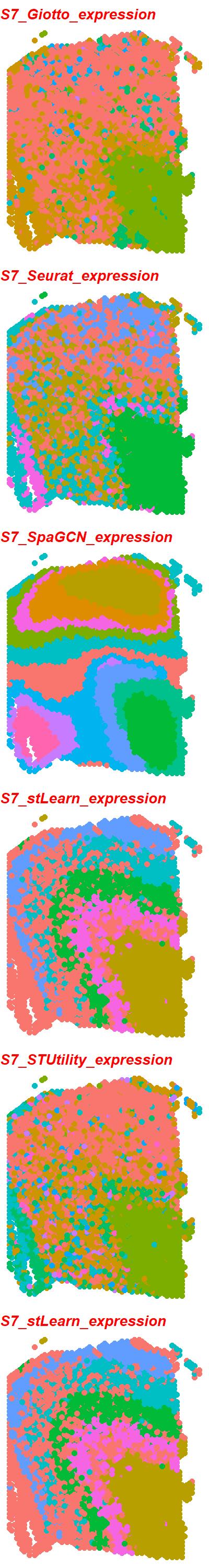

### S8_6_tools_DESeq2.jpeg

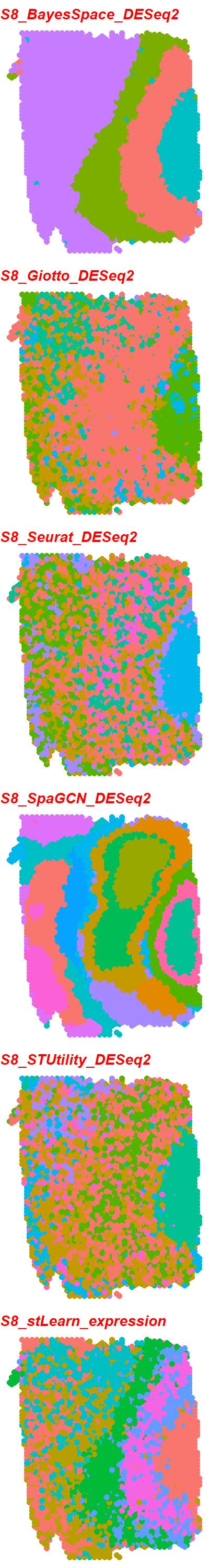

### S8_6_tools_expression.jpeg

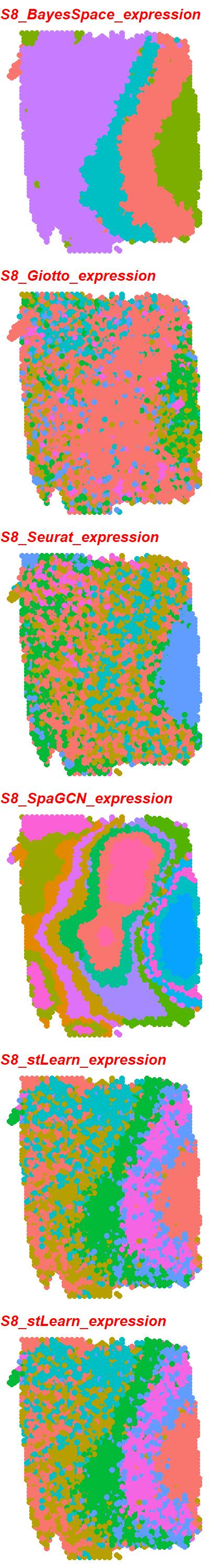

### S9_6_tools_DESeq2.jpeg

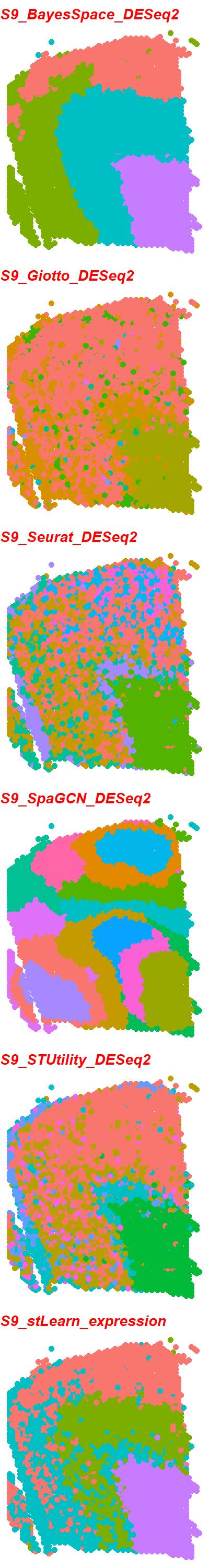

### S9_6_tools_expression.jpeg

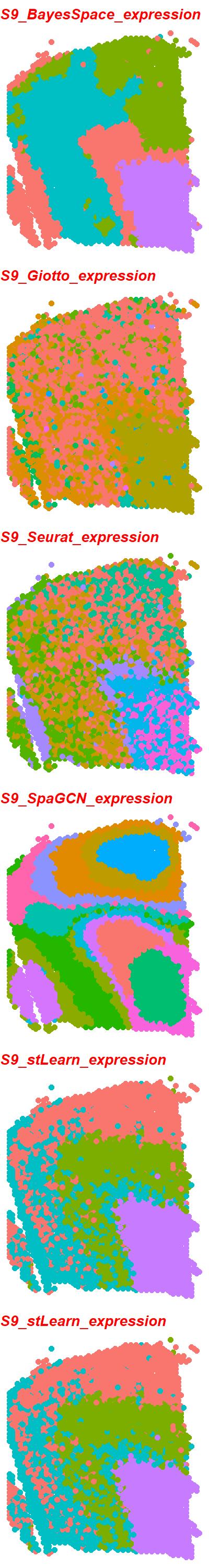

### S10_6_tools_DESeq2.jpeg

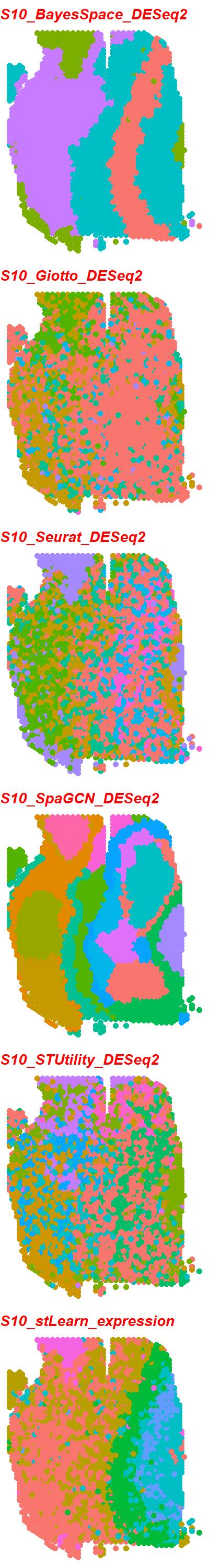

### S10_6_tools_expression.jpeg

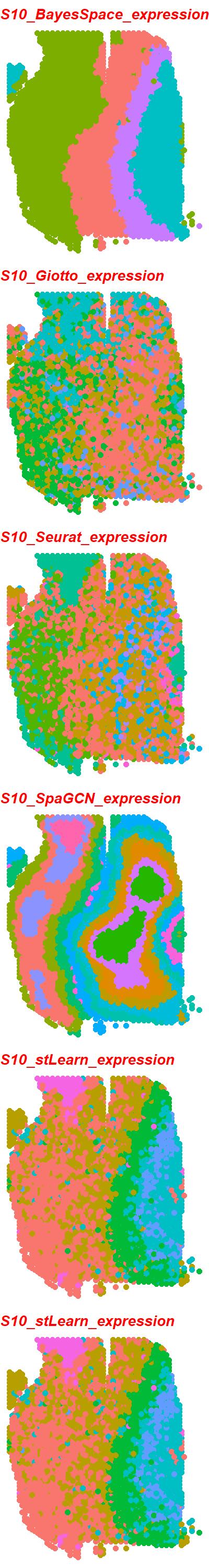

### S10_6_tools_LogCPM.jpeg

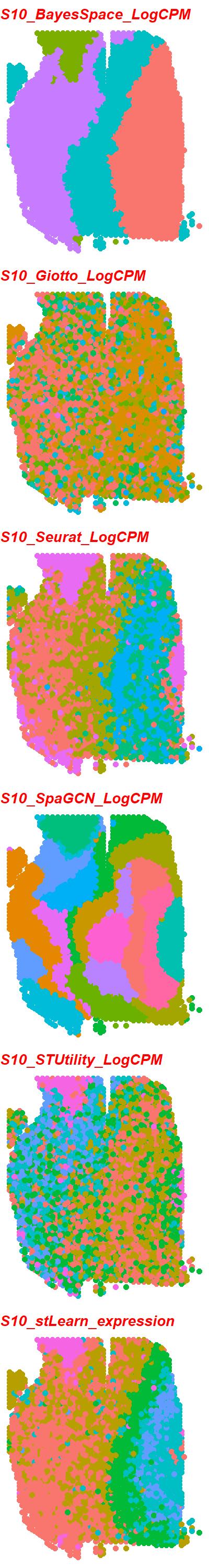

### S11_6_tools_DESeq2.jpeg

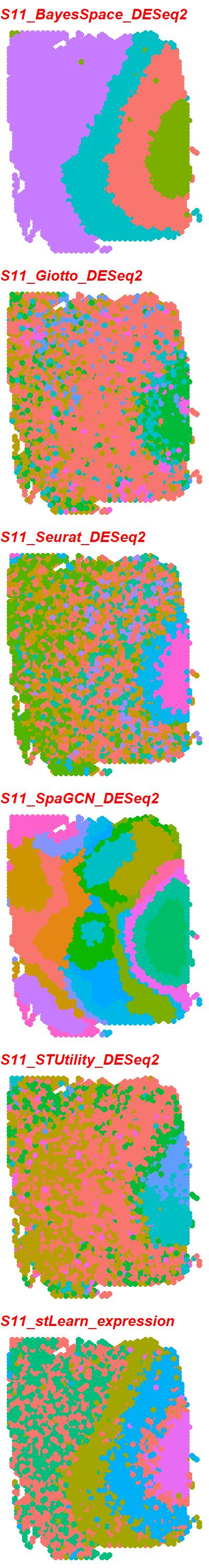

### S11_6_tools_expression.jpeg

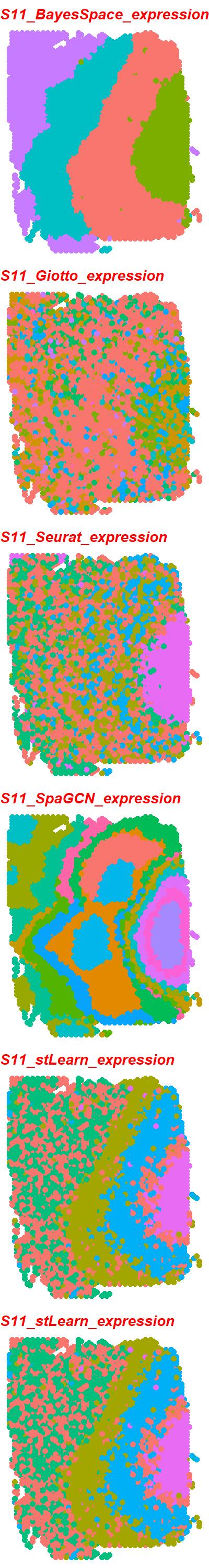

### S11_6_tools_LogCPM.jpeg

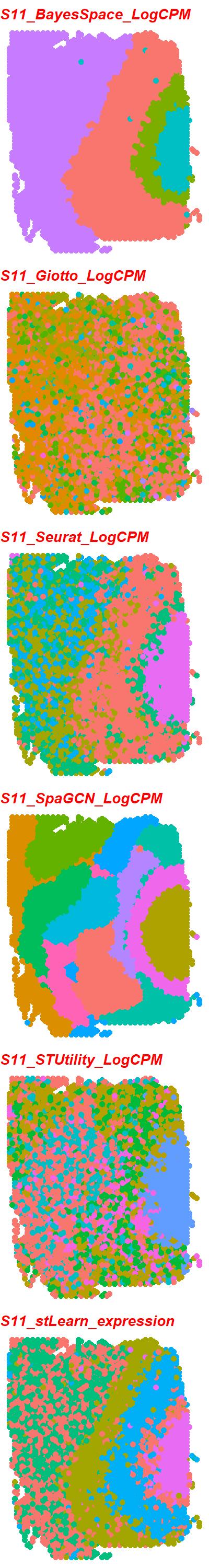

### S12_6_tools_DESeq2.jpeg

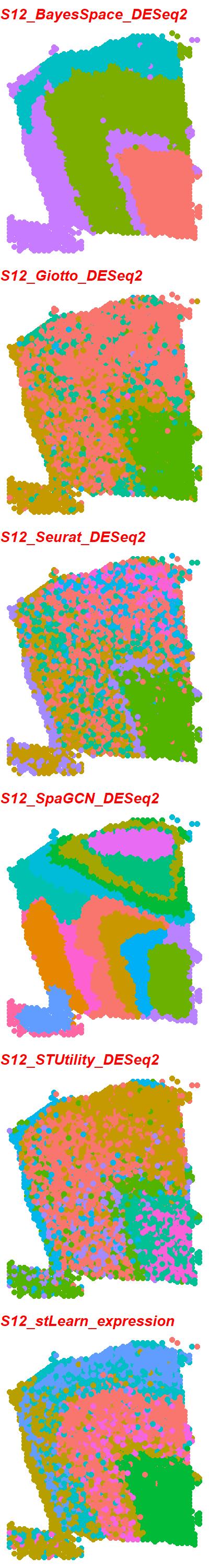

### S12_6_tools_expression.jpeg

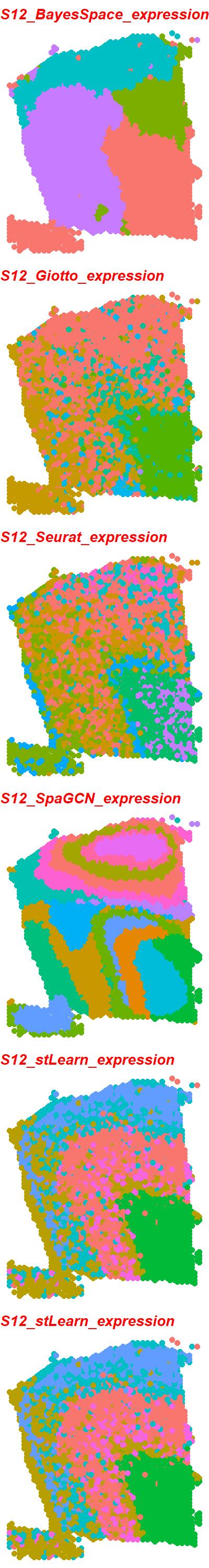

### S12_6_tools_LogCPM.jpeg

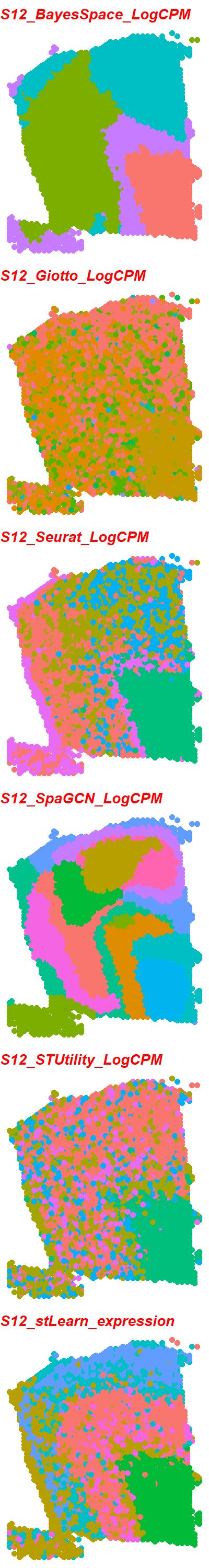

### S13_6_tools_DESeq2.jpeg

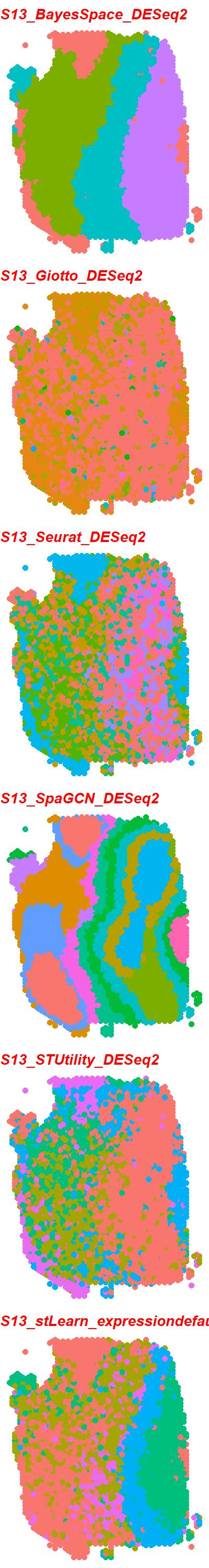

### S13_6_tools_expression.jpeg

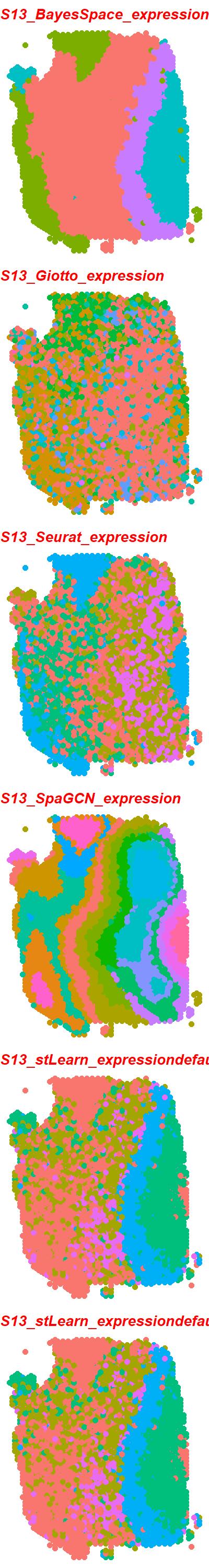

### S13_6_tools_LogCPM.jpeg

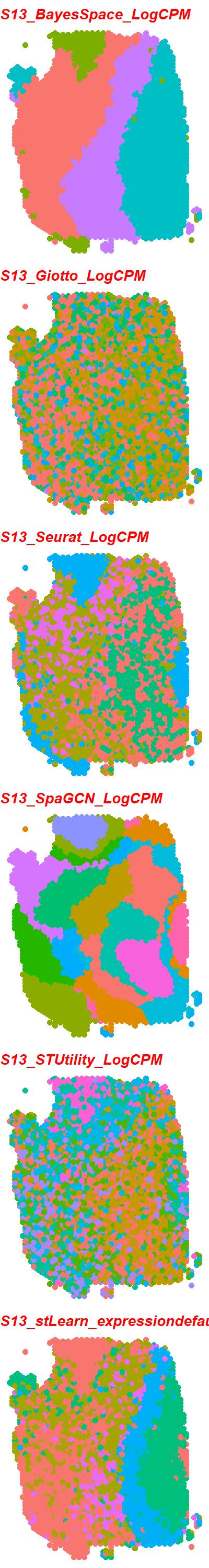

### S14_6_tools_DESeq2.jpeg

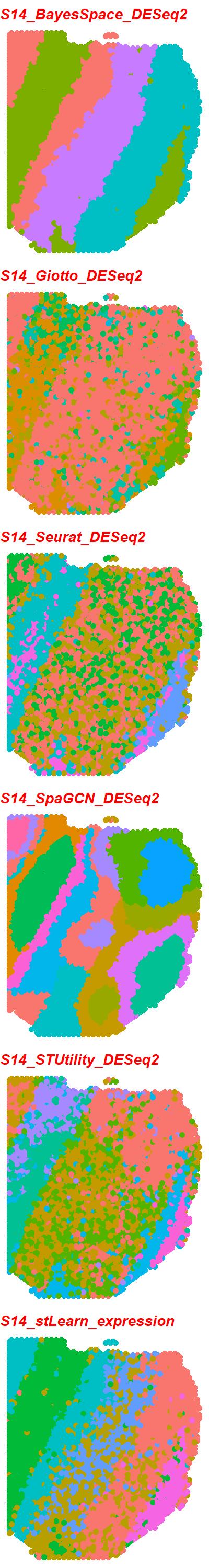

### S14_6_tools_expression.jpeg

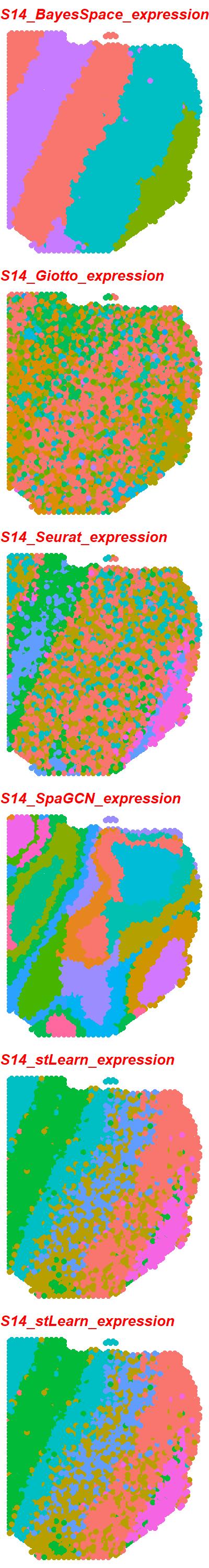

### S14_6_tools_LogCPM.jpeg

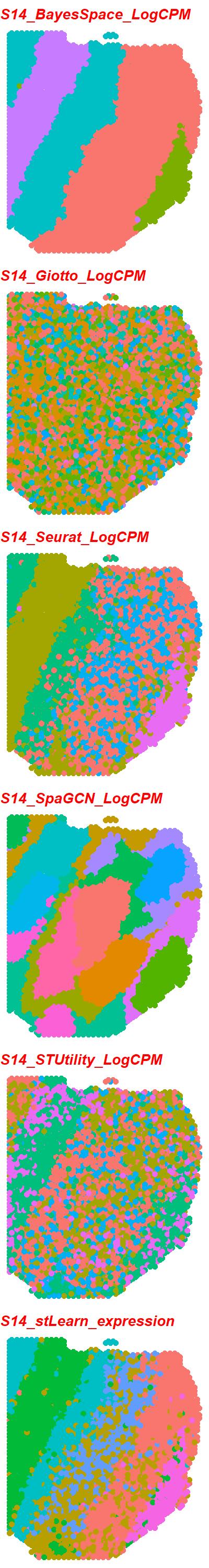

### S15_6_tools_DESeq2.jpeg

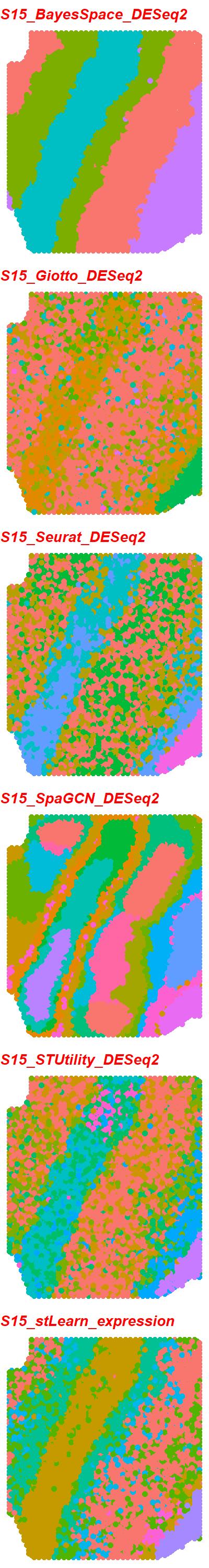

### S15_6_tools_expression.jpeg

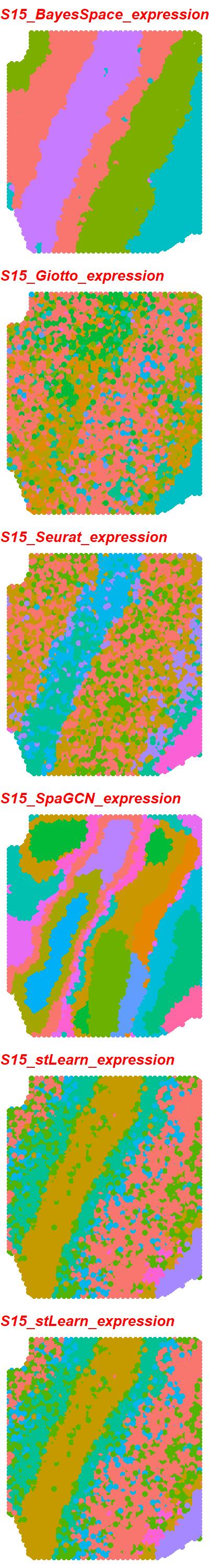

### S15_6_tools_LogCPM.jpeg

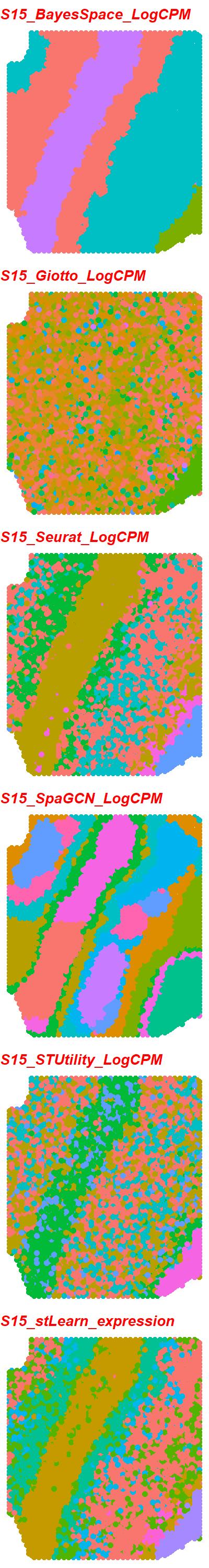

### S16_6_tools_DESeq2.jpeg

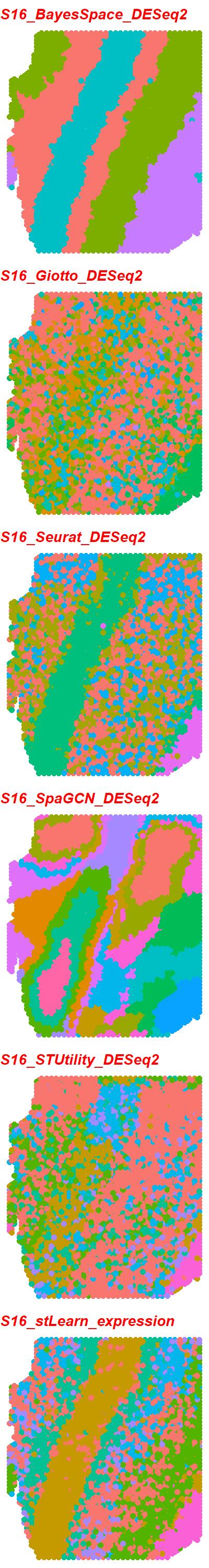

### S16_6_tools_expression.jpeg

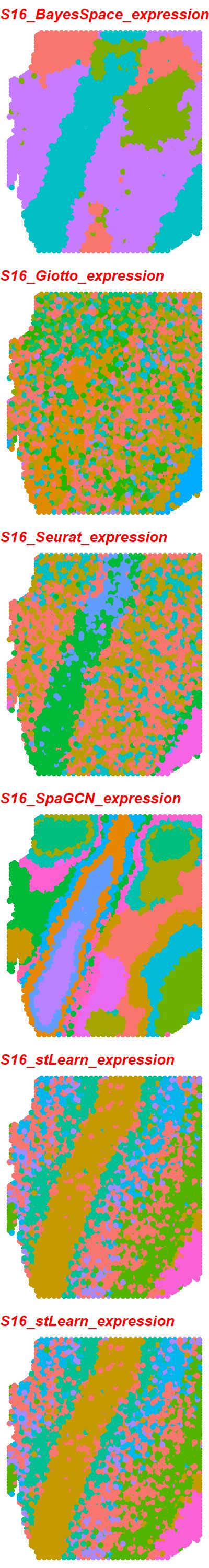

### S16_6_tools_LogCPM.jpeg

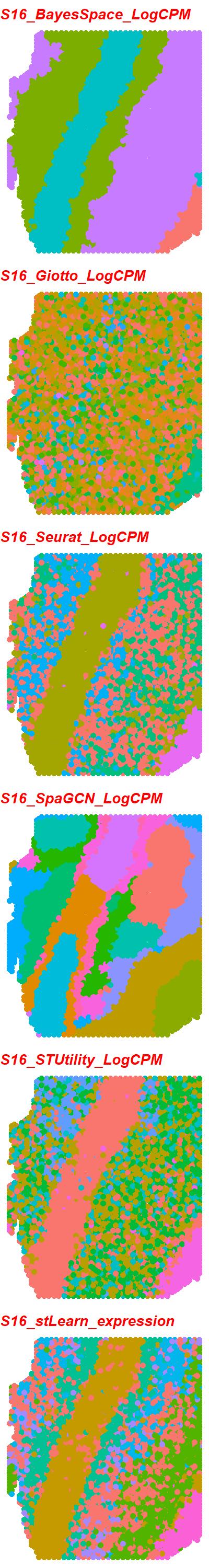

### S17_6_tools_DESeq2.jpeg

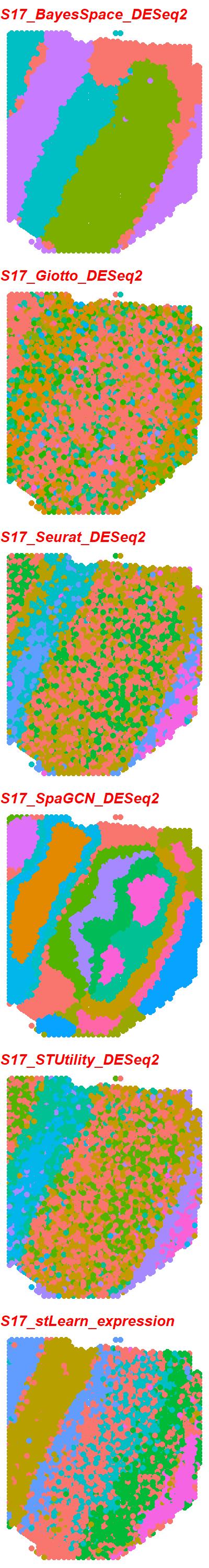

### S17_6_tools_expression.jpeg

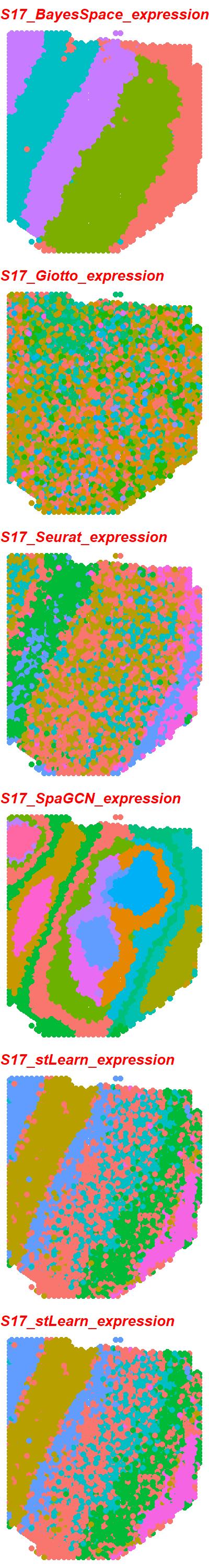

### S17_6_tools_LogCPM.jpeg

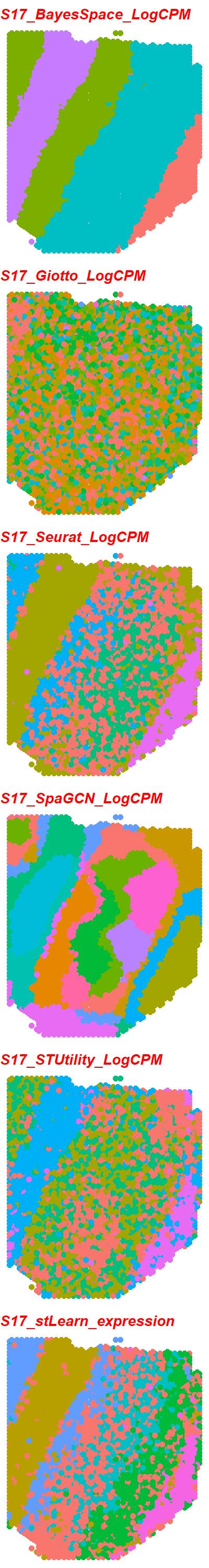
