## Supplementary figures and images for "Define and visualize pathological architectures of human tissues from spatially resolved transcriptomics using deep learning"

### S2_scGNN_expression_PEalpha0.5_zdim128.png

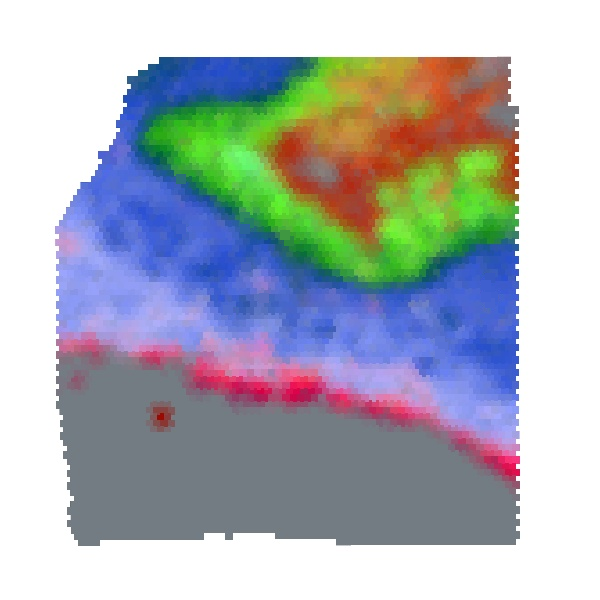

### S2_scGNN_RPKM_PEalpha1.0_zdim16.png

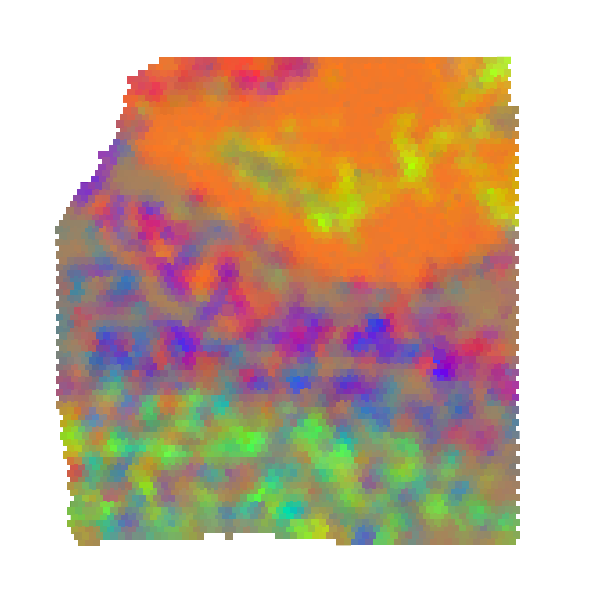

### S2_scGNN_scran_PEalpha0.5_zdim10.png

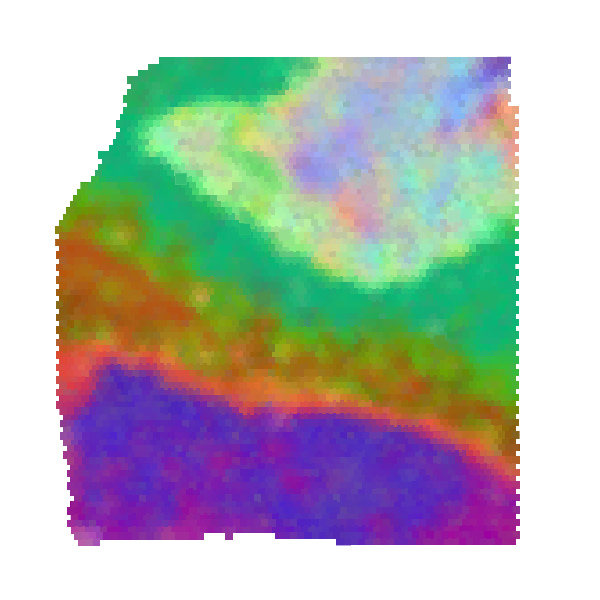

### S3_scGNN_expression_PEalpha0.5_zdim10.png

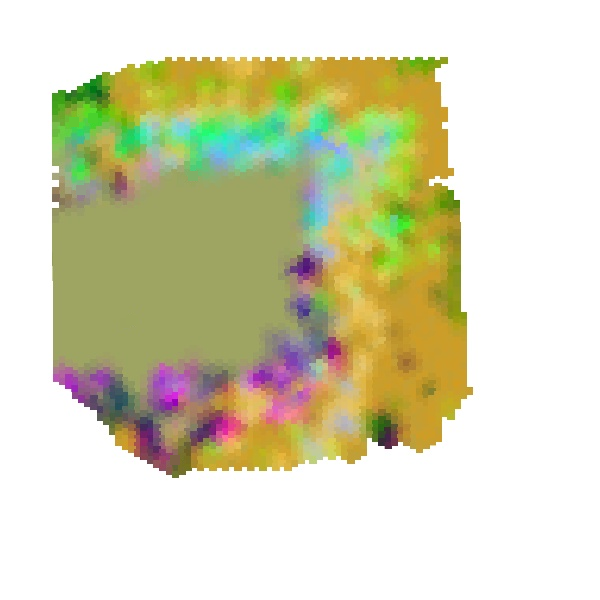

### S3_scGNN_RPKM_PEalpha0.5_zdim128.png

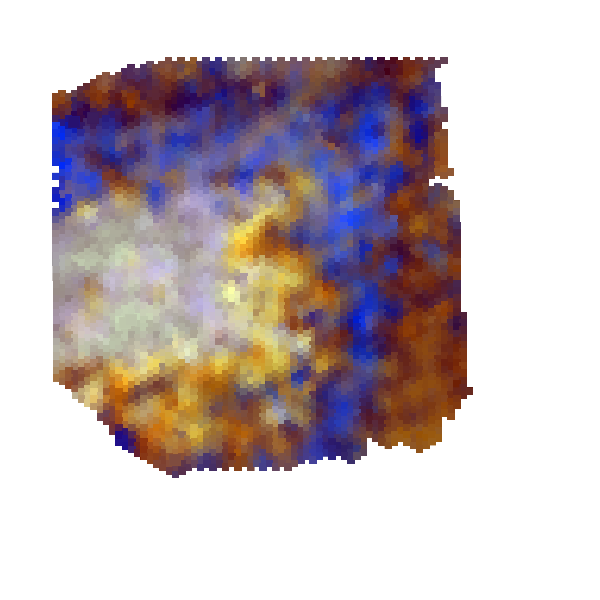

### S3_scGNN_scran_PEalpha2.0_zdim10.png

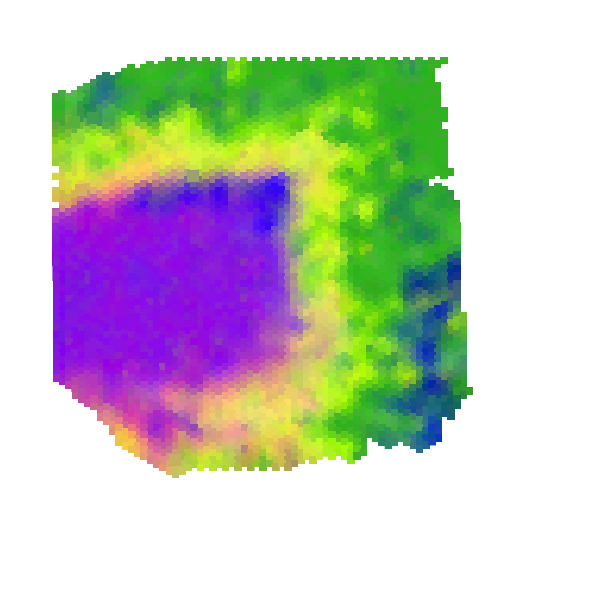

### S4_scGNN_expression_PEalpha1.5_zdim10.png

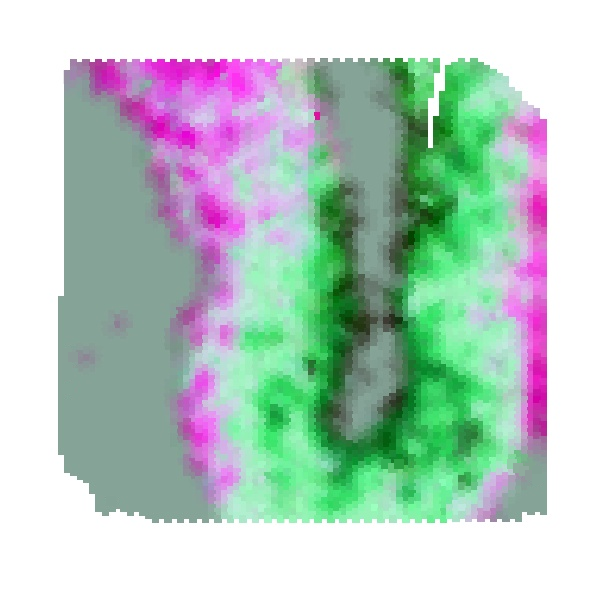

### S4_scGNN_RPKM_PEalpha0.5_zdim10.png

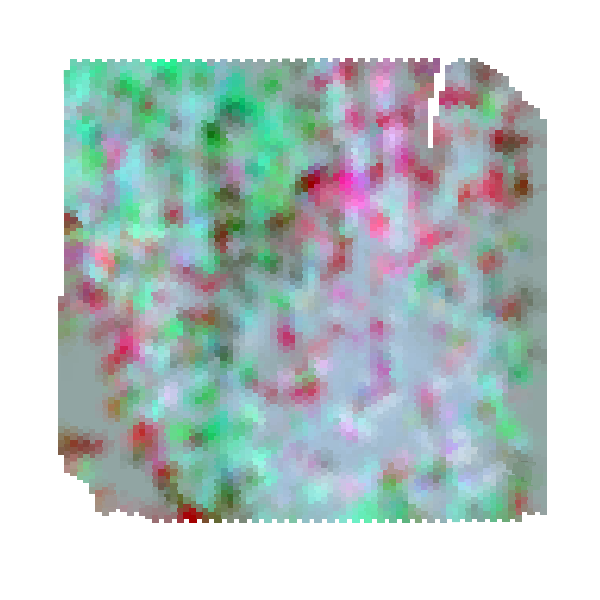

### S4_scGNN_scran_PEalpha1.5_zdim64.png

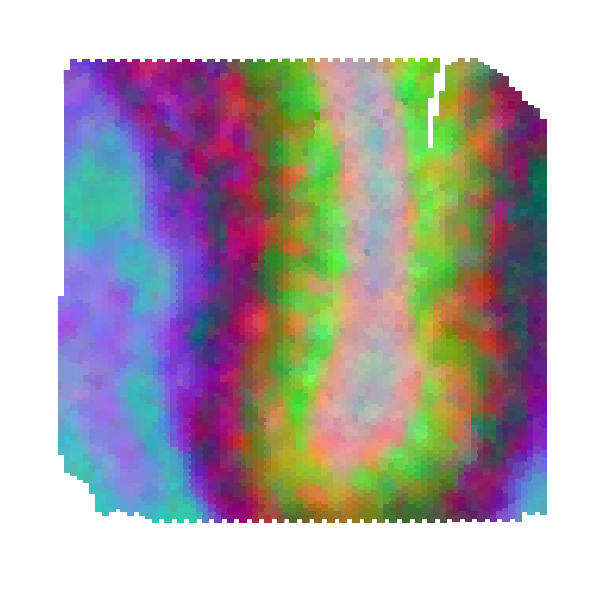

### S5_scGNN_expression_PEalpha0.1_zdim3.png

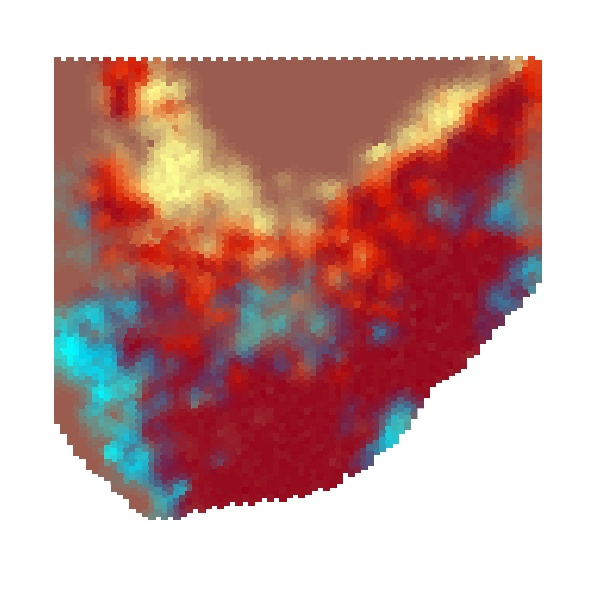

### S5_scGNN_RPKM_PEalpha0.2_zdim10.png

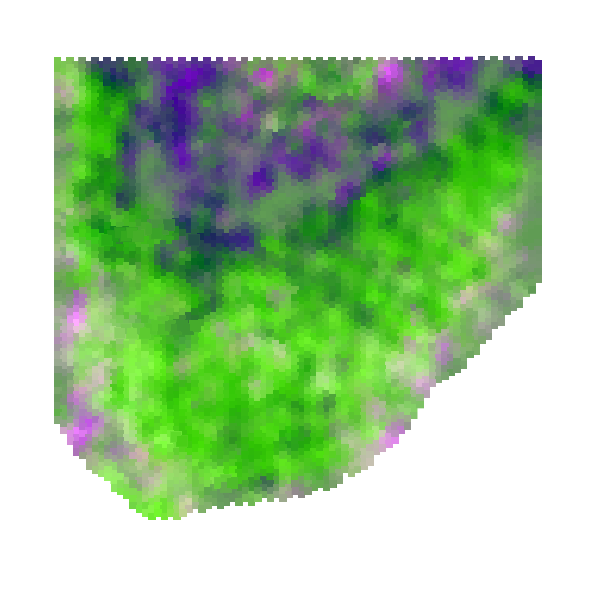

### S5_scGNN_scran_PEalpha0.3_zdim32.png

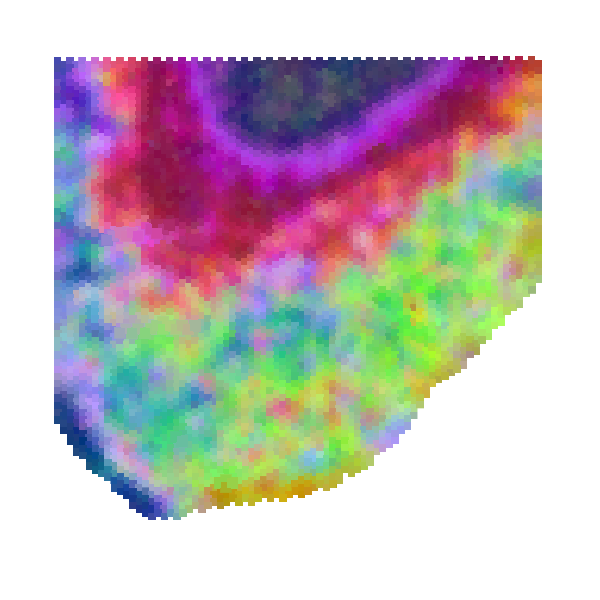

### S6_scGNN_expression_PEalpha2.0_zdim256.png

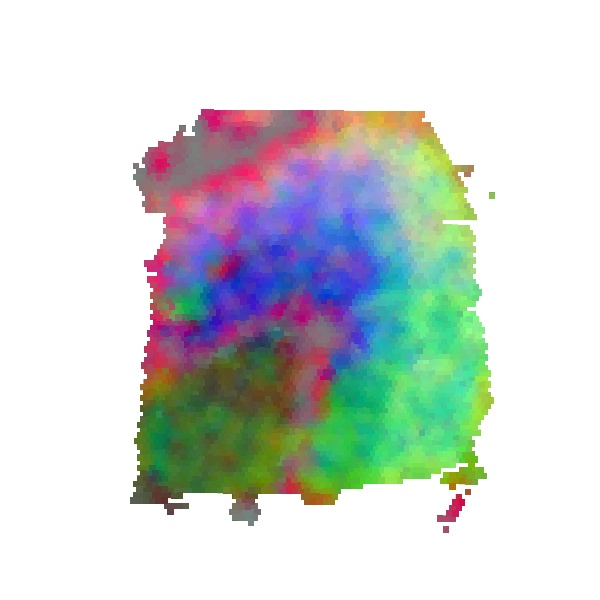

### S6_scGNN_RPKM_PEalpha1.0_zdim3.png

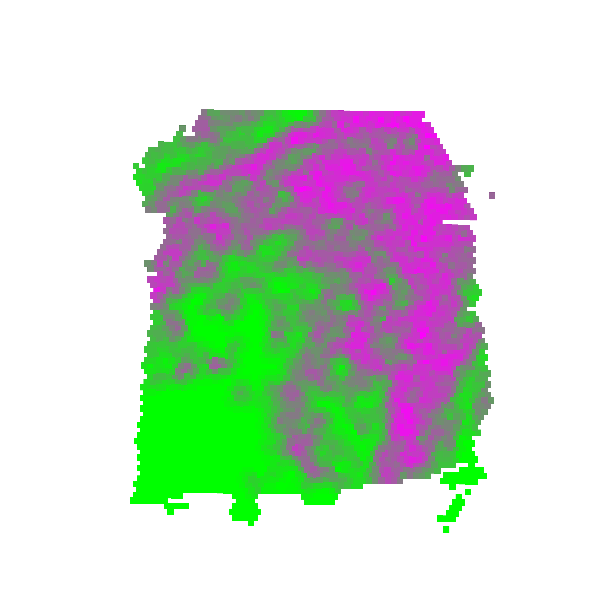

### S6_scGNN_scran_PEalpha0.1_zdim32.png

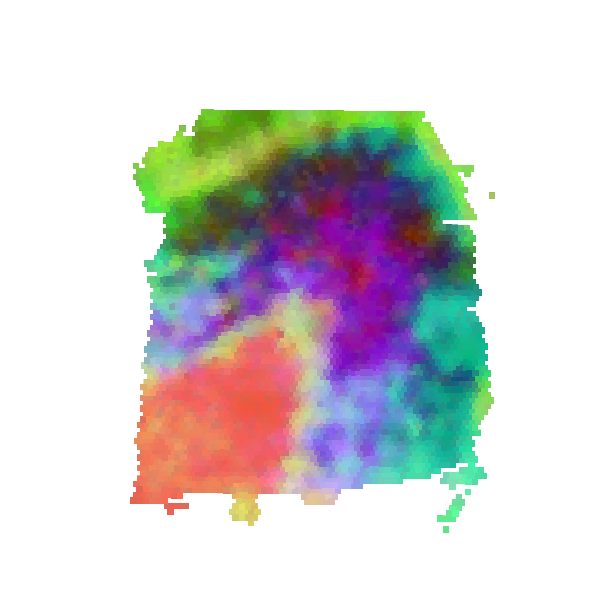

### S7_scGNN_expression_PEalpha0.2_zdim256.png

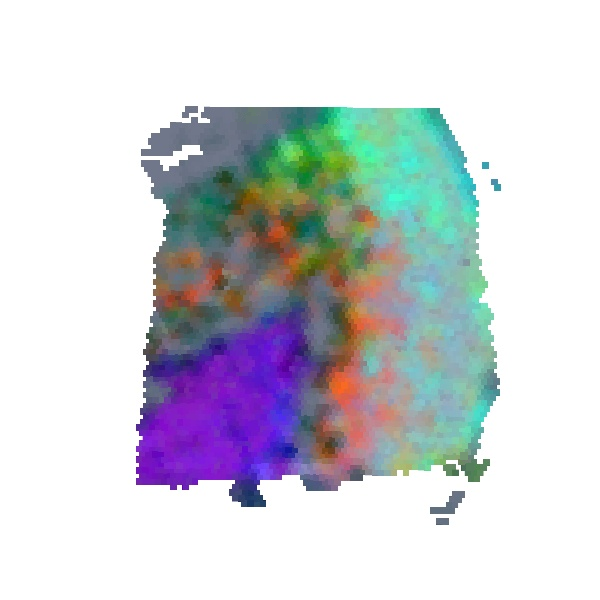

### S7_scGNN_RPKM_PEalpha0.3_zdim128.png

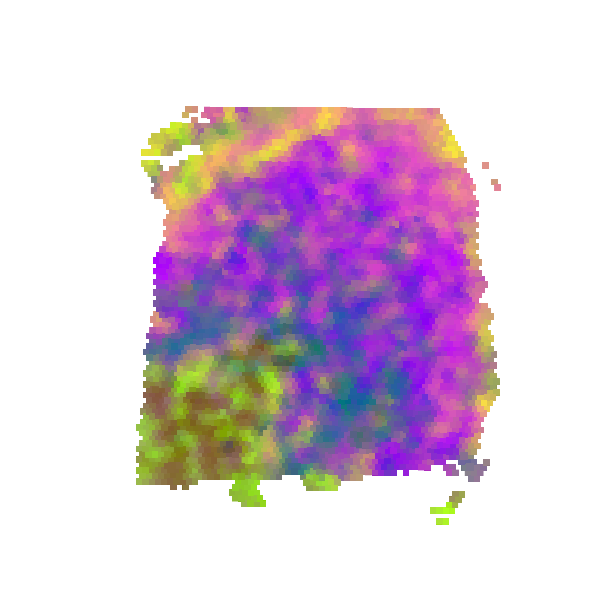

### S7_scGNN_scran_PEalpha1.0_zdim32.png

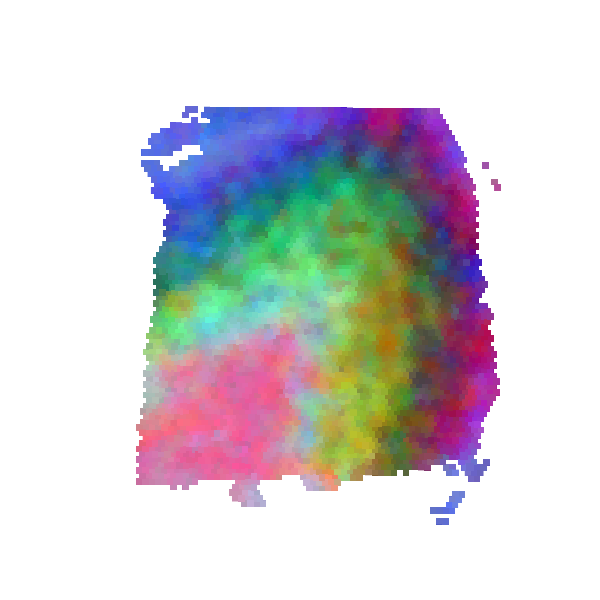

### S8_scGNN_DESeq2_PEalpha1.2_zdim3.png

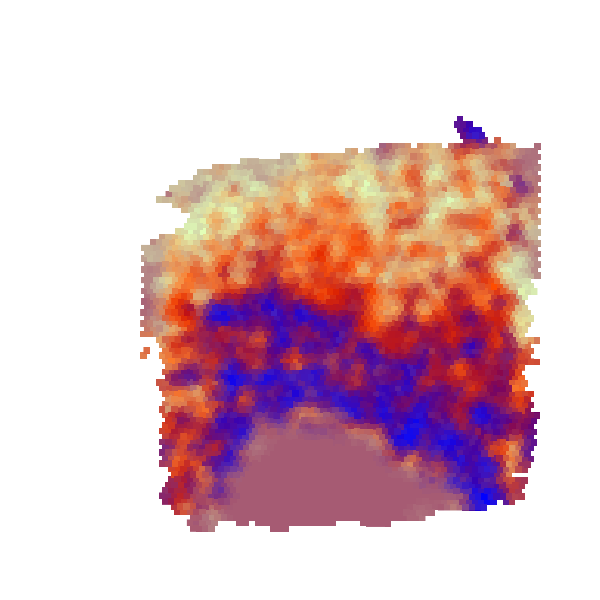

### S8_scGNN_expression_PEalpha2.0_zdim10.png

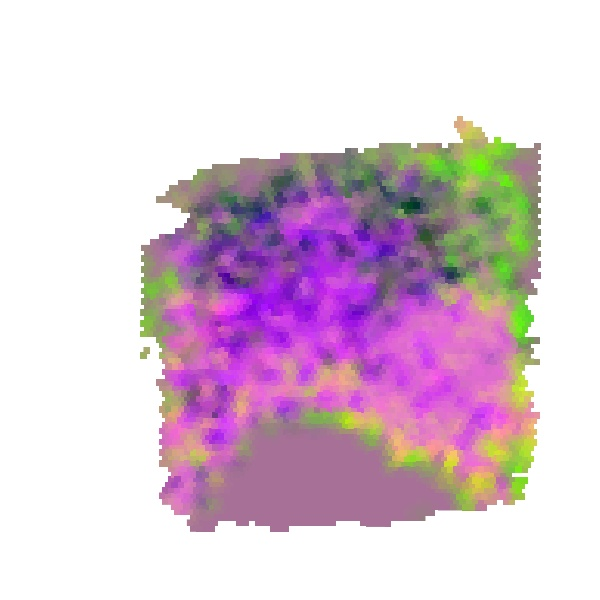

### S8_scGNN_RPKM_PEalpha0.5_zdim64.png

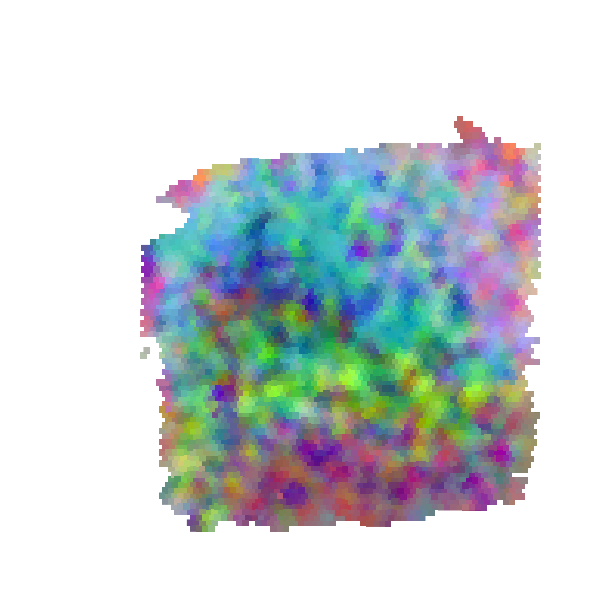

### S8_scGNN_scran_PEalpha0.1_zdim256.png

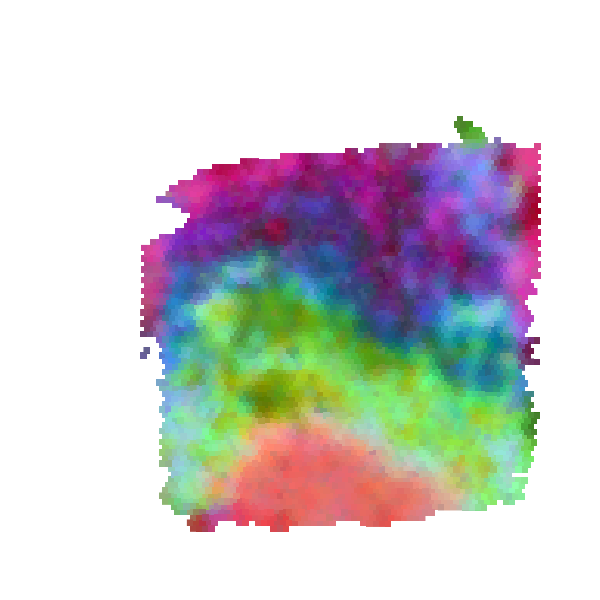

### S9_scGNN_DESeq2_PEalpha2.0_zdim16.png

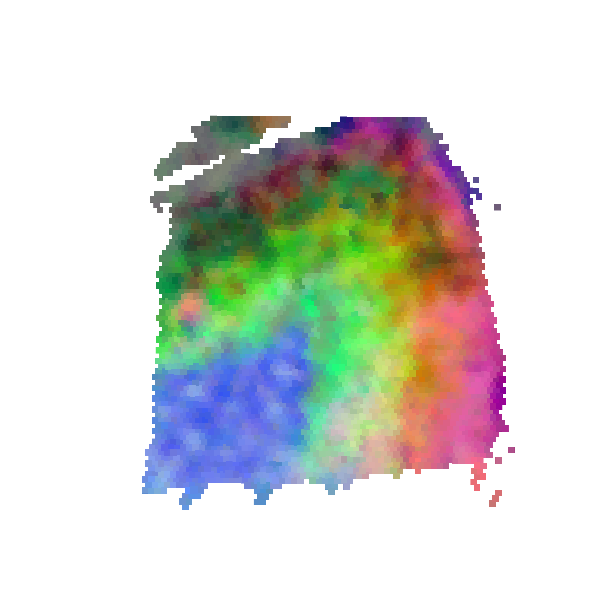

### S9_scGNN_expression_PEalpha1.0_zdim32.png

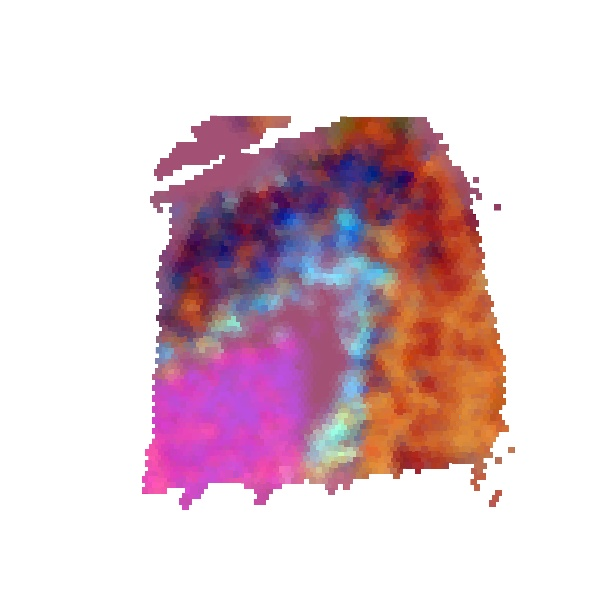

### S9_scGNN_RPKM_PEalpha1.5_zdim256.png

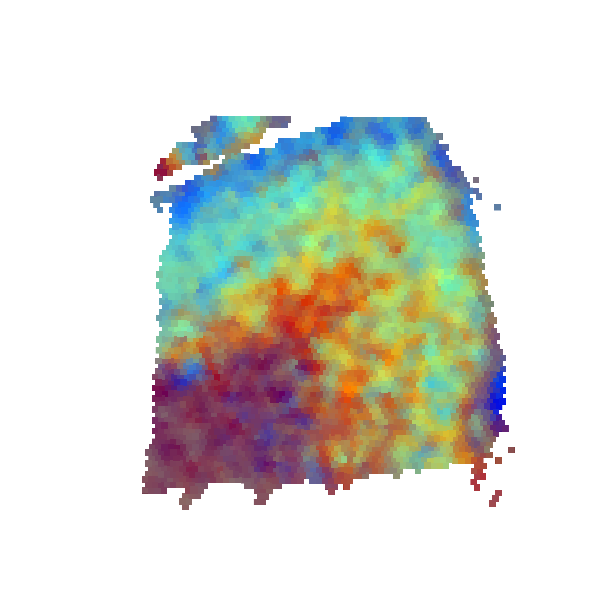

### S9_scGNN_scran_PEalpha0.2_zdim64.png

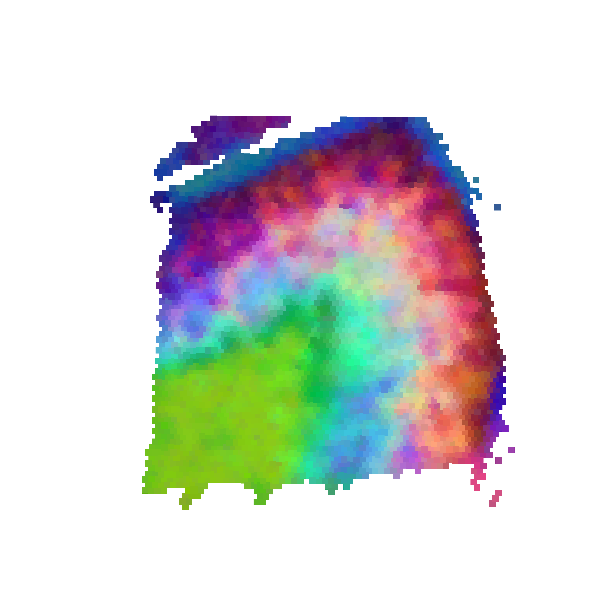

### S10_scGNN_expression_PEalpha2.0_zdim16.png

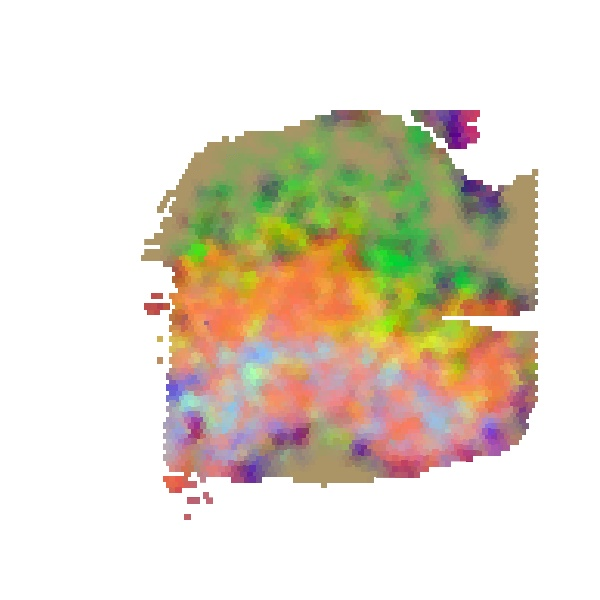

### S10_scGNN_RPKM_PEalpha0.2_zdim256.png

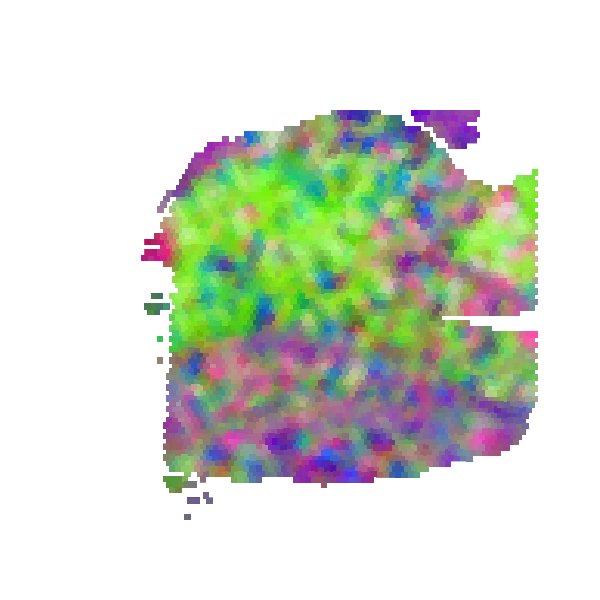

### S10_scGNN_scran_PEalpha0.1_zdim16.png

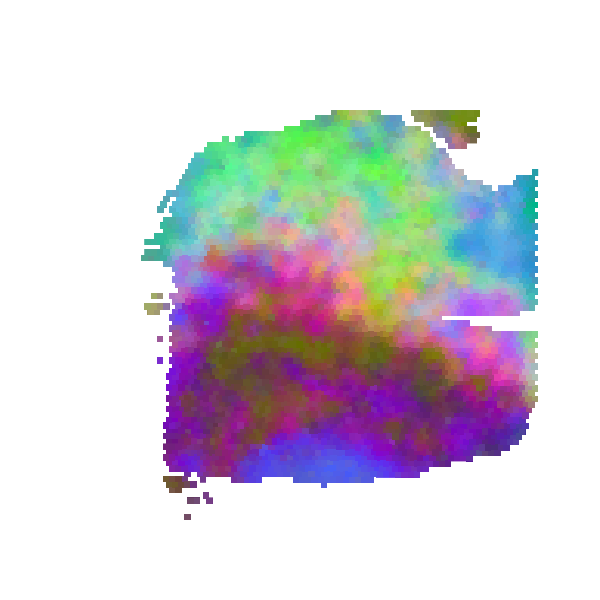

### S10_scGNN_sctransform_PEalpha0.5_zdim128.png

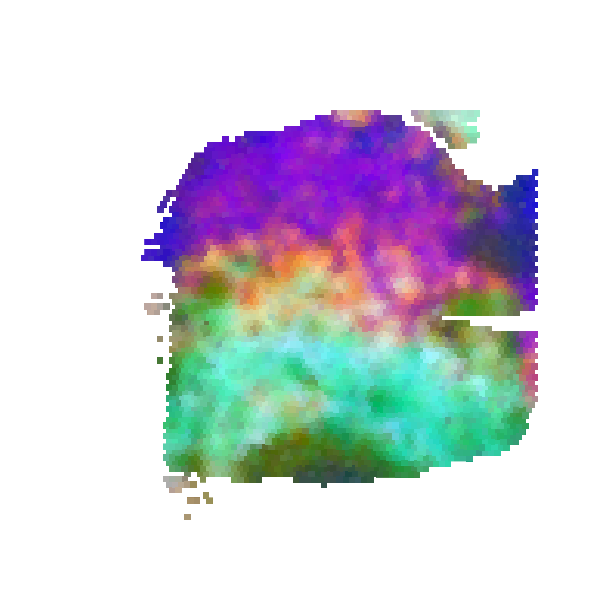

### S11_scGNN_expression_PEalpha1.5_zdim3.png

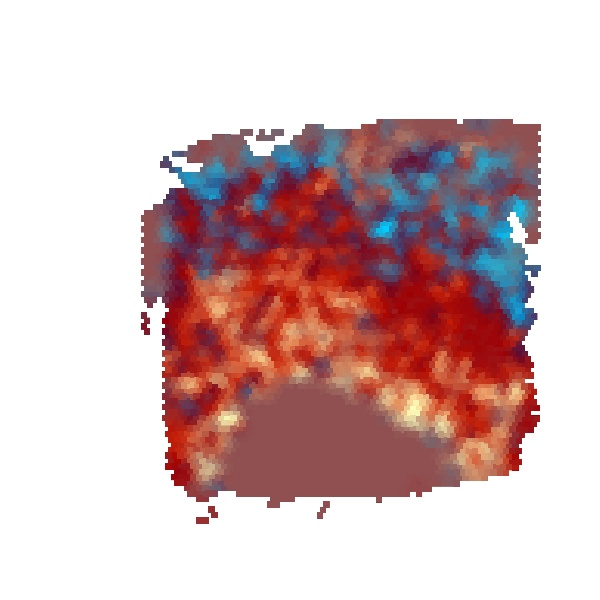

### S11_scGNN_RPKM_PEalpha1.5_zdim128.png

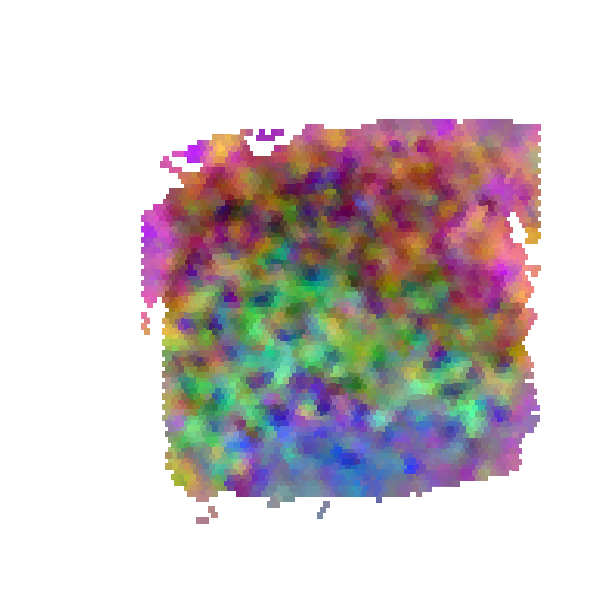

### S11_scGNN_scran_PEalpha0.1_zdim3.png

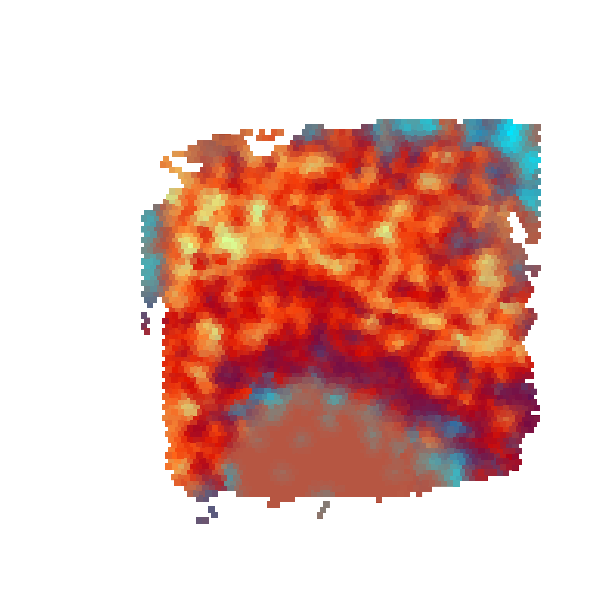

### S11_scGNN_sctransform_PEalpha0.3_zdim256.png

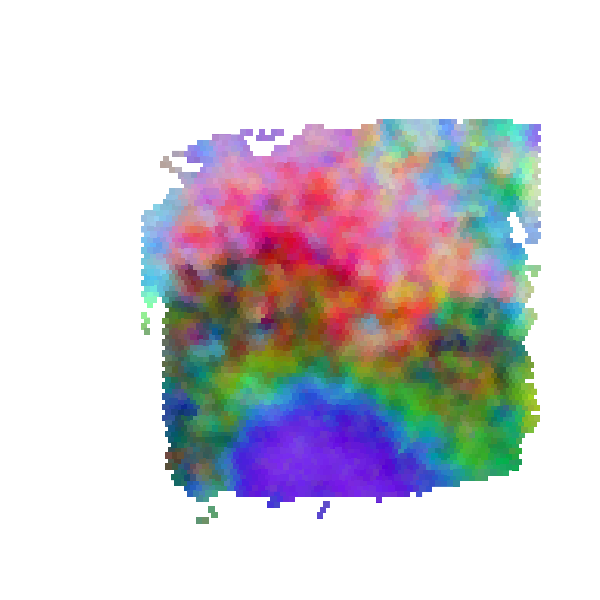

### S12_scGNN_expression_PEalpha1.5_zdim128.png

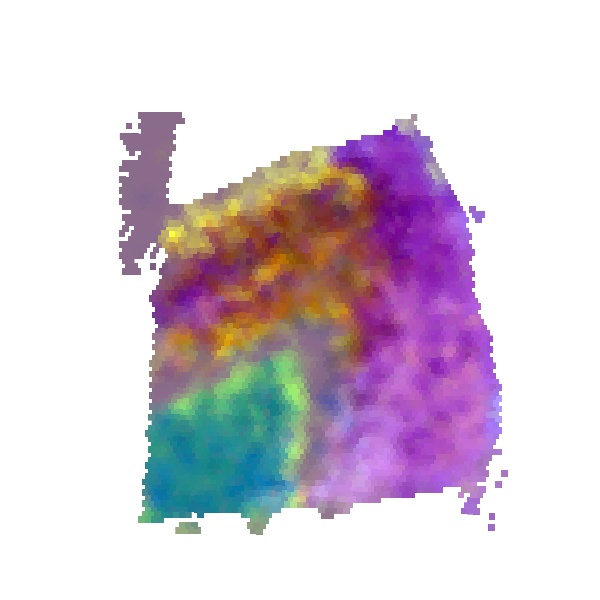

### S12_scGNN_RPKM_PEalpha1.2_zdim10.png

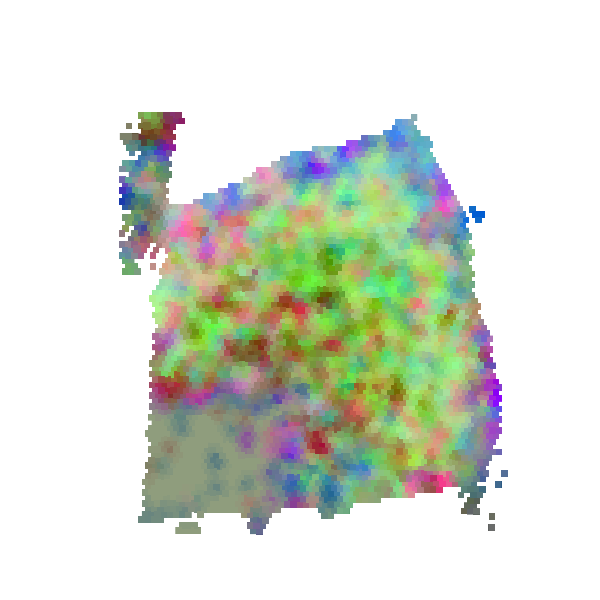

### S12_scGNN_scran_PEalpha0.1_zdim128.png

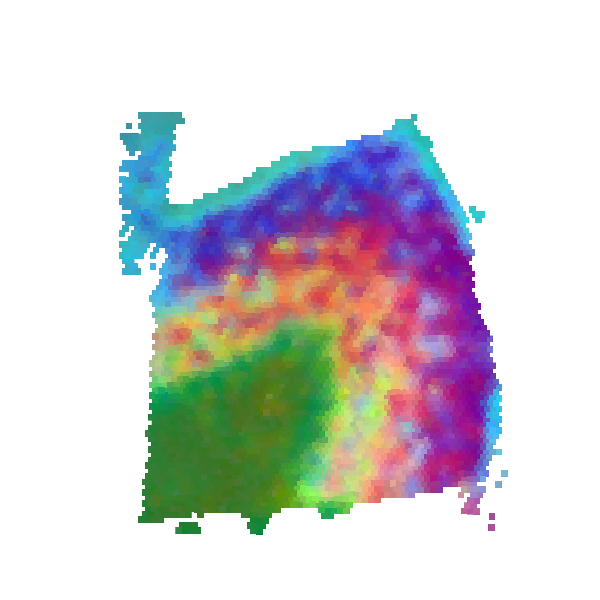

### S12_scGNN_sctransform_PEalpha1.2_zdim256.png

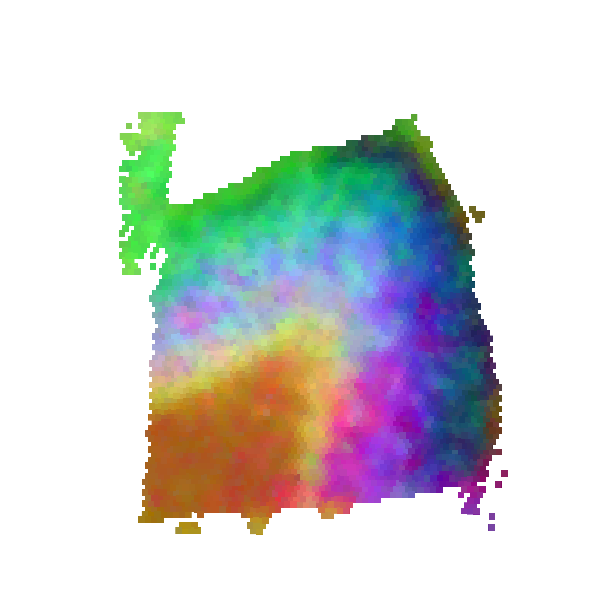

### S13_scGNN_expression_PEalpha1.0_zdim3.png

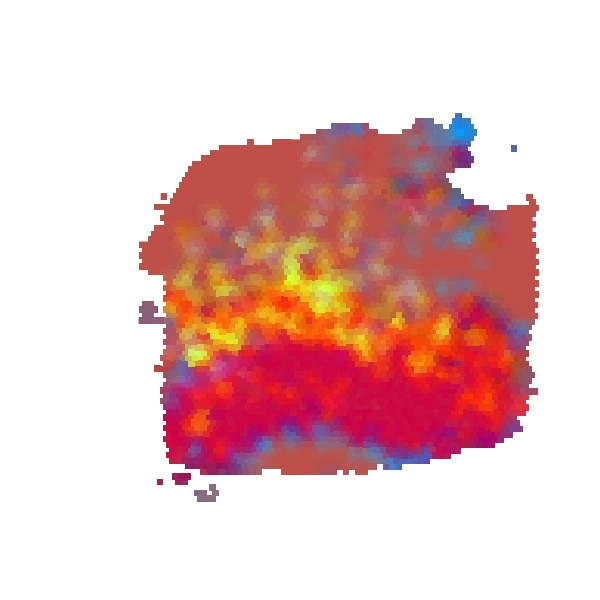

### S13_scGNN_RPKM_PEalpha1.0_zdim64.png

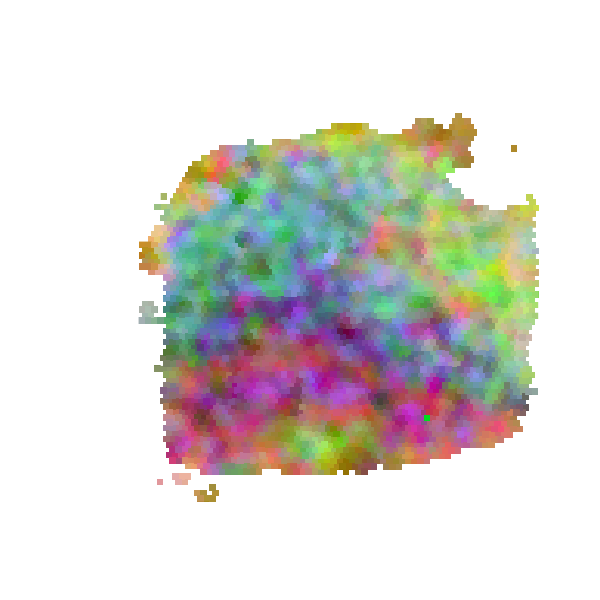

### S13_scGNN_scran_PEalpha0.3_zdim256.png

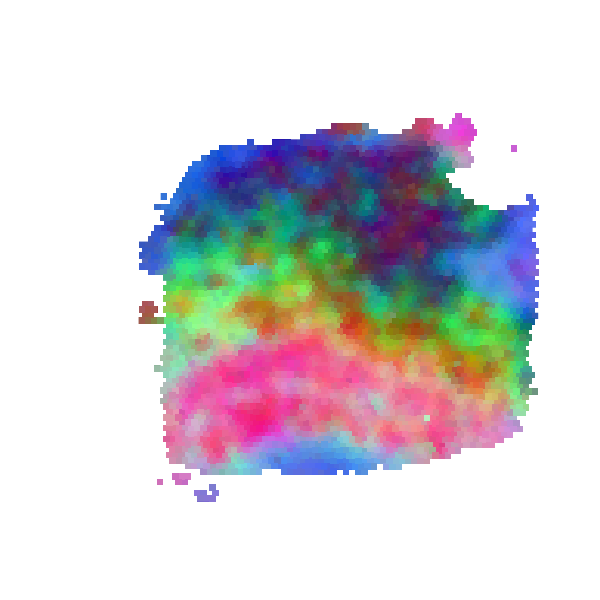

### S13_scGNN_sctransform_PEalpha0.2_zdim10.png

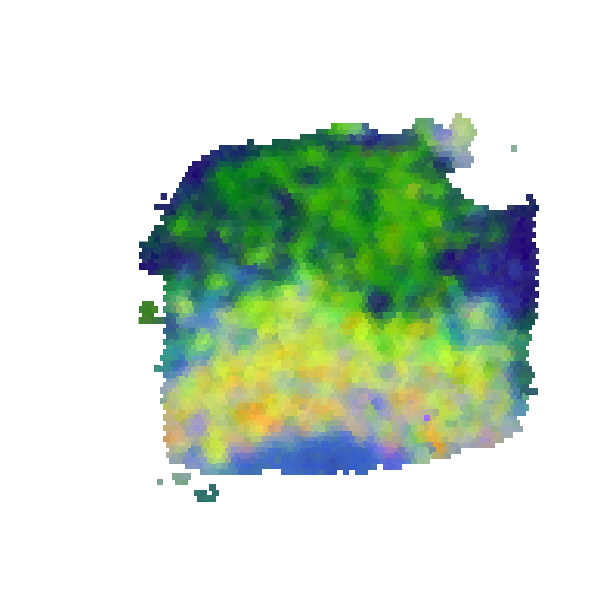

### S14_scGNN_expression_PEalpha0.2_zdim32.png

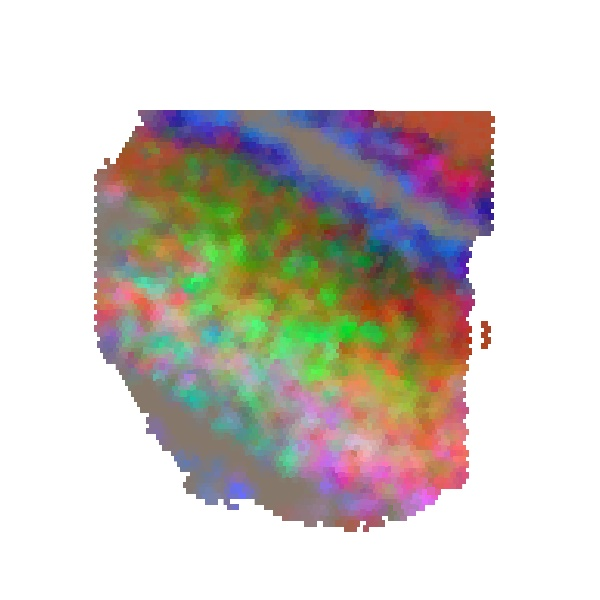

### S14_scGNN_RPKM_PEalpha0.5_zdim3.png

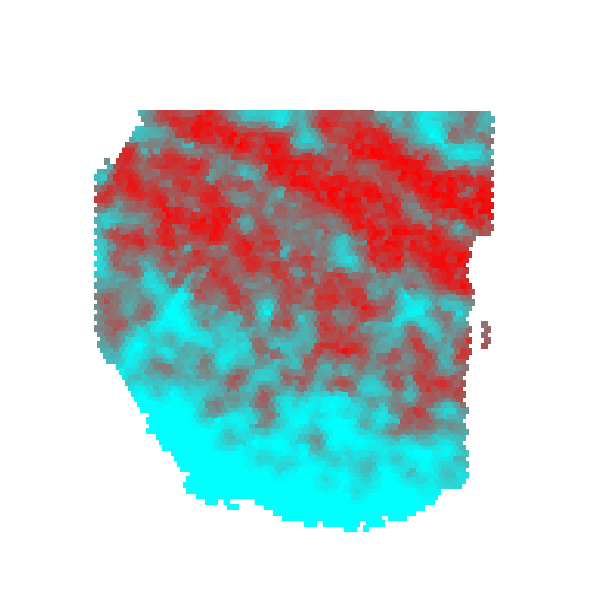

### S14_scGNN_scran_PEalpha1.5_zdim16.png

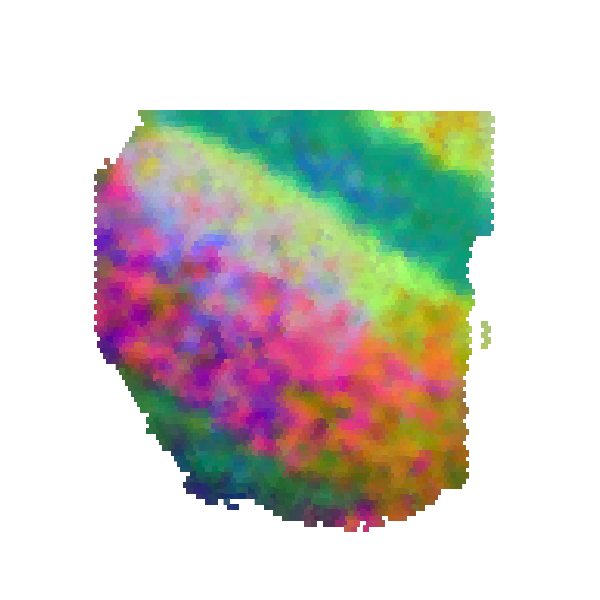

### S14_scGNN_sctransform_PEalpha0.2_zdim128.png

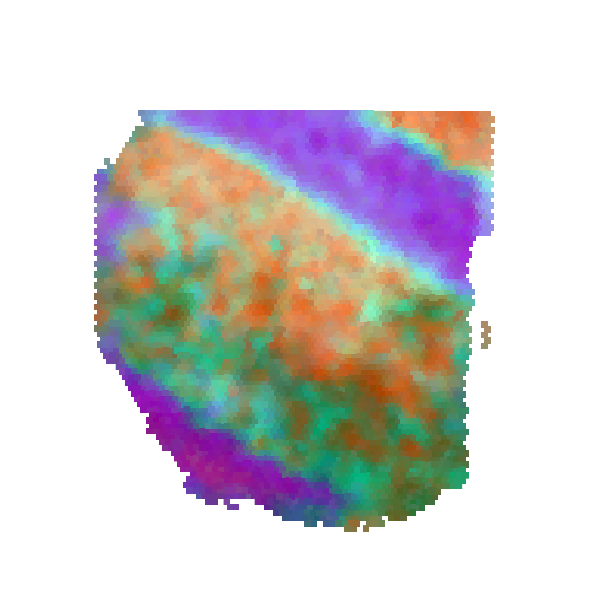

### S15_scGNN_expression_PEalpha0.3_zdim128.png

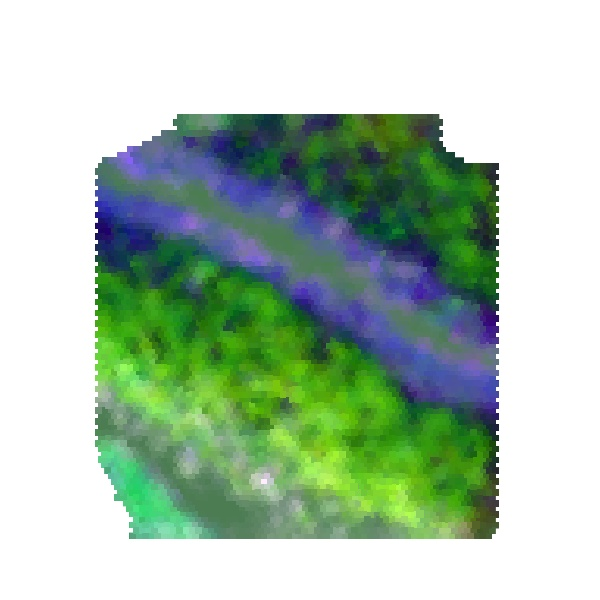

### S15_scGNN_RPKM_PEalpha1.0_zdim16.png

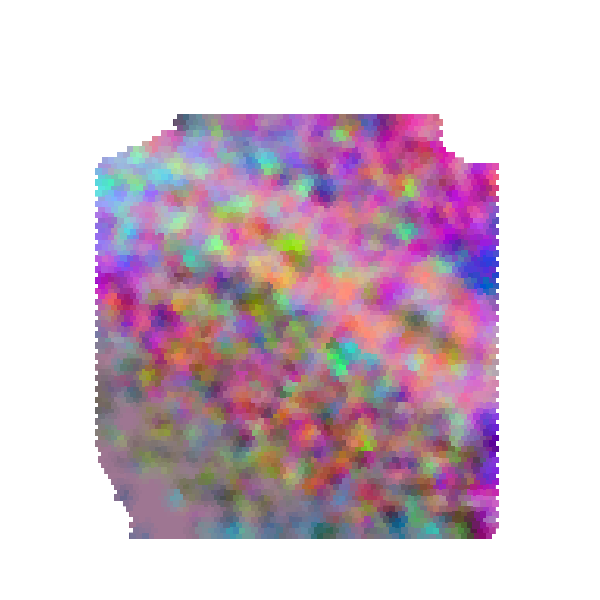

### S15_scGNN_scran_PEalpha0.2_zdim10.png

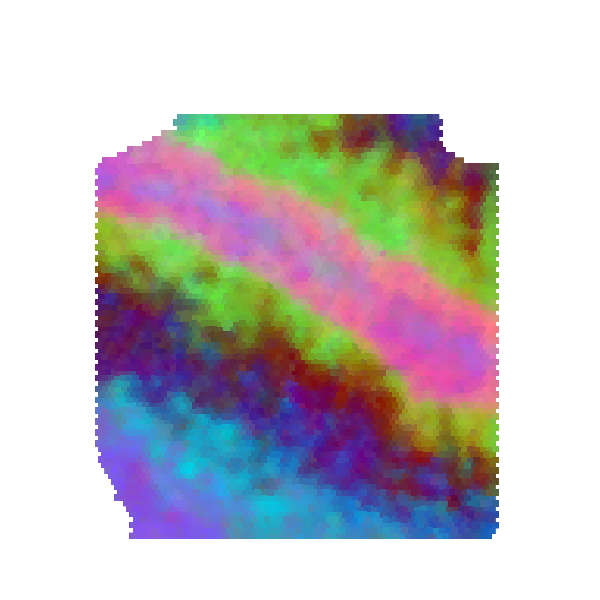

### S15_scGNN_sctransform_PEalpha0.1_zdim128.png

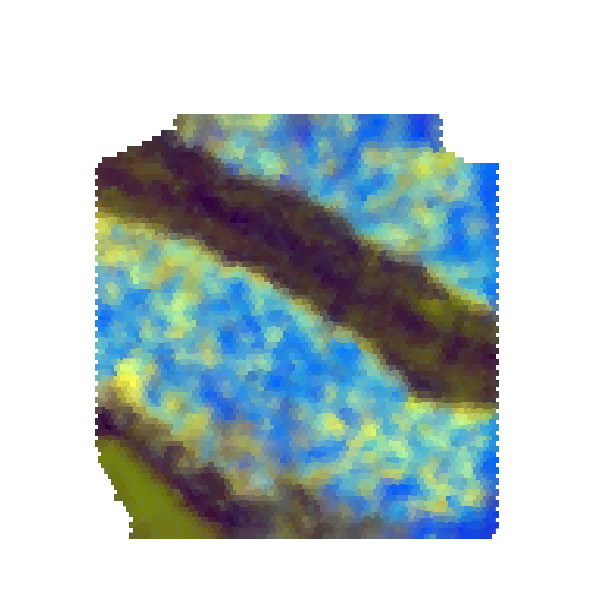

### S16_scGNN_expression_PEalpha1.2_zdim64.png

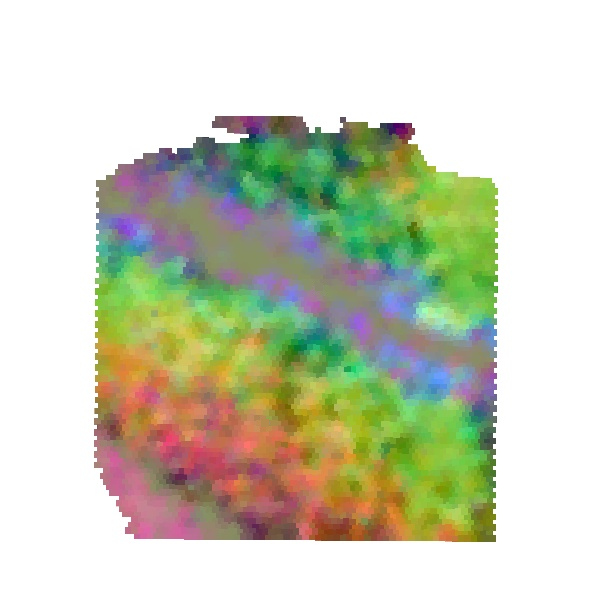

### S16_scGNN_RPKM_PEalpha1.2_zdim10.png

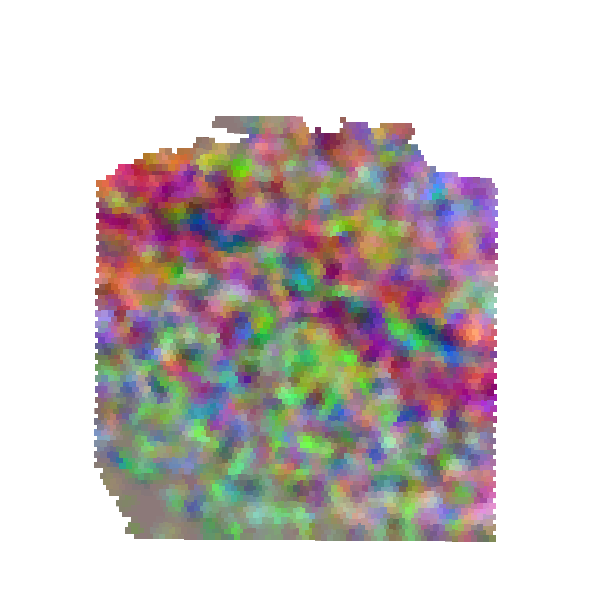

### S16_scGNN_scran_PEalpha0.5_zdim32.png

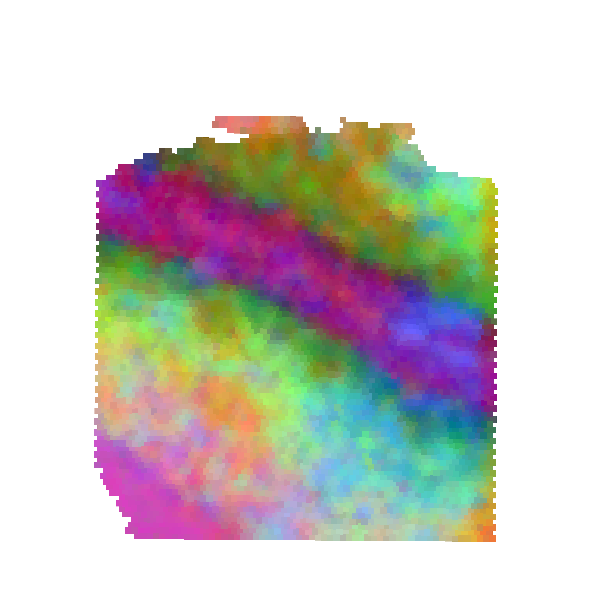

### S16_scGNN_sctransform_PEalpha2.0_zdim64.png

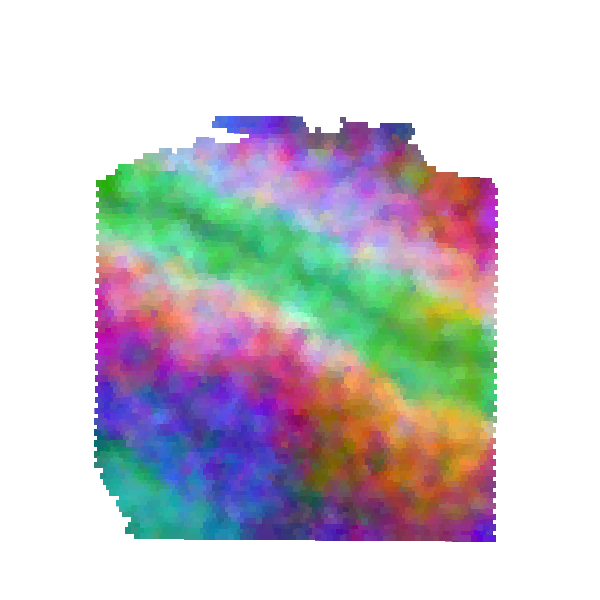

### S17_scGNN_expression_PEalpha1.2_zdim256.png

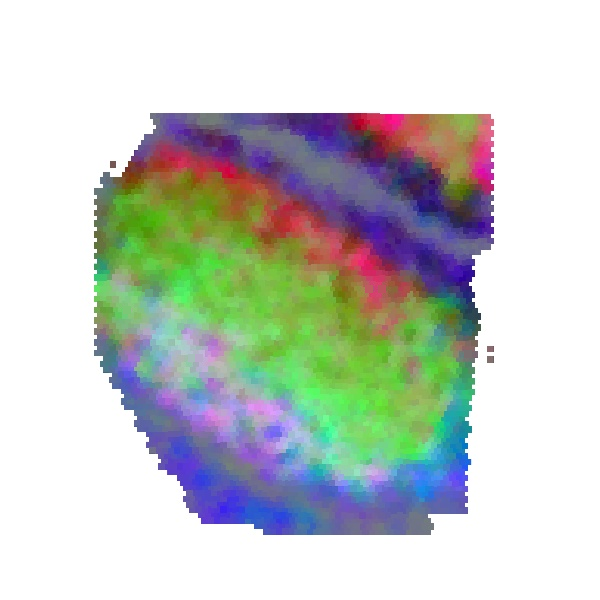

### S17_scGNN_RPKM_PEalpha1.5_zdim10.png

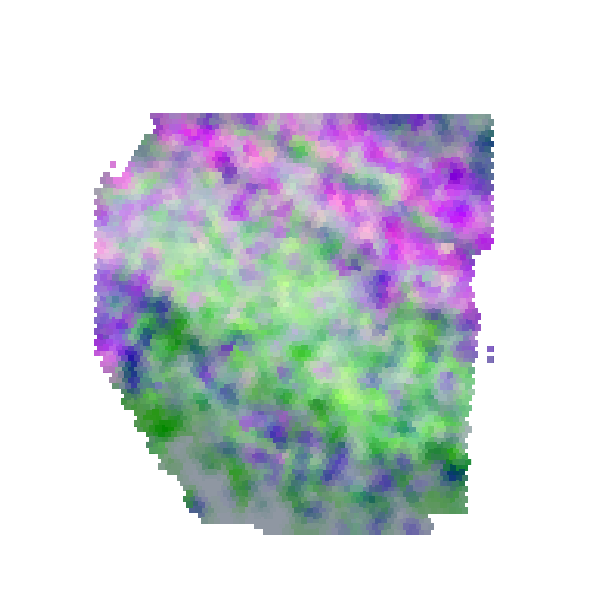

### S17_scGNN_scran_PEalpha0.1_zdim32.png

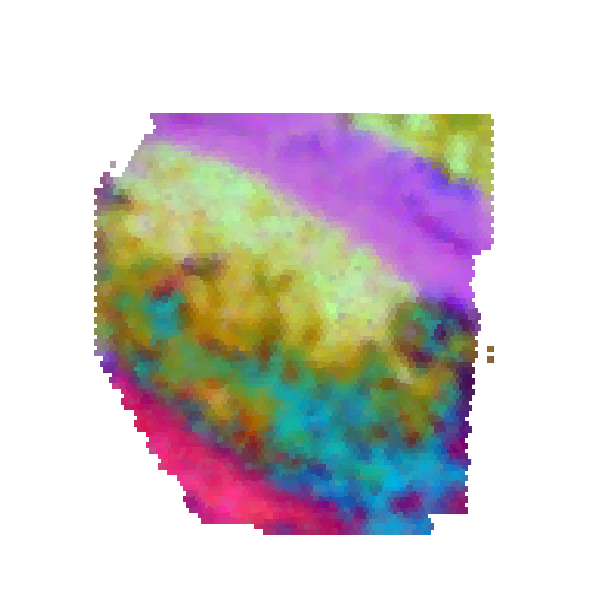

### S17_scGNN_sctransform_PEalpha2.0_zdim10.png

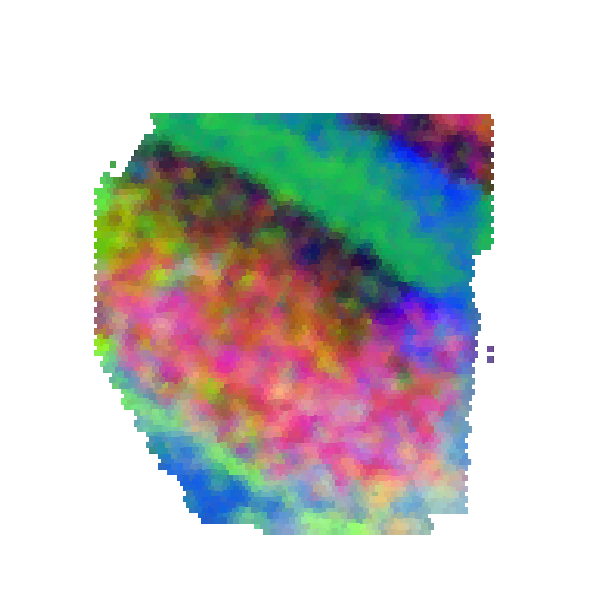
