## Supplementary figures and images for "Define and visualize pathological architectures of human tissues from spatially resolved transcriptomics using deep learning"

### S2_6_tools_RPKM.jpeg

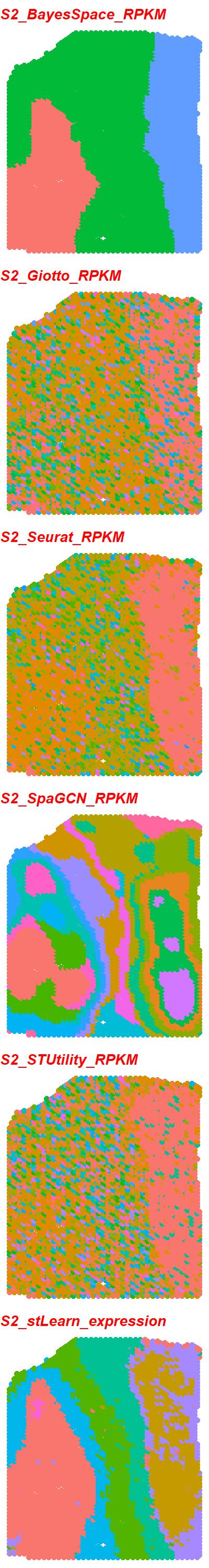

### S2_6_tools_scran.jpeg

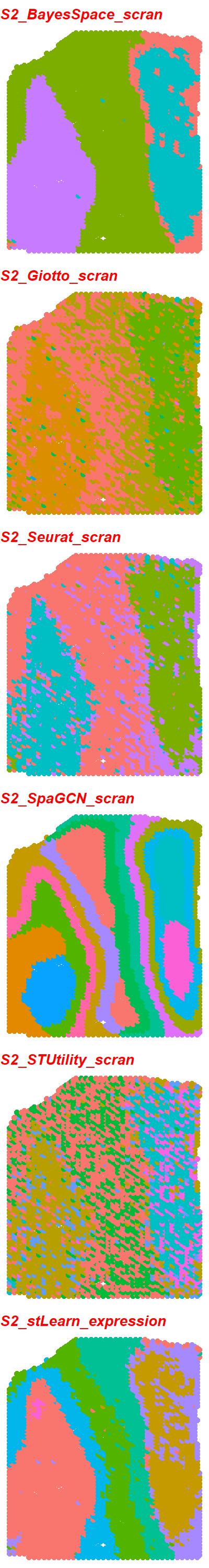

### S2_6_tools_scTransform.jpeg

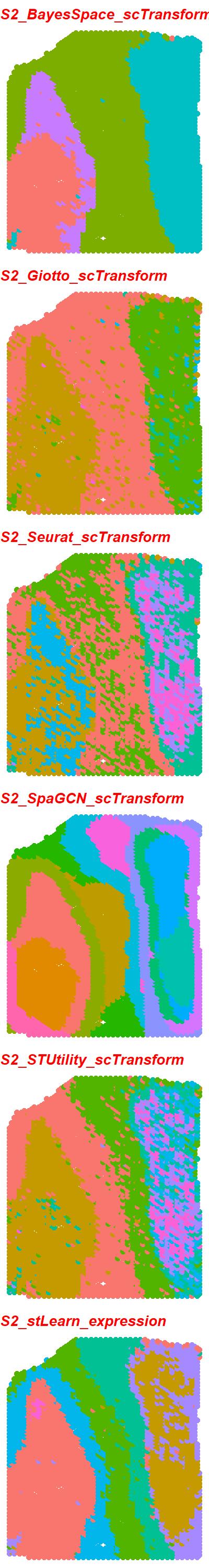

### S2_6_tools_TMM.jpeg

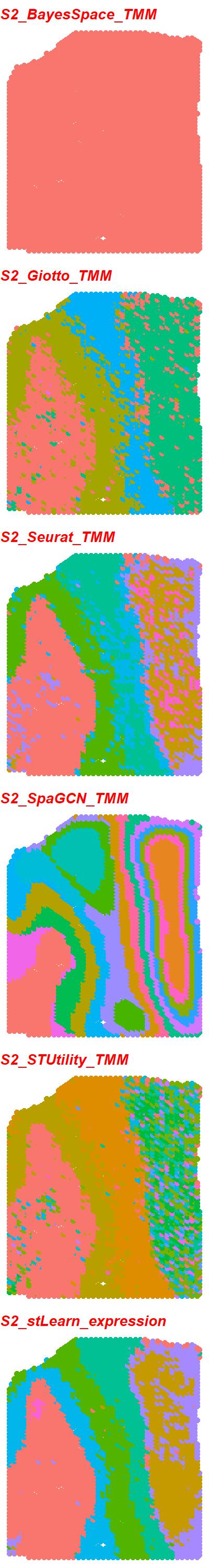

### S2_6_tools_TPM.jpeg

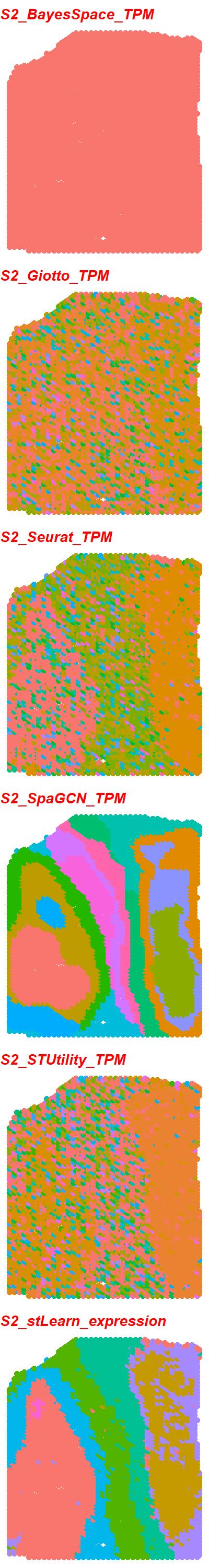

### S2_scGNN_DESeq2_PEalpha1.5_zdim10.png

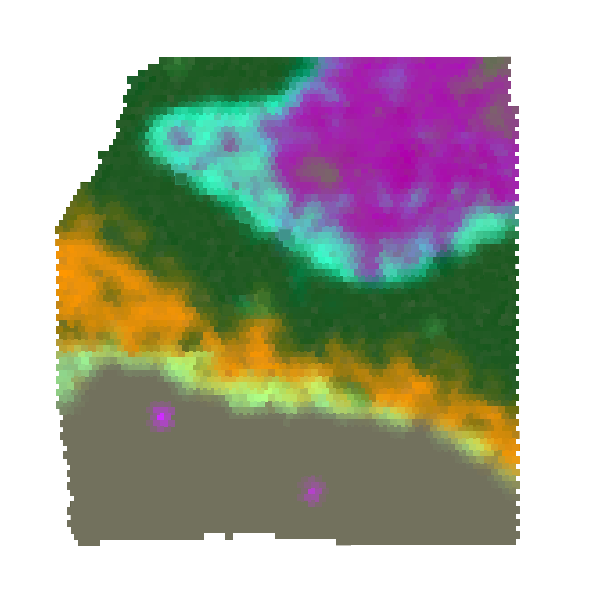

### S3_6_tools_RPKM.jpeg

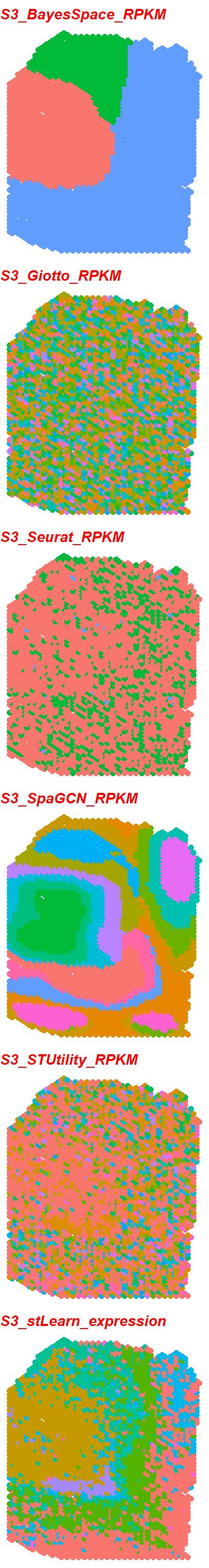

### S3_6_tools_scran.jpeg

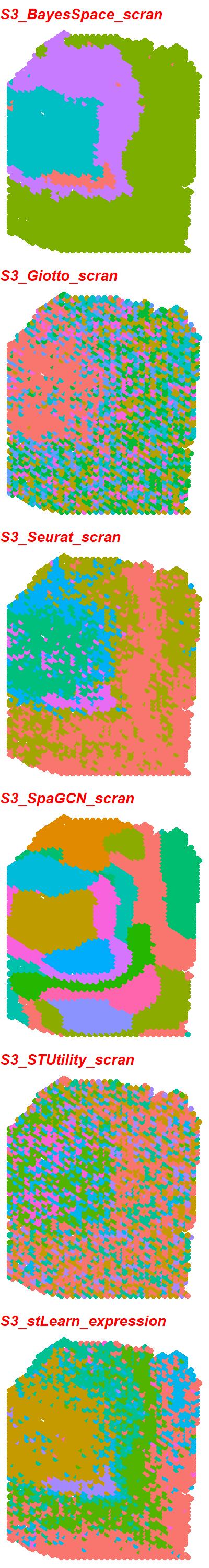

### S3_6_tools_scTransform.jpeg

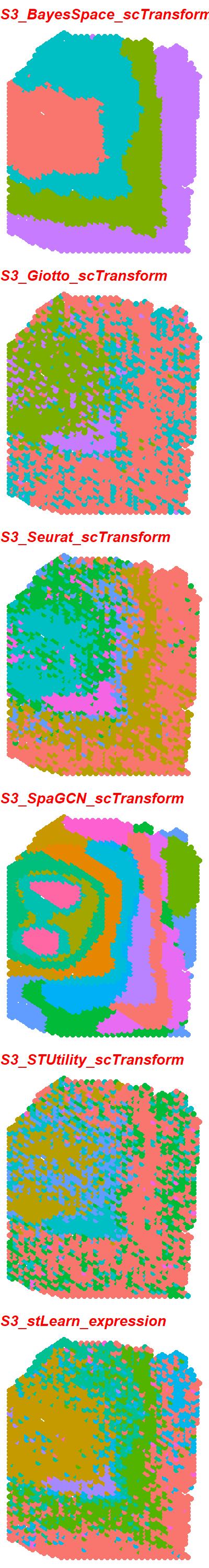

### S3_6_tools_TMM.jpeg

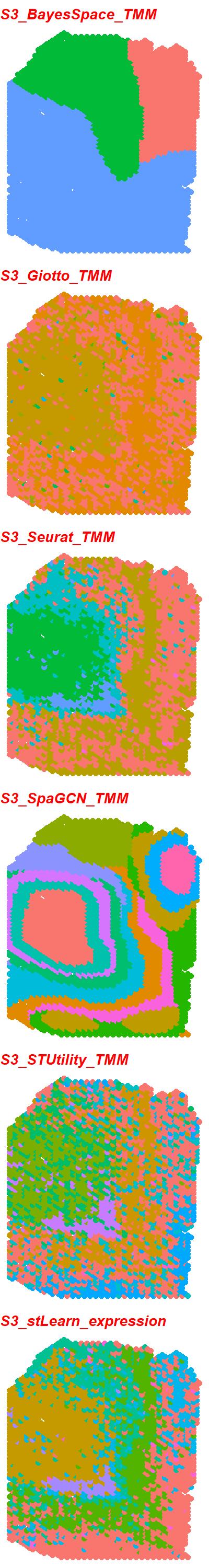

### S3_6_tools_TPM.jpeg

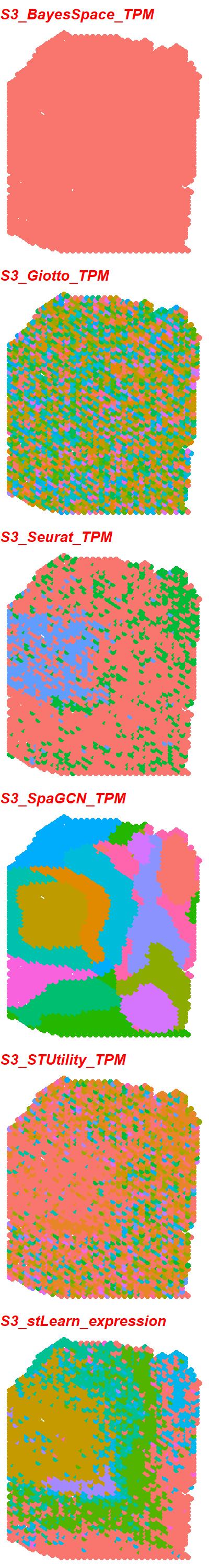

### S3_scGNN_DESeq2_PEalpha1.2_zdim16.png

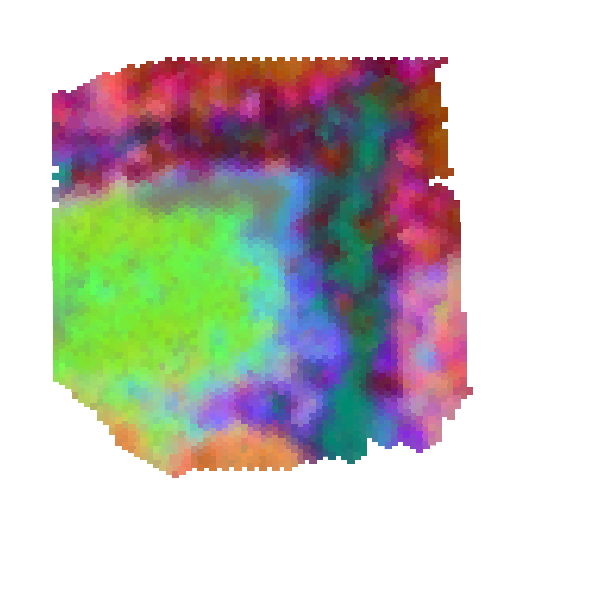

### S4_6_tools_LogCPM.jpeg

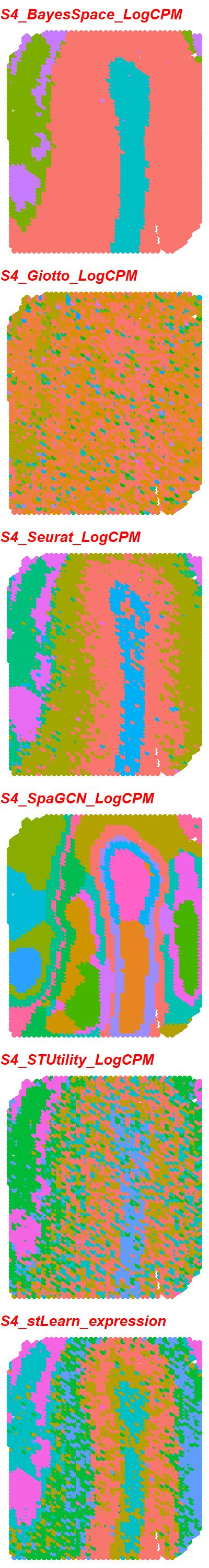

### S4_6_tools_RPKM.jpeg

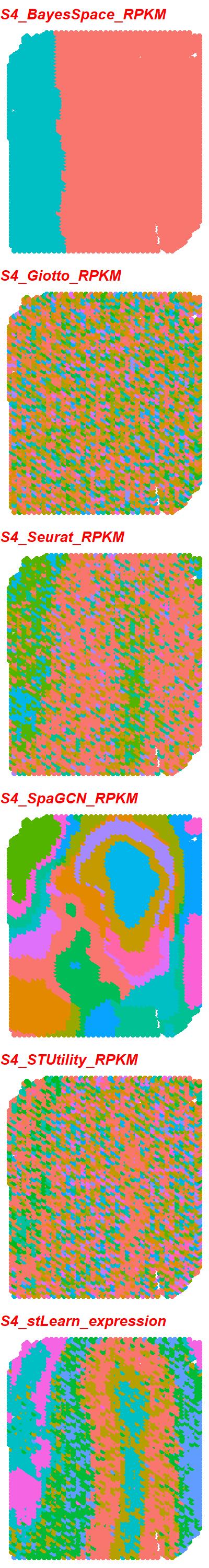

### S4_6_tools_scran.jpeg

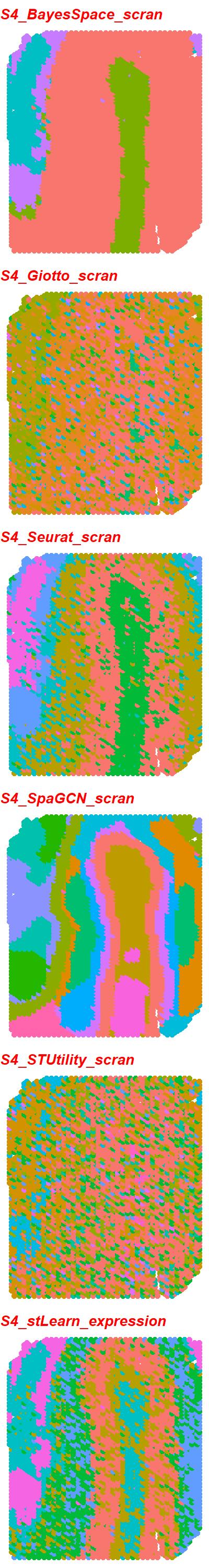

### S4_6_tools_scTransform.jpeg

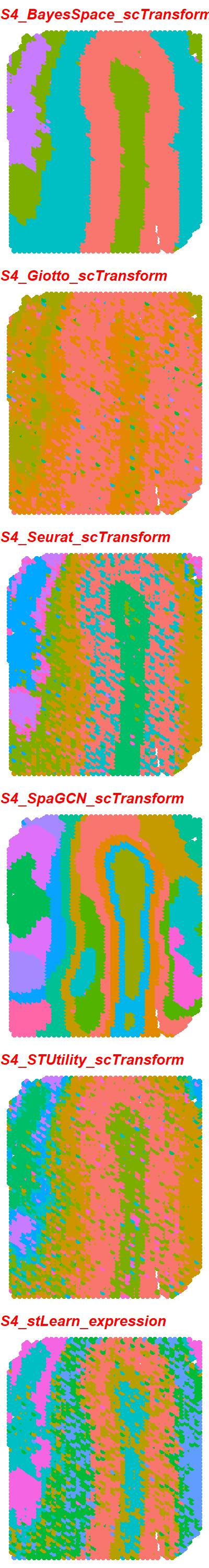

### S4_6_tools_TMM.jpeg

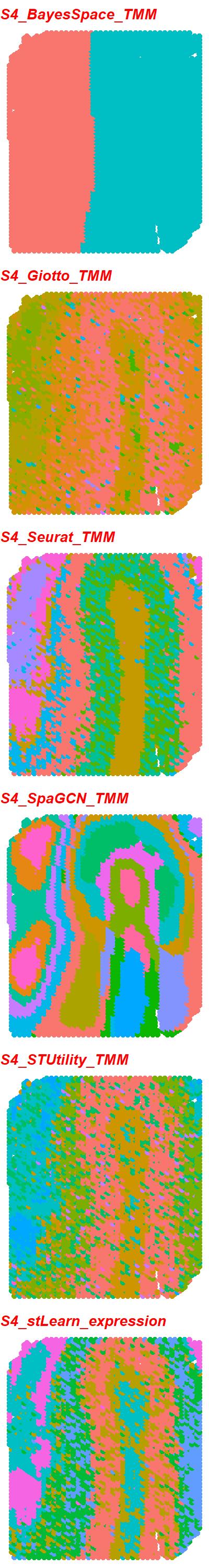

### S4_6_tools_TPM.jpeg

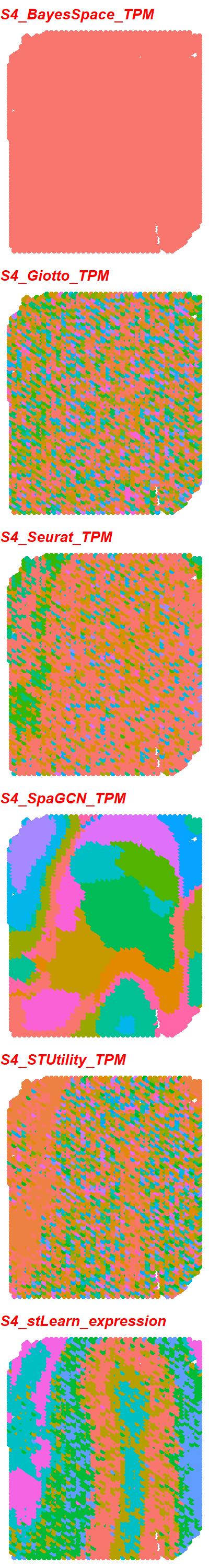

### S4_scGNN_DESeq2_PEalpha0.2_zdim16.png

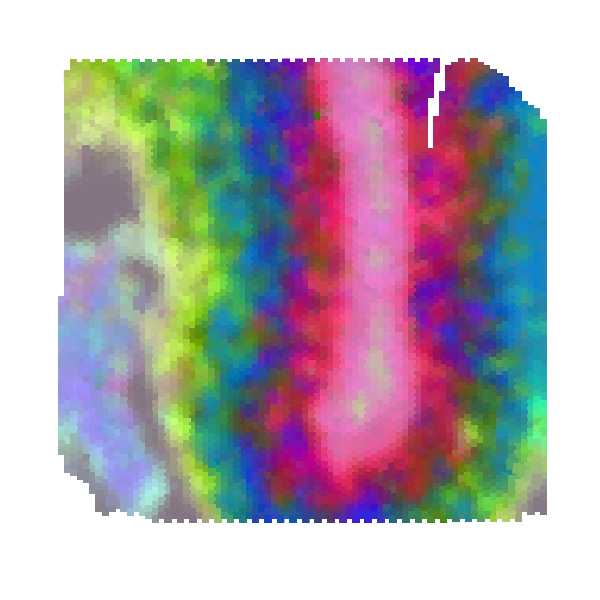

### S5_6_tools_LogCPM.jpeg

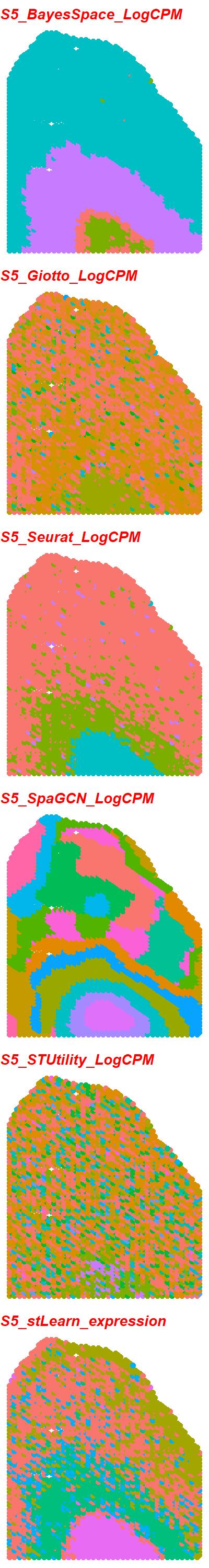

### S5_6_tools_RPKM.jpeg

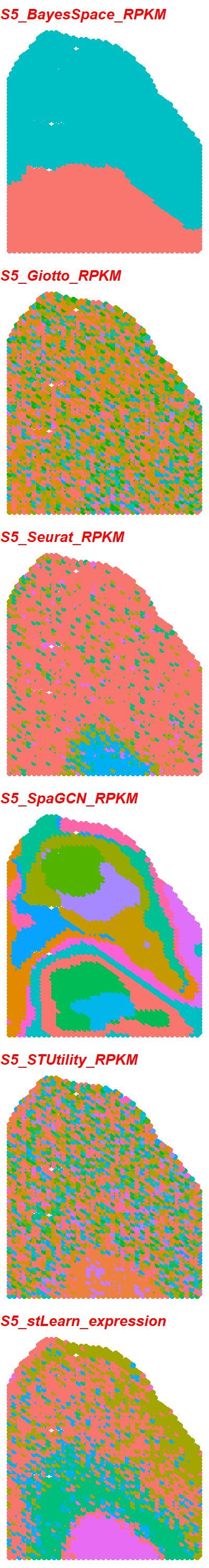

### S5_6_tools_scran.jpeg

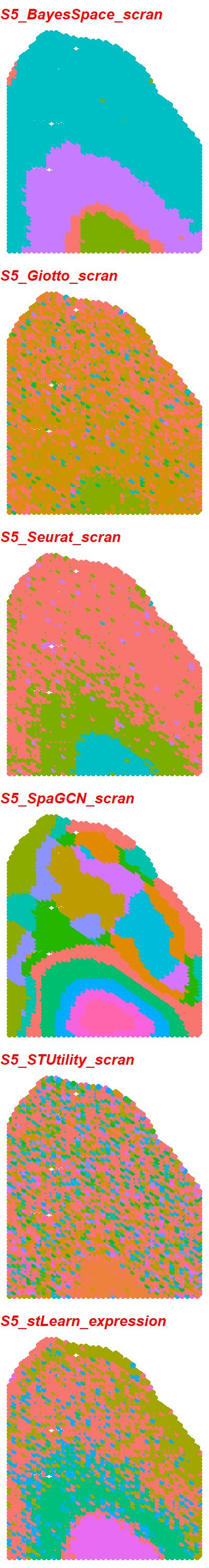

### S5_6_tools_scTransform.jpeg

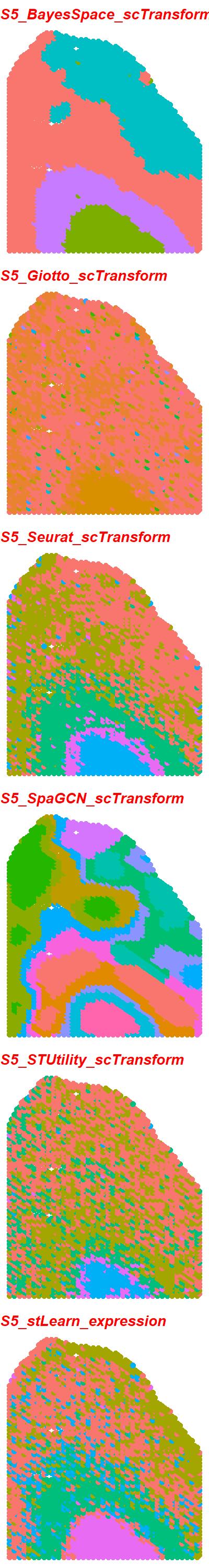

### S5_6_tools_TMM.jpeg

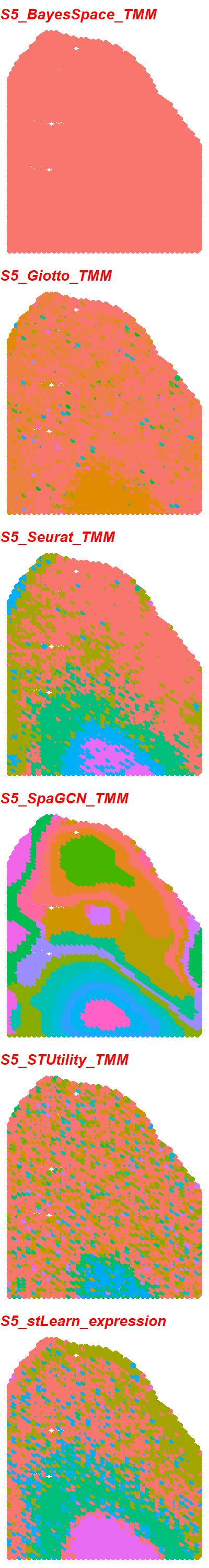

### S5_6_tools_TPM.jpeg

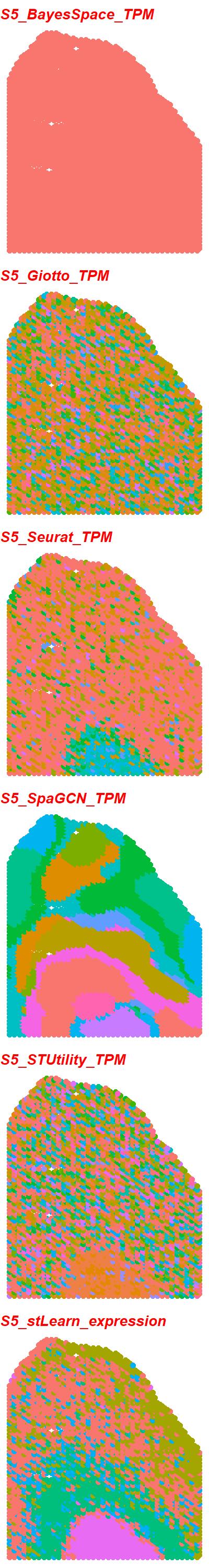

### S5_scGNN_DESeq2_PEalpha1.2_zdim128.png

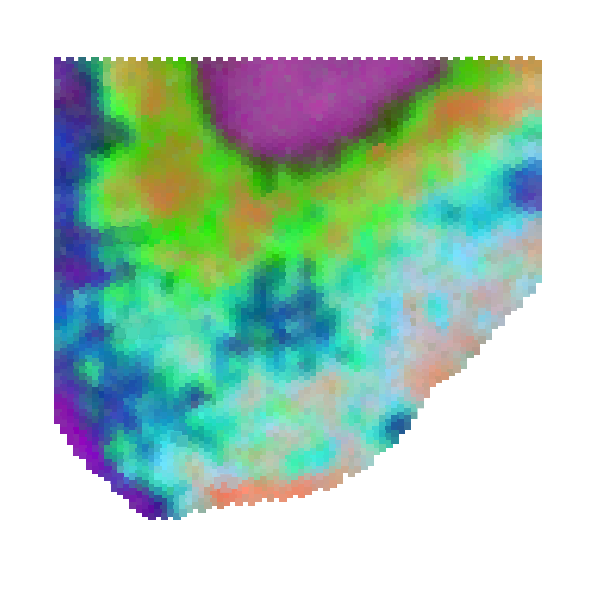

### S6_6_tools_LogCPM.jpeg

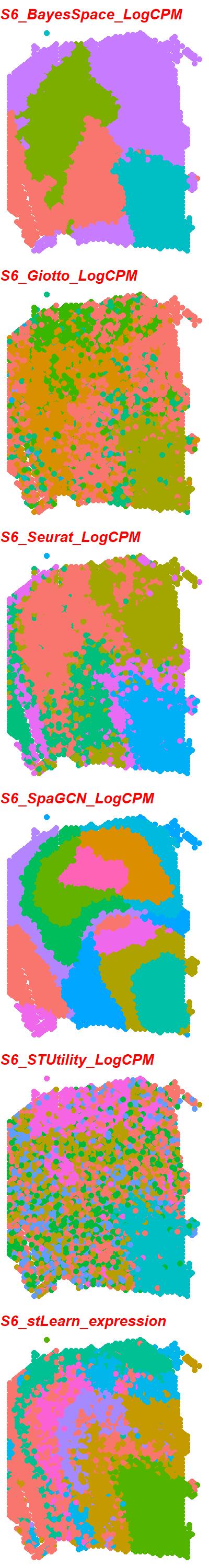

### S6_6_tools_RPKM.jpeg

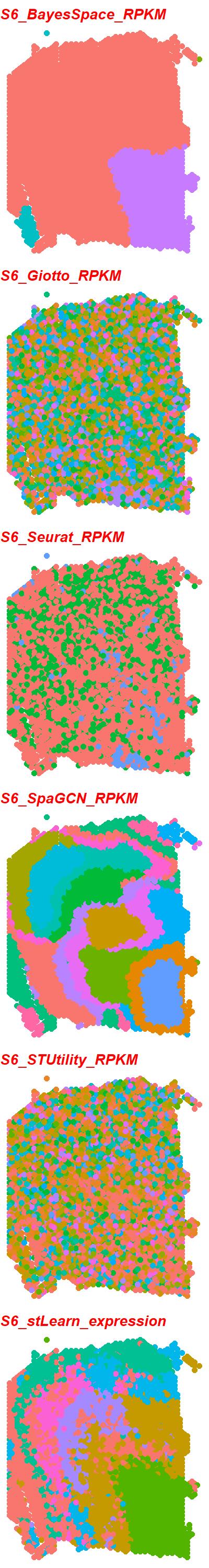

### S6_6_tools_scran.jpeg

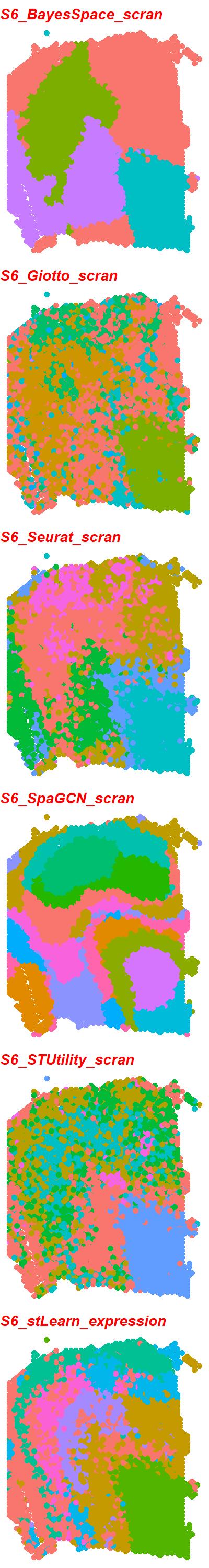

### S6_6_tools_scTransform.jpeg

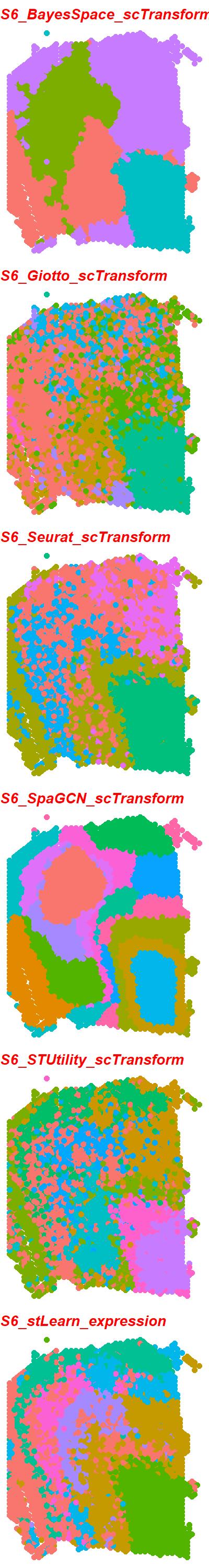

### S6_6_tools_TMM.jpeg

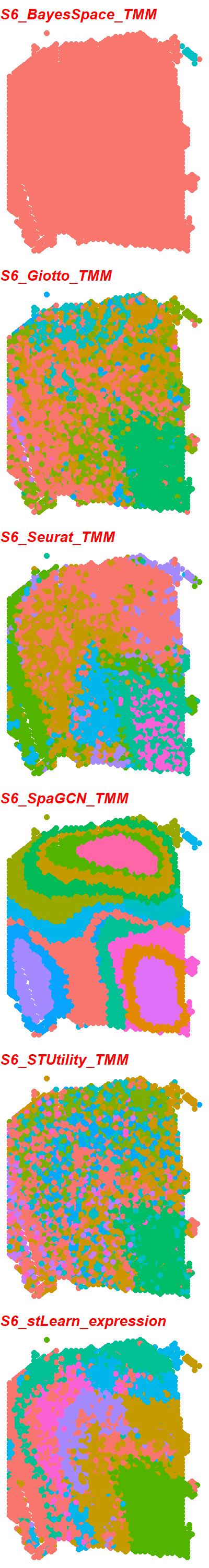

### S6_6_tools_TPM.jpeg

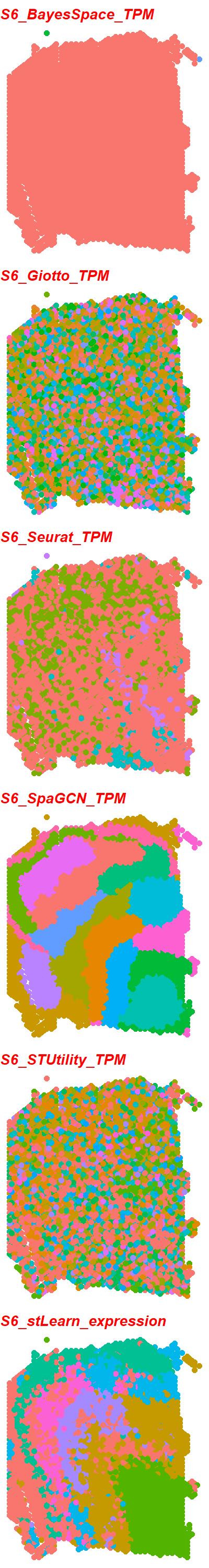

### S6_scGNN_DESeq2_PEalpha1.2_zdim32.png

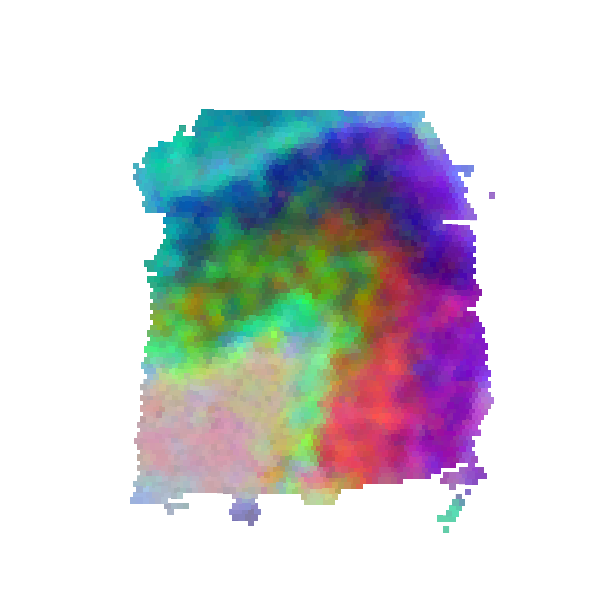

### S7_6_tools_LogCPM.jpeg

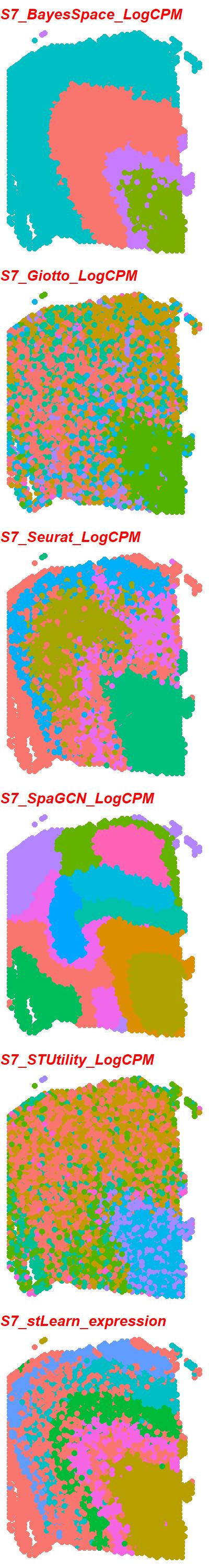

### S7_6_tools_RPKM.jpeg

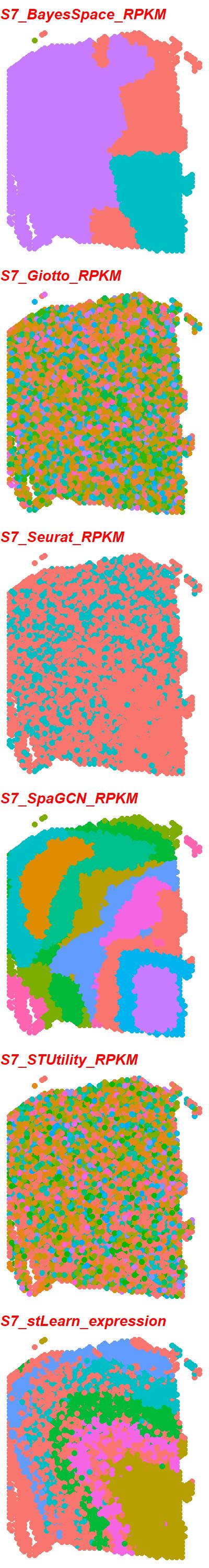

### S7_6_tools_scran.jpeg

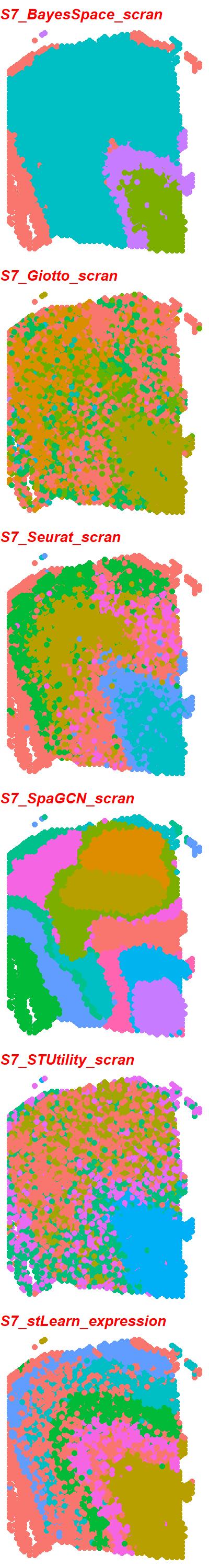

### S7_6_tools_scTransform.jpeg

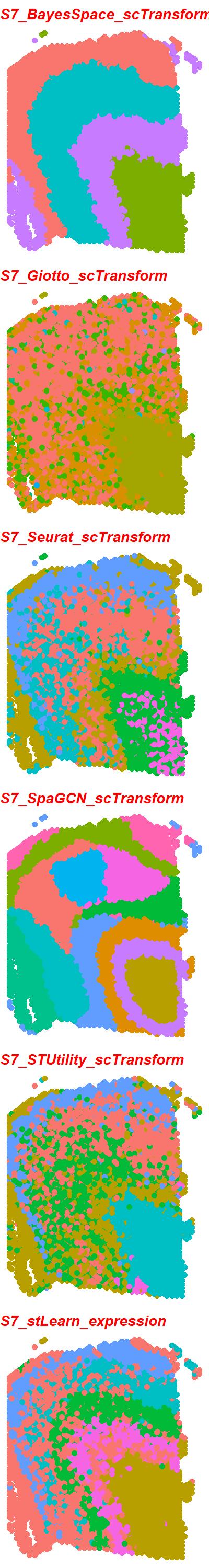

### S7_6_tools_TMM.jpeg

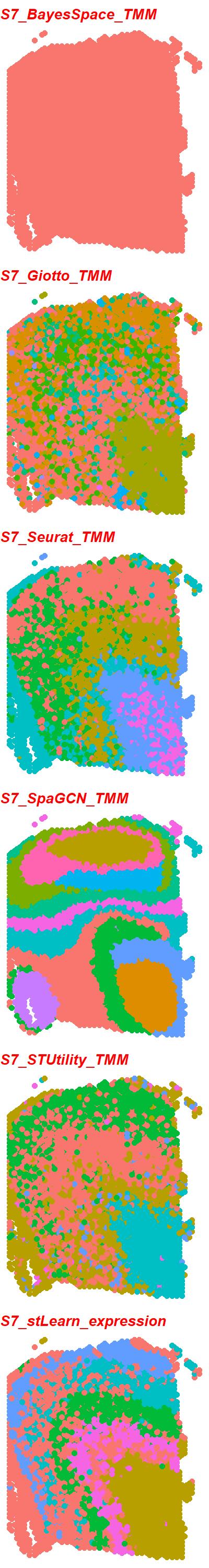

### S7_6_tools_TPM.jpeg

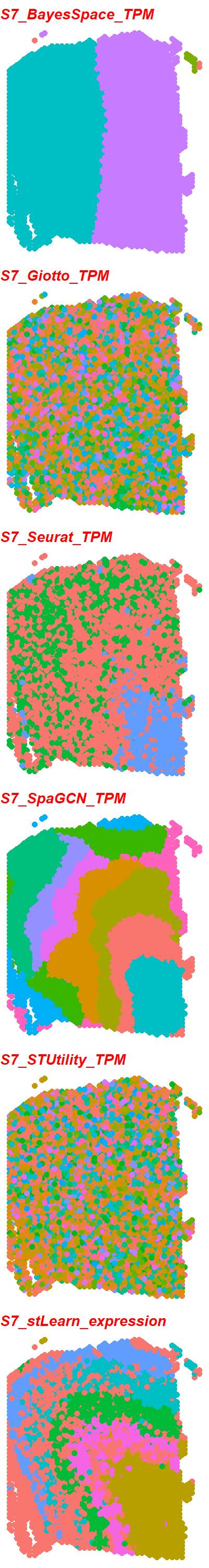

### S7_scGNN_DESeq2_PEalpha0.5_zdim16.png

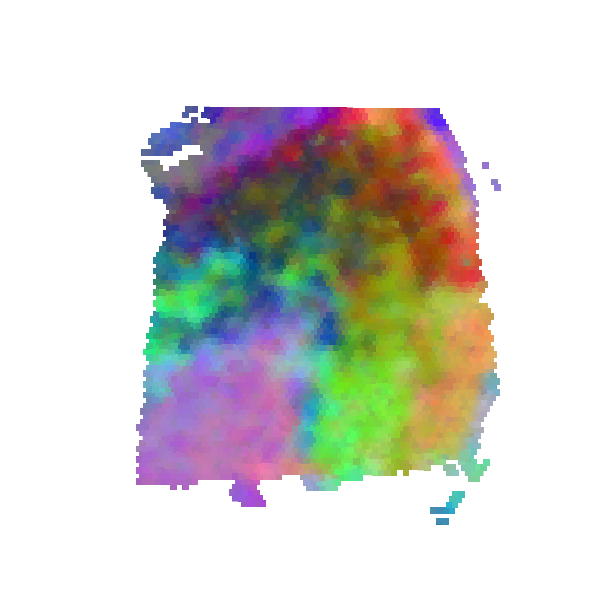

### S8_6_tools_LogCPM.jpeg

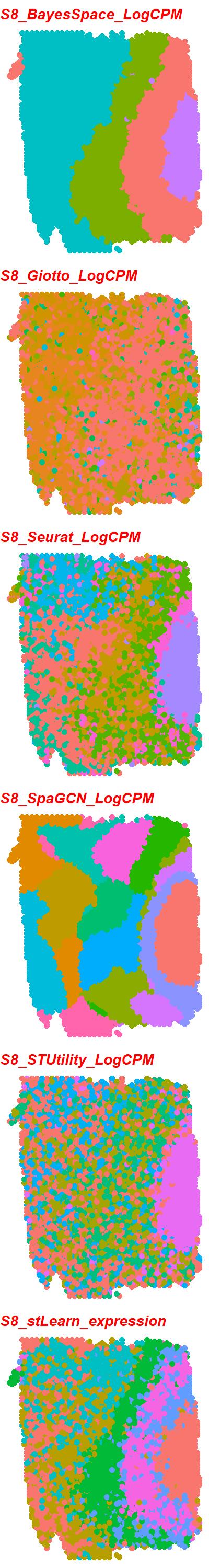

### S8_6_tools_RPKM.jpeg

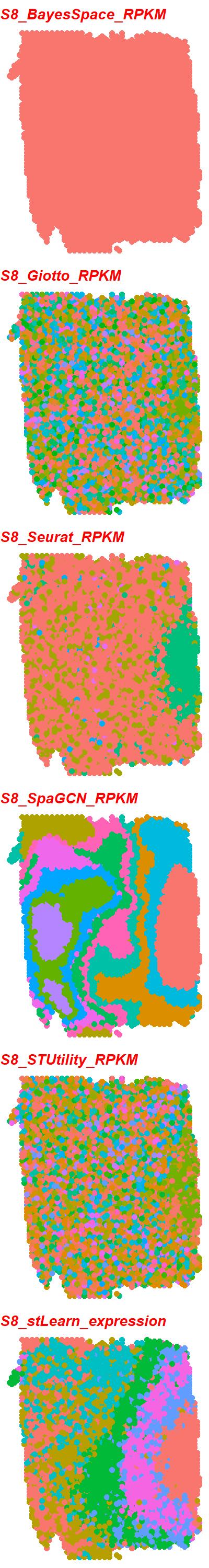

### S8_6_tools_scran.jpeg

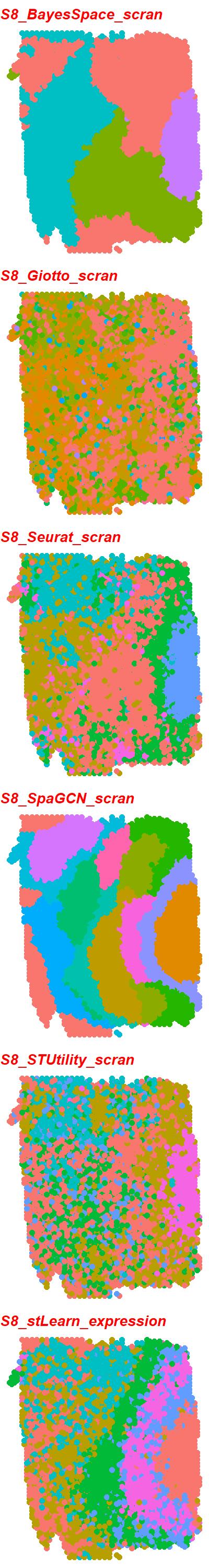

### S8_6_tools_scTransform.jpeg

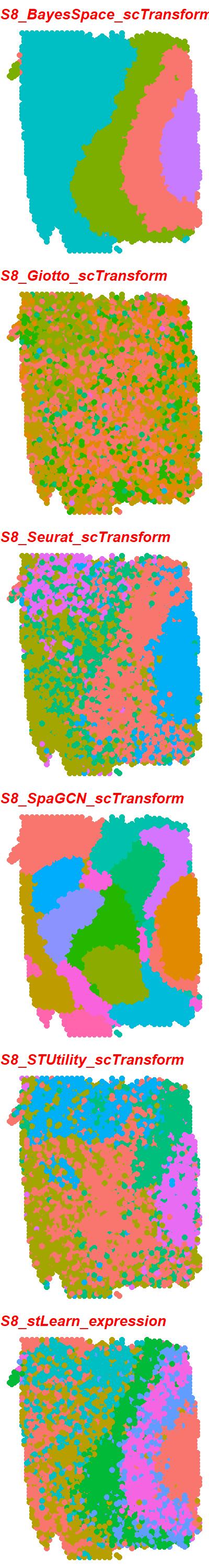

### S8_6_tools_TMM.jpeg

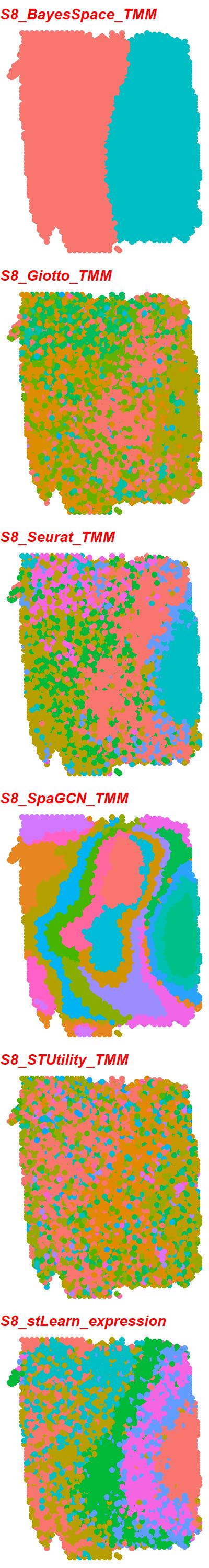

### S8_6_tools_TPM.jpeg

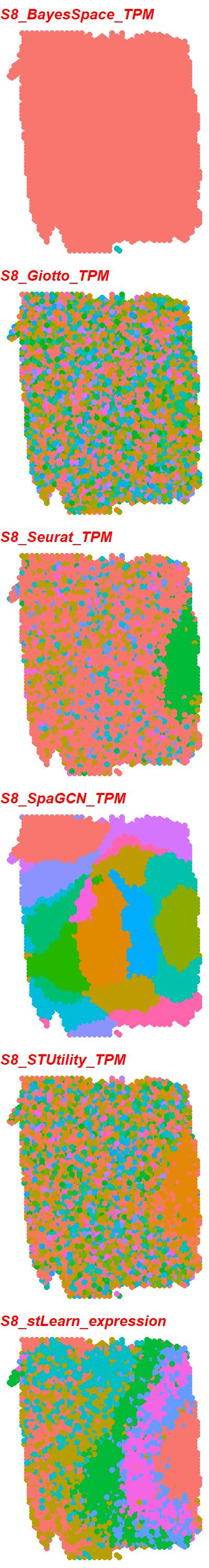

### S9_6_tools_LogCPM.jpeg

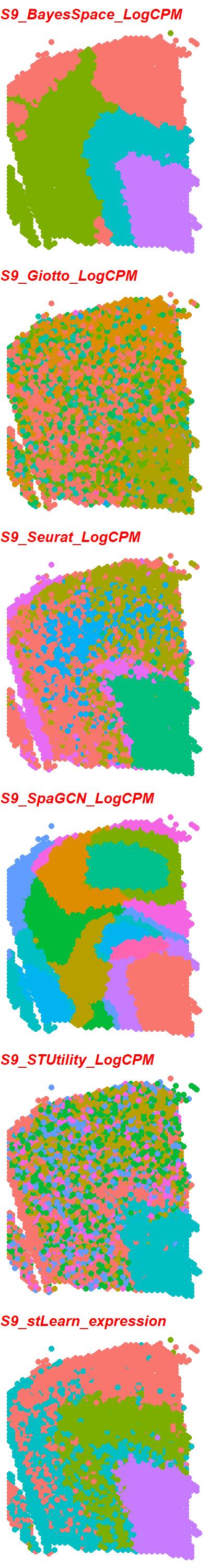

### S9_6_tools_RPKM.jpeg

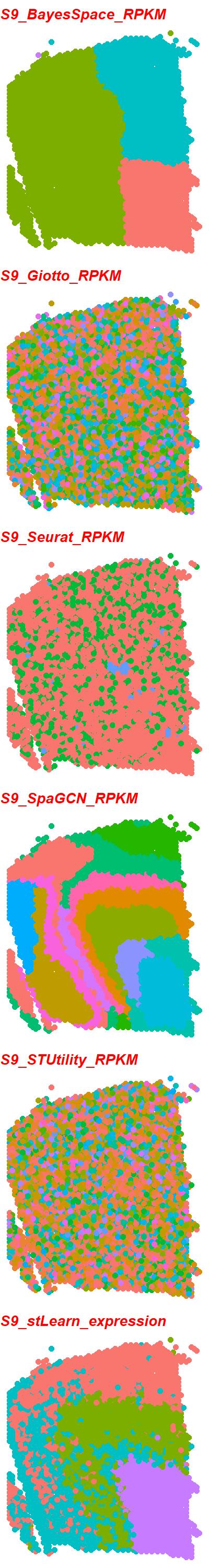

### S9_6_tools_scran.jpeg

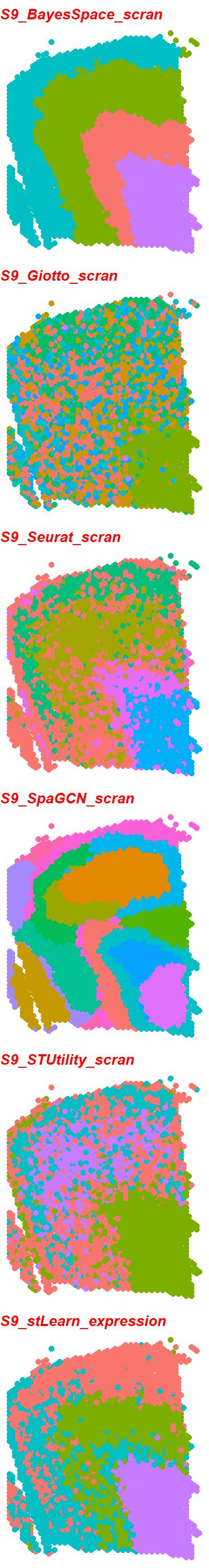

### S9_6_tools_scTransform.jpeg

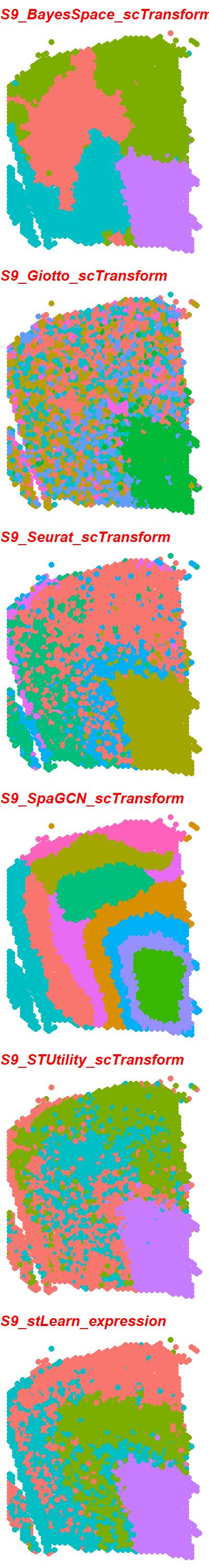

### S9_6_tools_TMM.jpeg

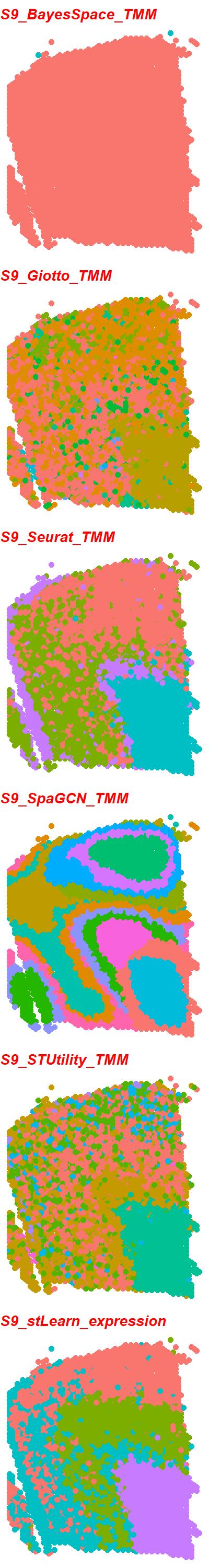

### S9_6_tools_TPM.jpeg

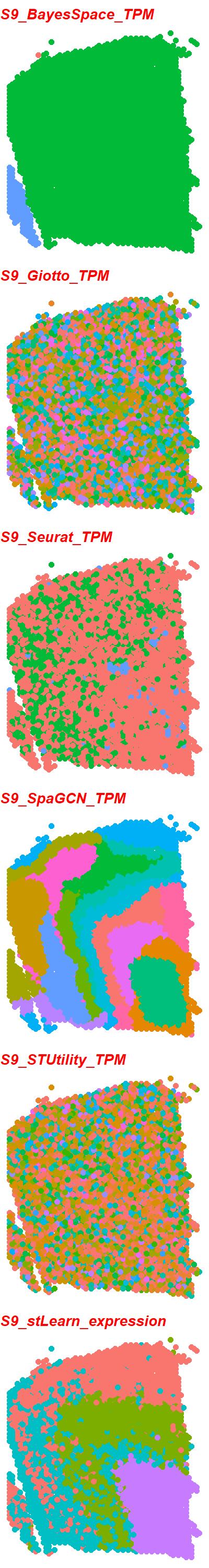

### S10_6_tools_RPKM.jpeg

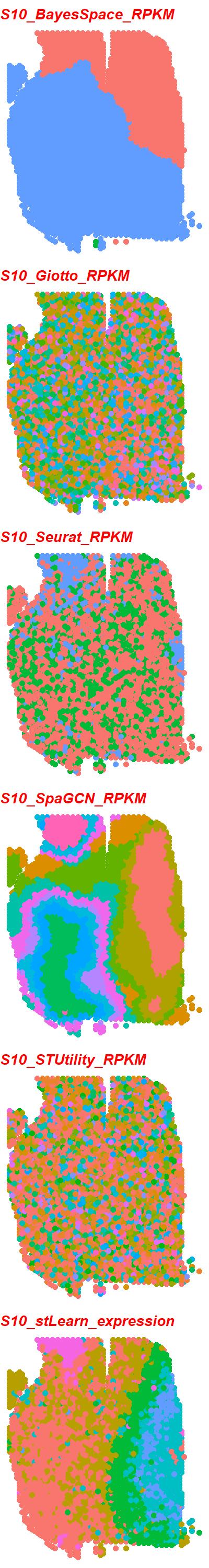

### S10_6_tools_scran.jpeg

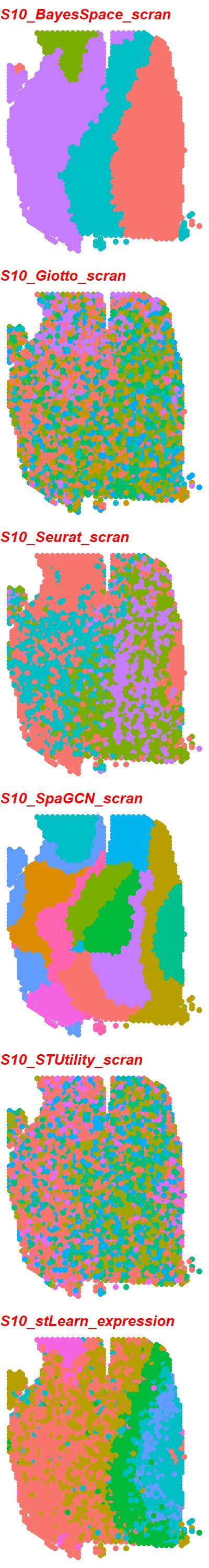

### S10_6_tools_scTransform.jpeg

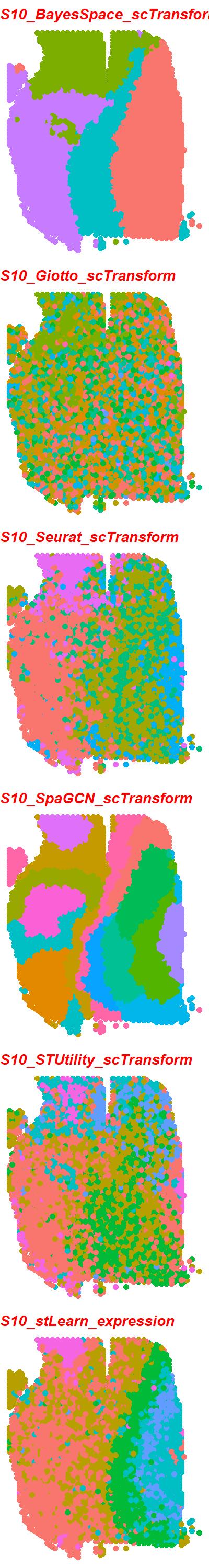

### S10_6_tools_TMM.jpeg

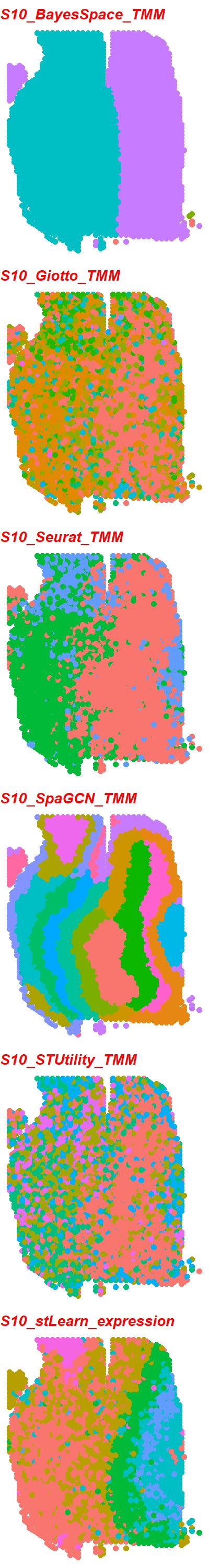

### S10_6_tools_TPM.jpeg

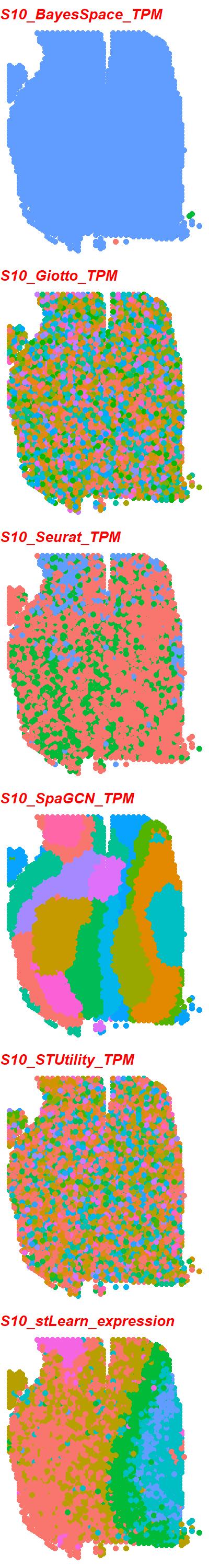

### S10_scGNN_DESeq2_PEalpha0.5_zdim32.png

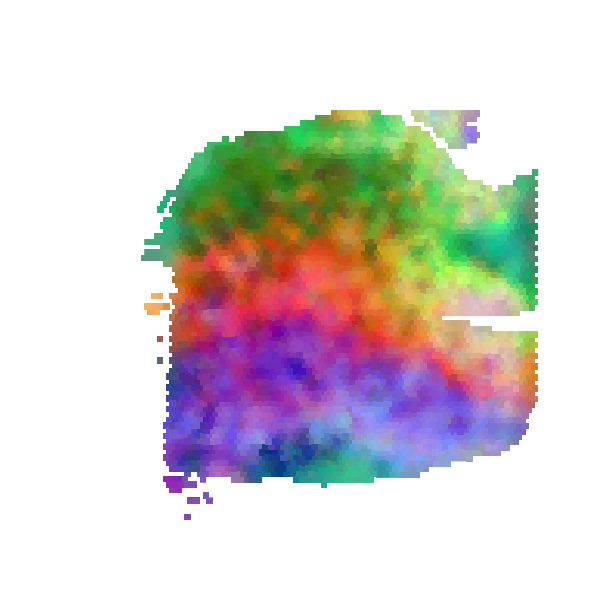

### S11_6_tools_RPKM.jpeg

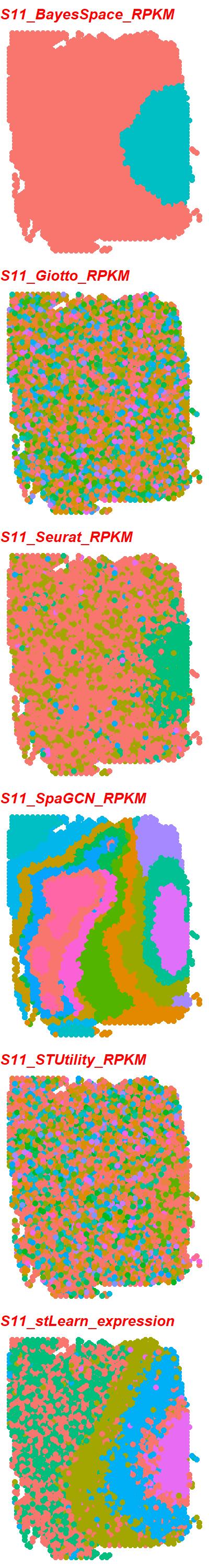

### S11_6_tools_scran.jpeg

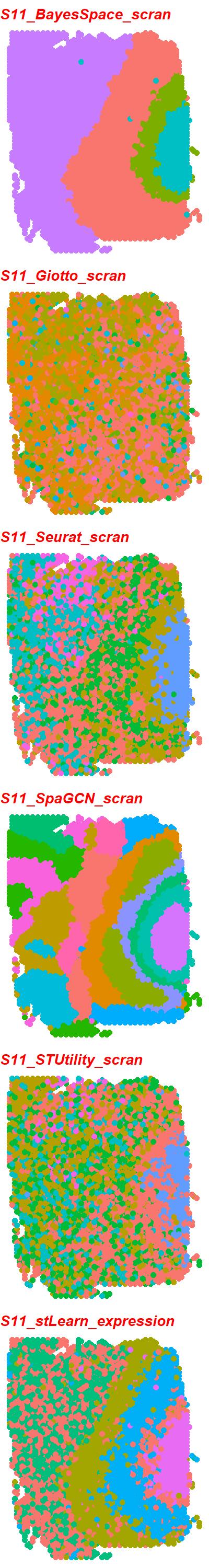

### S11_6_tools_scTransform.jpeg

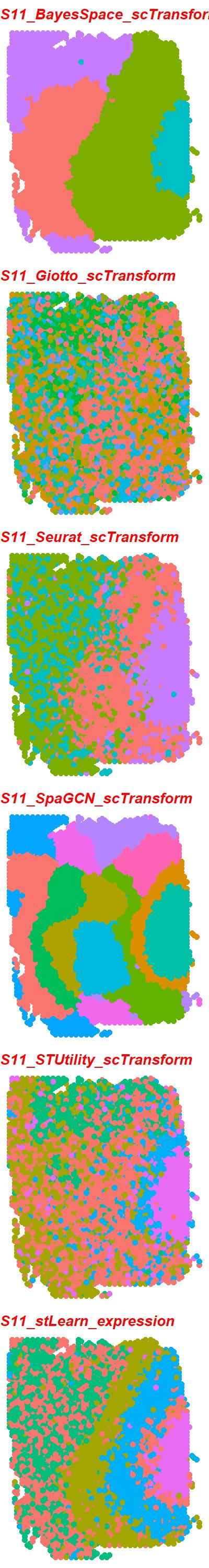

### S11_6_tools_TMM.jpeg

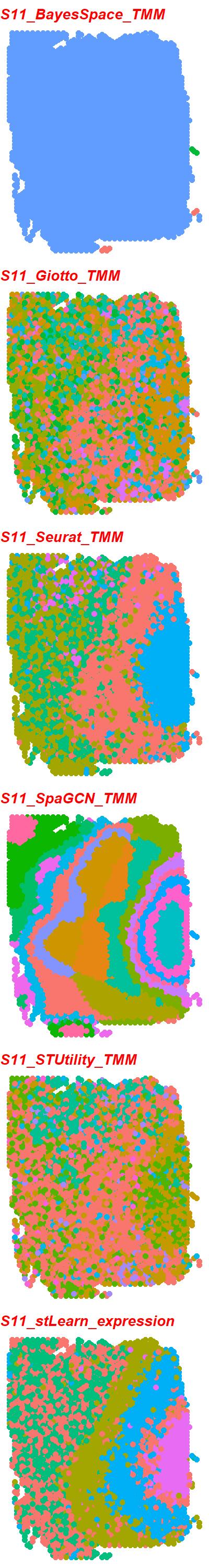

### S11_6_tools_TPM.jpeg

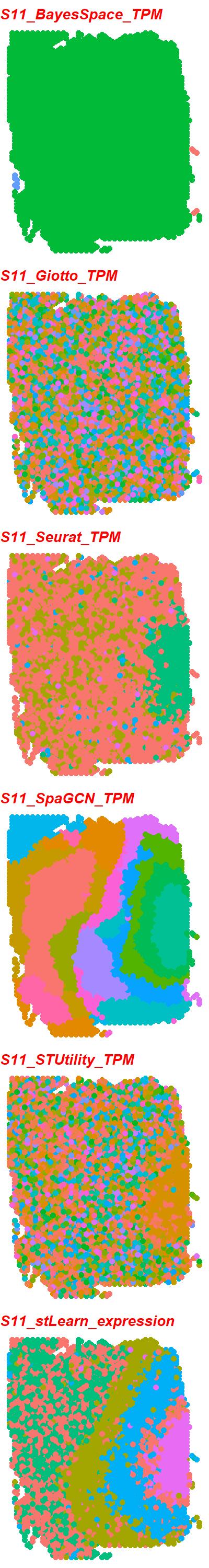

### S11_scGNN_DESeq2_PEalpha1.5_zdim256.png

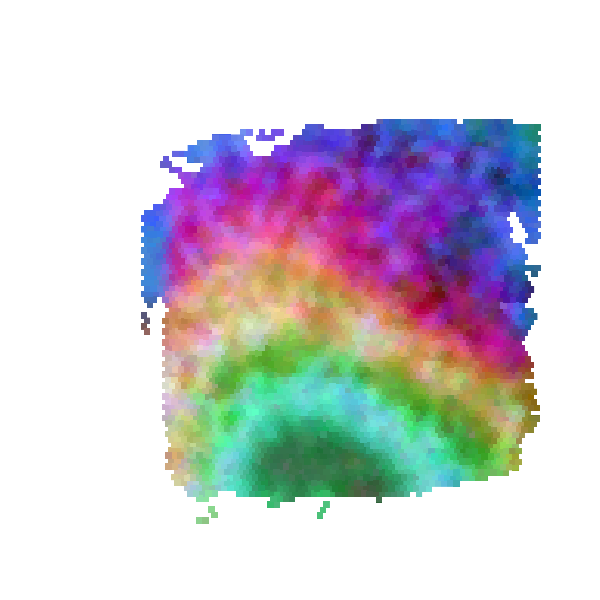

### S12_6_tools_RPKM.jpeg

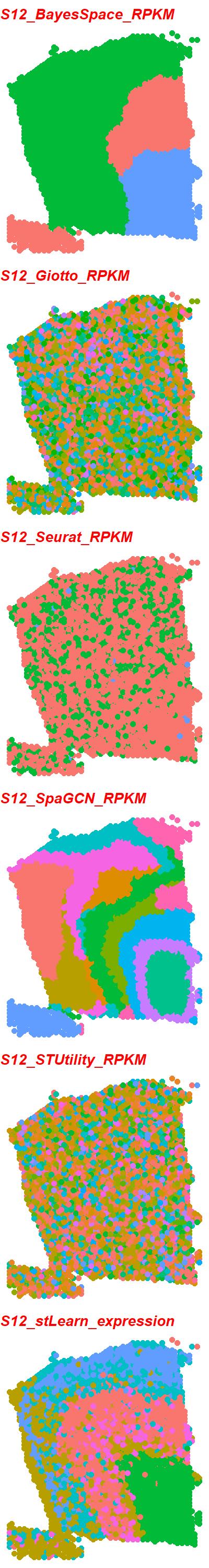

### S12_6_tools_scran.jpeg

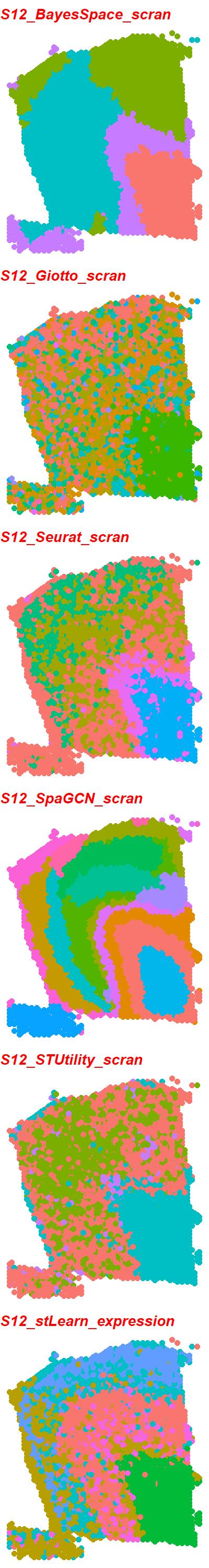

### S12_6_tools_scTransform.jpeg

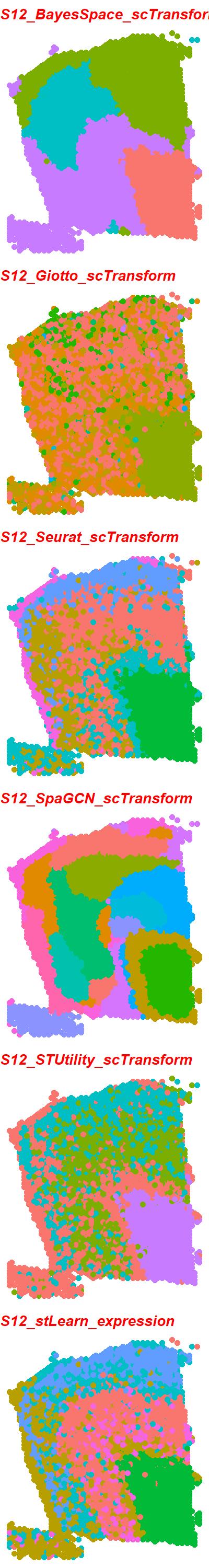

### S12_6_tools_TMM.jpeg

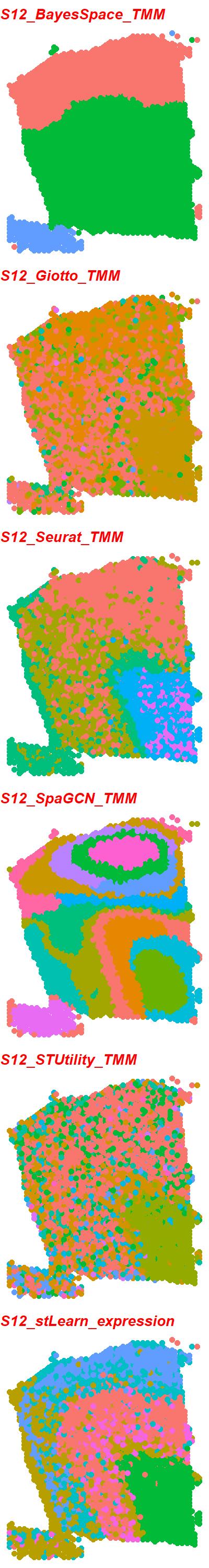

### S12_6_tools_TPM.jpeg

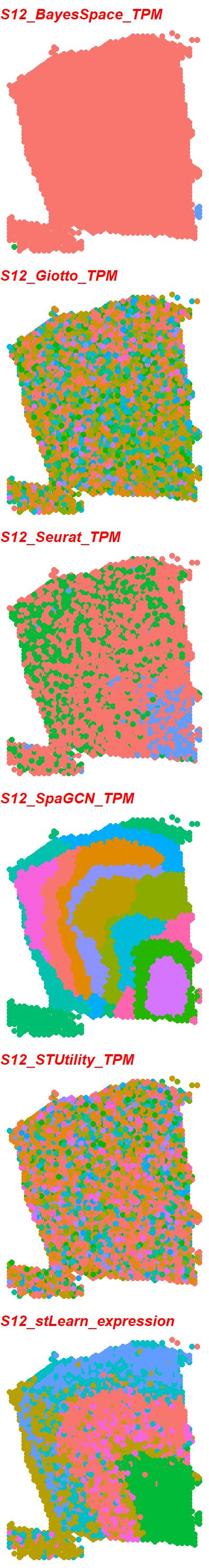

### S12_scGNN_DESeq2_PEalpha0.2_zdim128.png

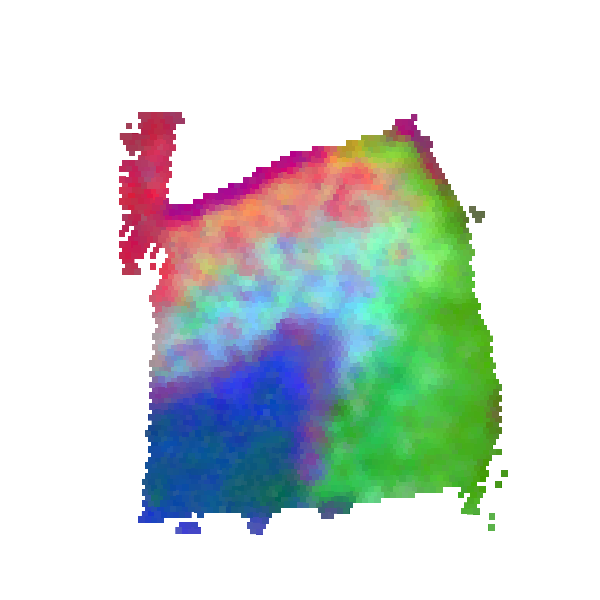

### S13_6_tools_RPKM.jpeg

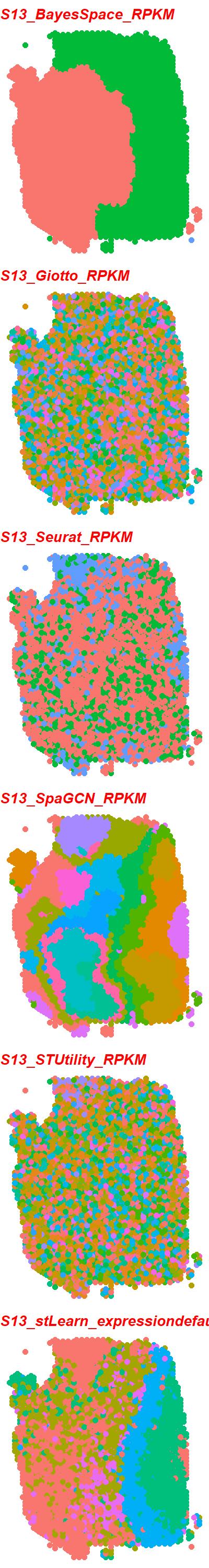

### S13_6_tools_scran.jpeg

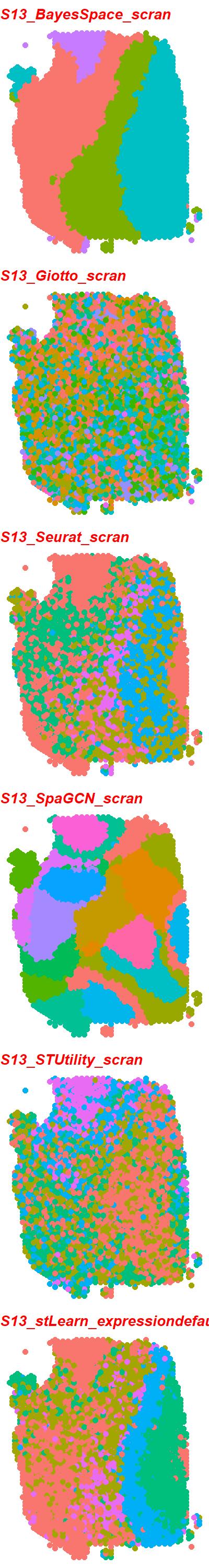

### S13_6_tools_scTransform.jpeg

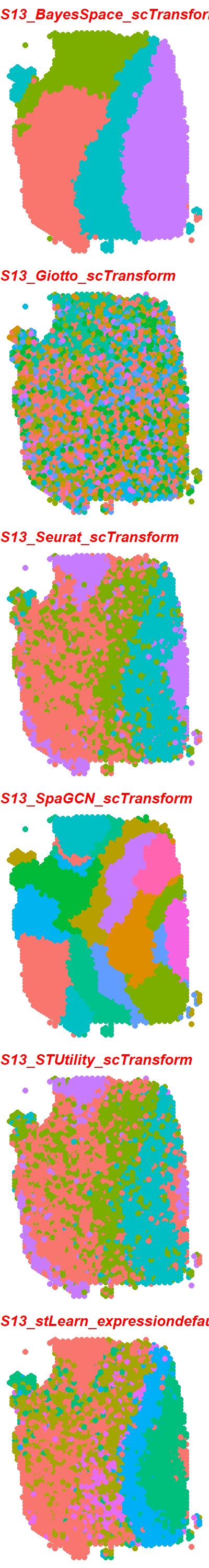

### S13_6_tools_TMM.jpeg

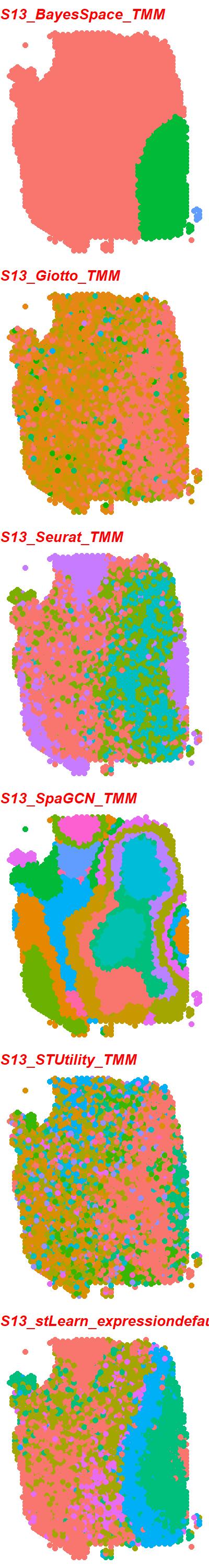

### S13_6_tools_TPM.jpeg

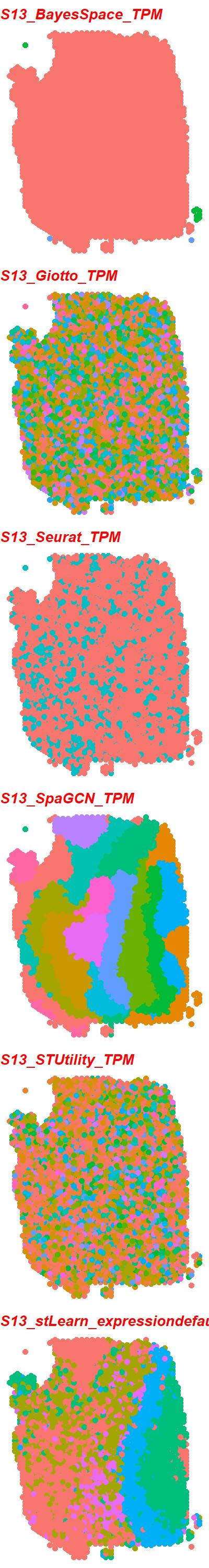

### S13_scGNN_DESeq2_PEalpha1.5_zdim10.png

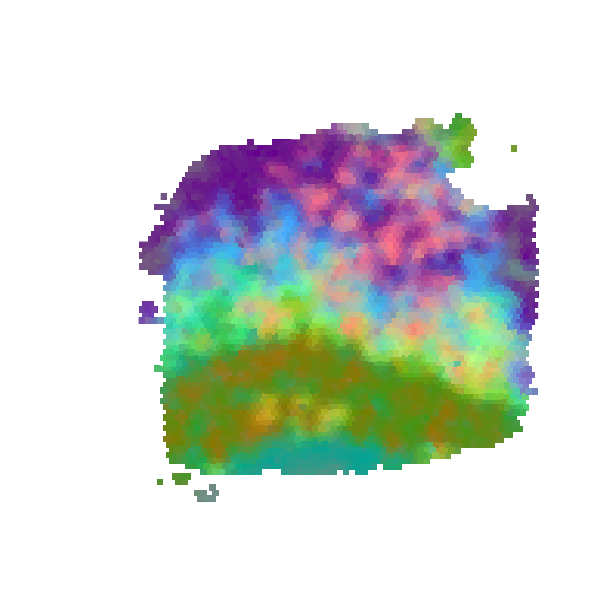

### S14_6_tools_RPKM.jpeg

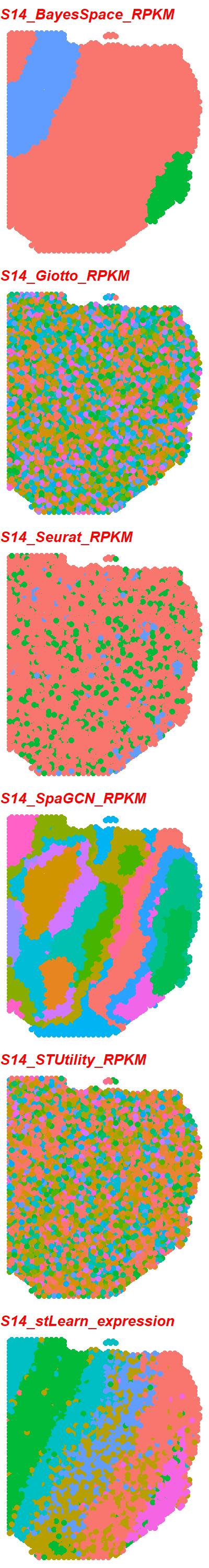

### S14_6_tools_scran.jpeg

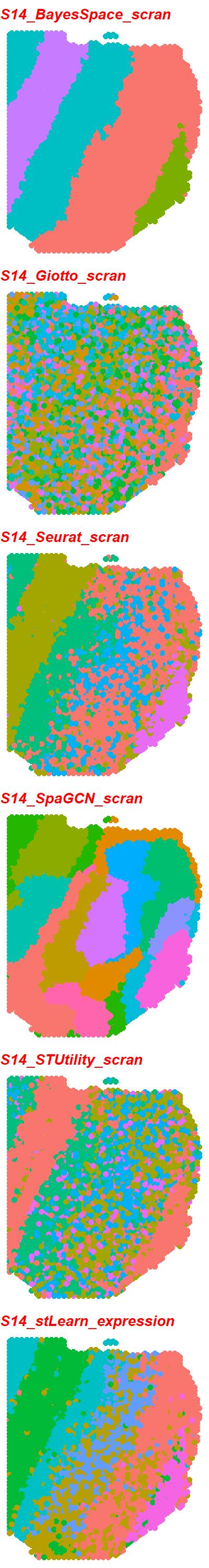

### S14_6_tools_scTransform.jpeg

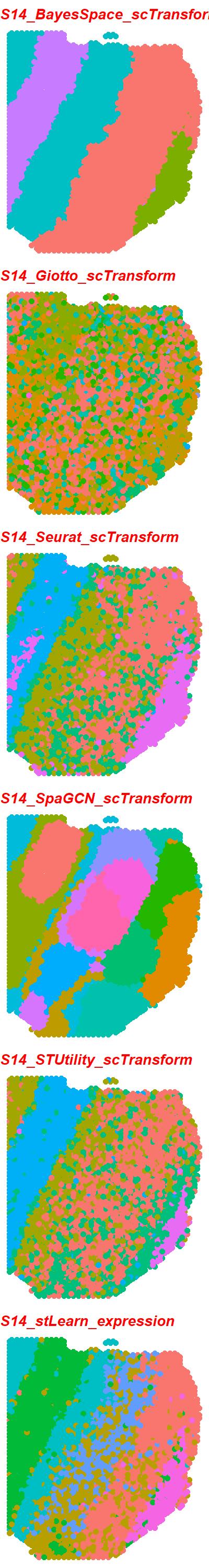

### S14_6_tools_TMM.jpeg

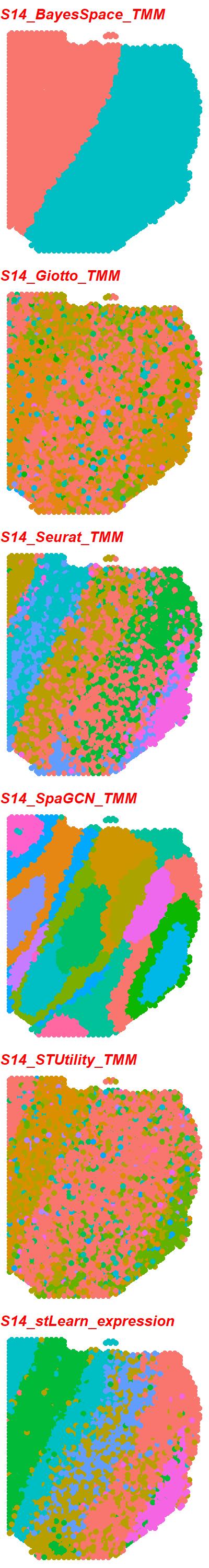

### S14_6_tools_TPM.jpeg

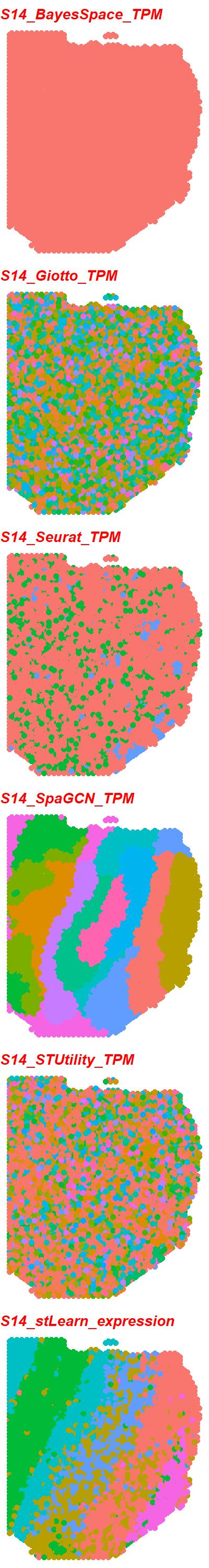

### S14_scGNN_DESeq2_PEalpha0.3_zdim128.png

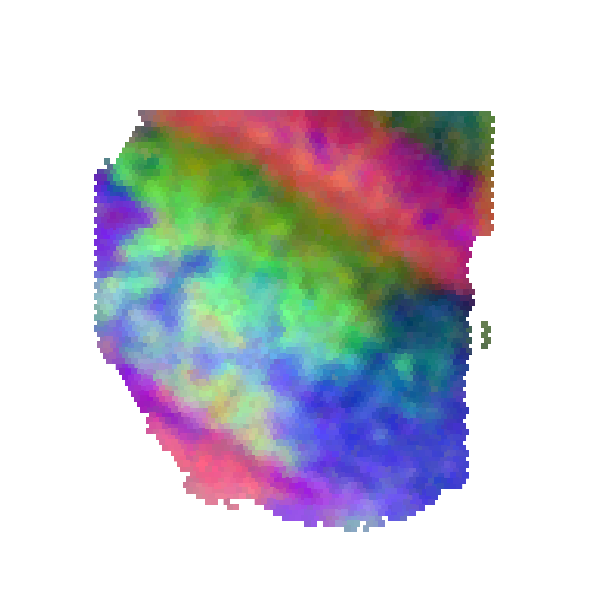

### S15_6_tools_RPKM.jpeg

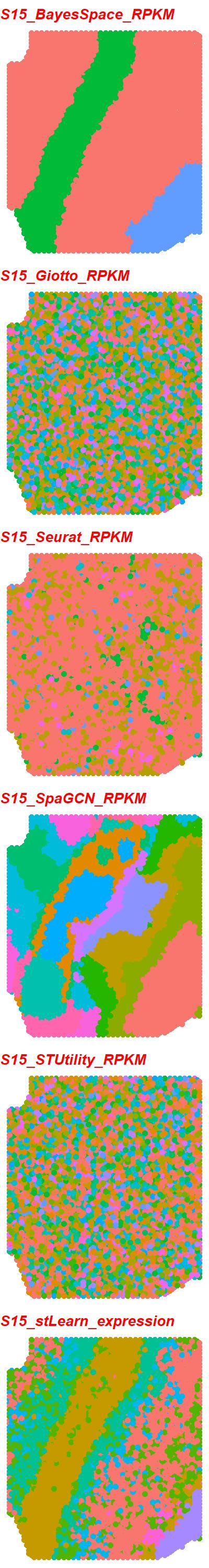

### S15_6_tools_scran.jpeg

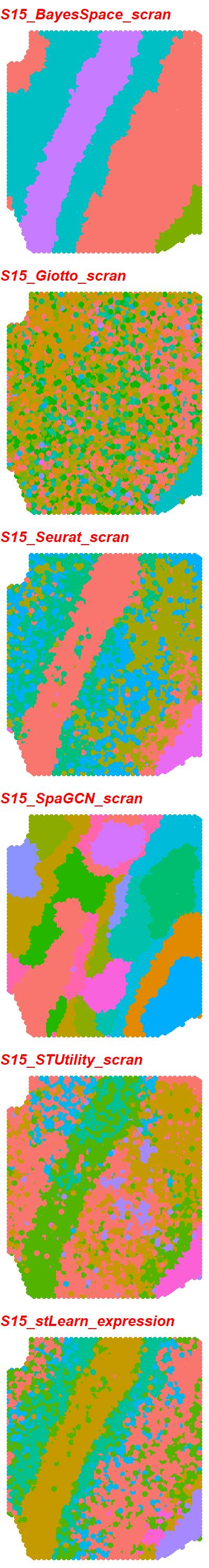

### S15_6_tools_scTransform.jpeg

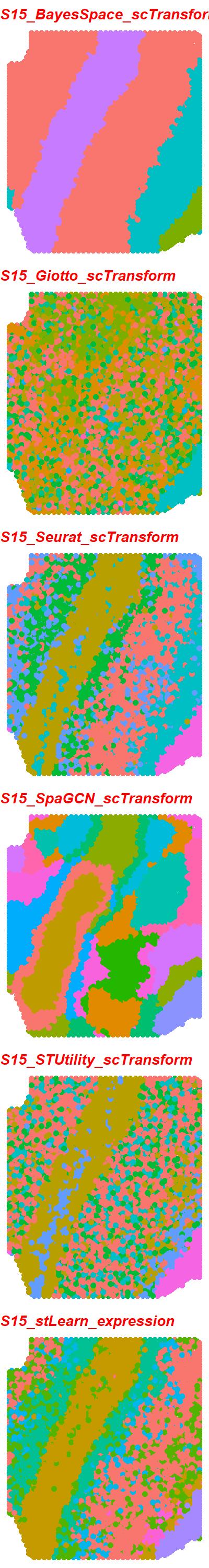

### S15_6_tools_TMM.jpeg

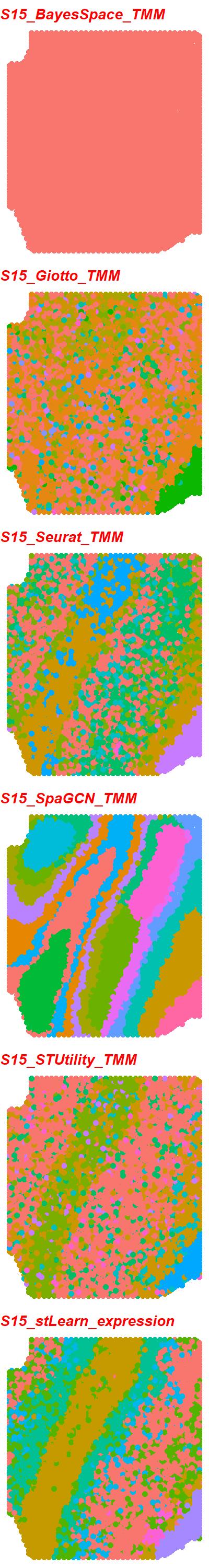

### S15_6_tools_TPM.jpeg

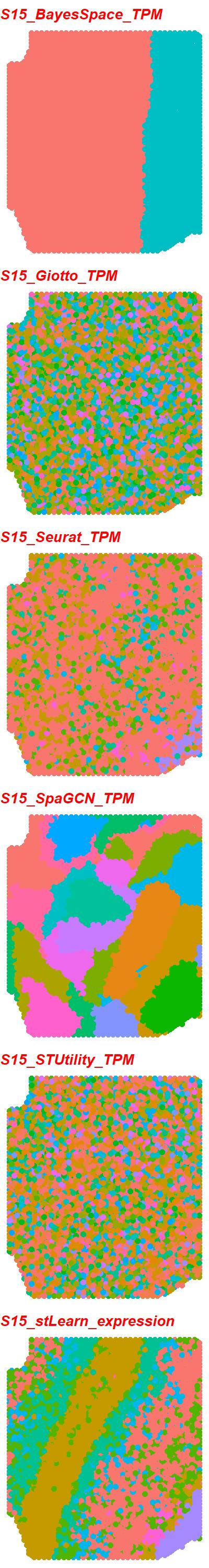

### S15_scGNN_DESeq2_PEalpha2.0_zdim32.png

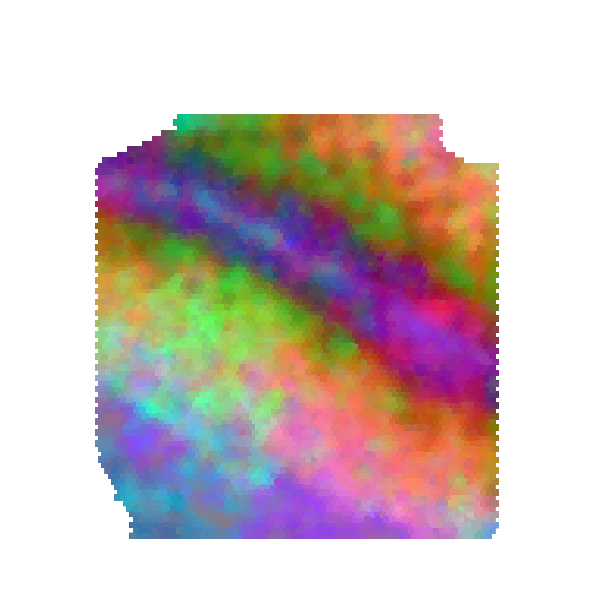

### S16_6_tools_RPKM.jpeg

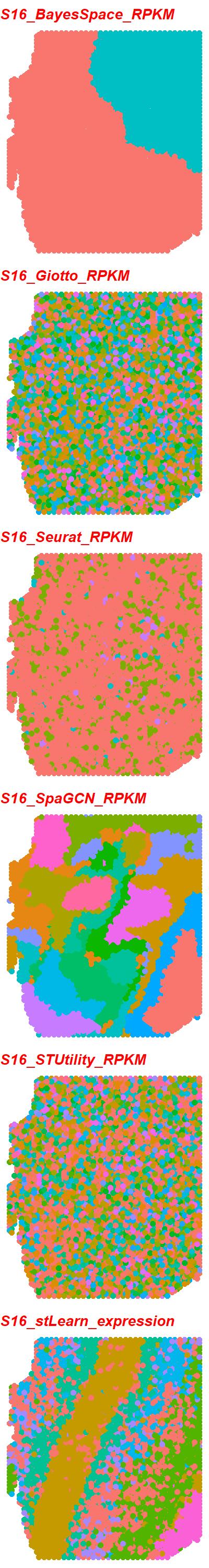

### S16_6_tools_scran.jpeg

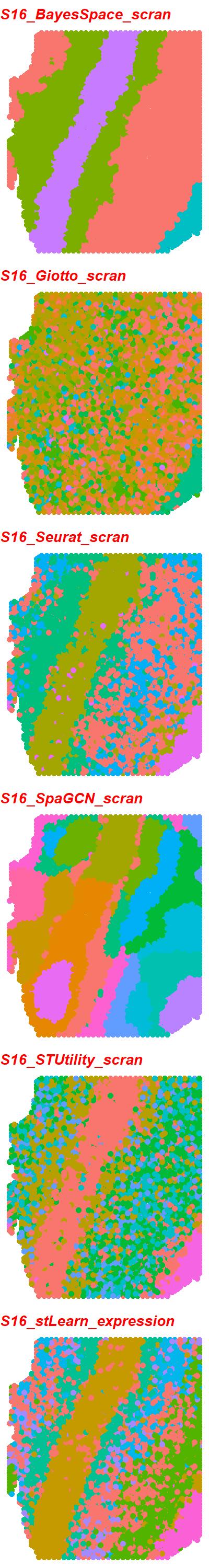

### S16_6_tools_scTransform.jpeg

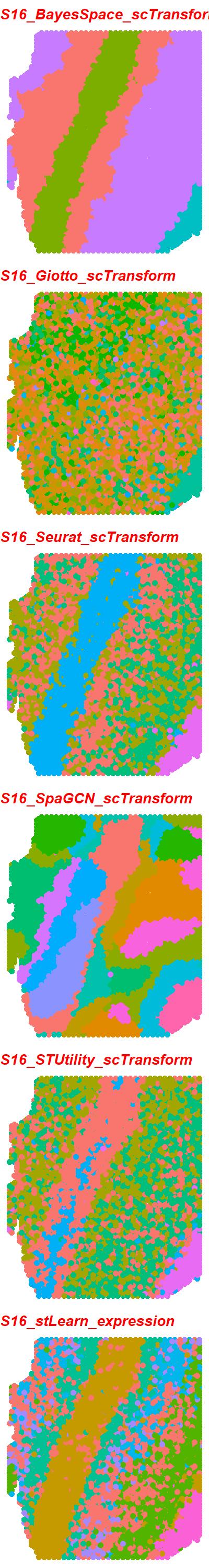

### S16_6_tools_TMM.jpeg

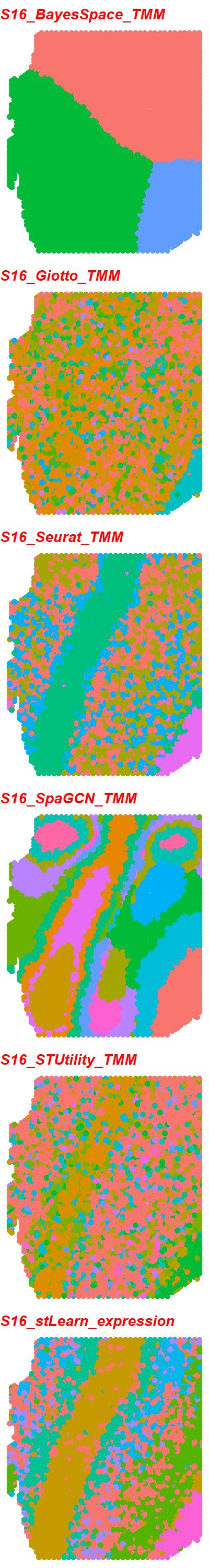

### S16_6_tools_TPM.jpeg

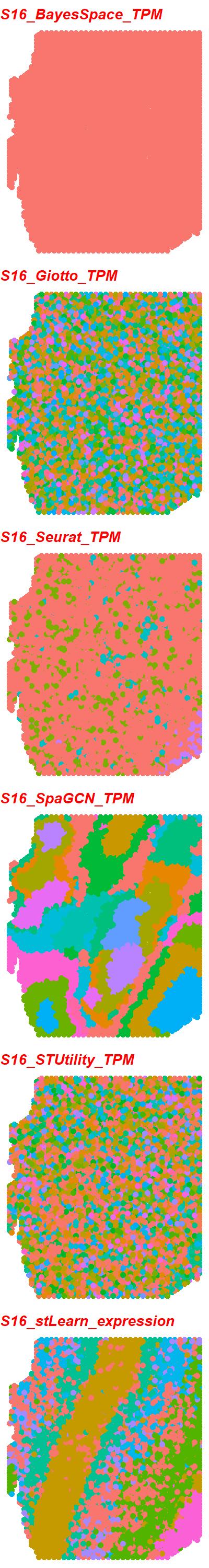

### S16_scGNN_DESeq2_PEalpha0.3_zdim32.png

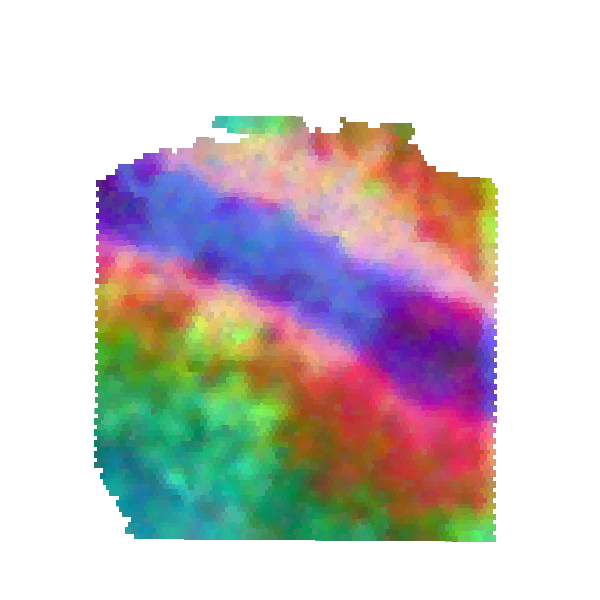

### S17_6_tools_RPKM.jpeg

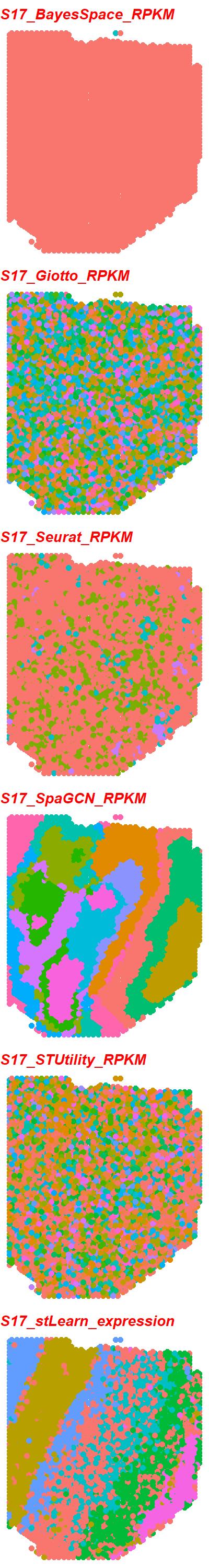

### S17_6_tools_scran.jpeg

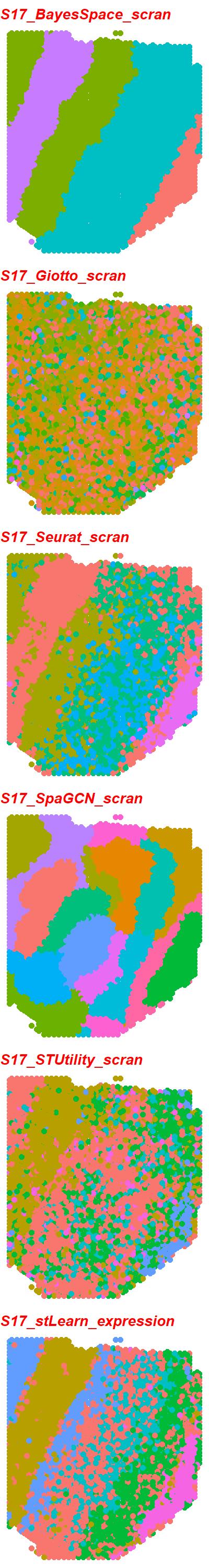

### S17_6_tools_scTransform.jpeg

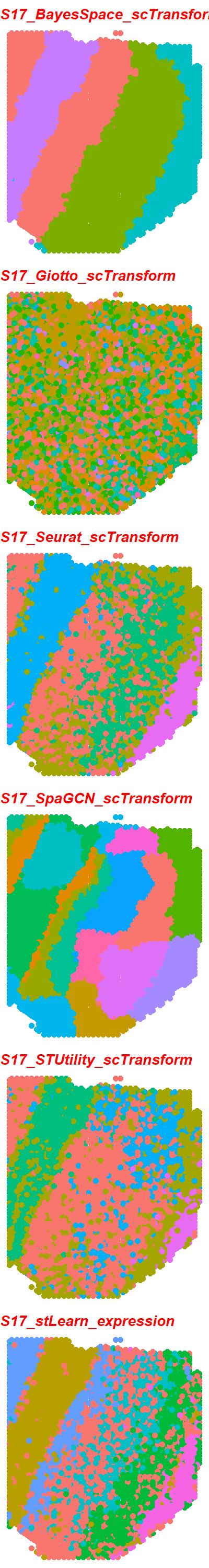

### S17_6_tools_TMM.jpeg

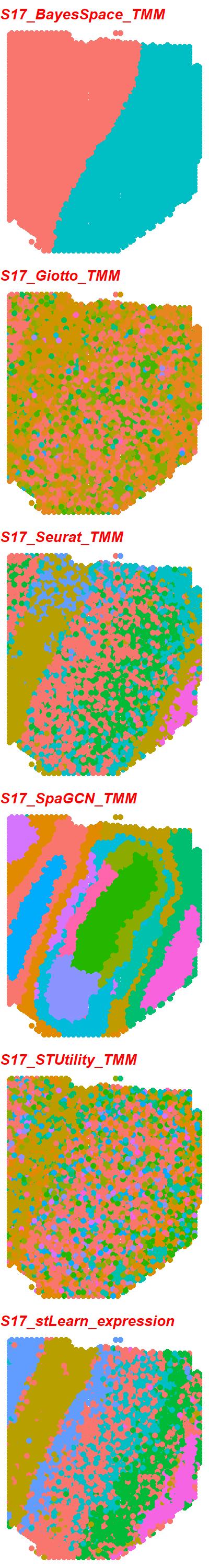

### S17_6_tools_TPM.jpeg

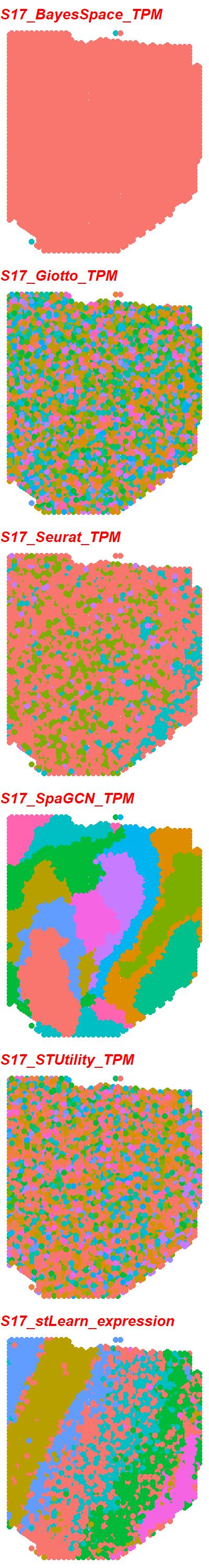

### S17_scGNN_DESeq2_PEalpha2.0_zdim128.png

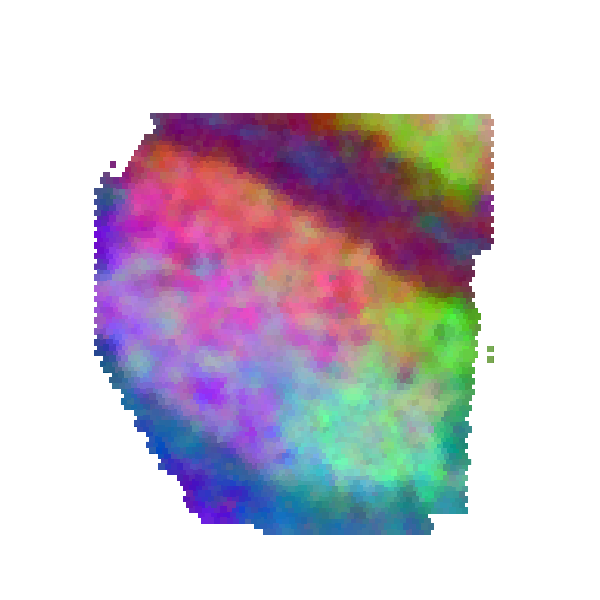
